# A reversible allosteric inhibitor of GlyT2 alleviates neuropathic pain without on-target side effects

**DOI:** 10.1101/2025.04.21.649698

**Authors:** Ryan P. Cantwell Chater, Julian Peiser-Oliver, Tanmay K. Pati, Ada S. Quinn, Irina Lotsaris, Zachary J. Frangos, Kristen E. Anderson, Anna E. Tischer, Billy J Williams-Noonan, Karin R. Aubrey, Megan L. O’Mara, Michael Michaelides, Sarasa A. Mohammadi, Christopher L. Cioffi, Robert J. Vandenberg, Azadeh Shahsavar

## Abstract

Chronic neuropathic pain, caused by nerve damage or disease, is increasing in prevalence, but current treatments are ineffective and over-reliant on opioids. The neuronal glycine transporter, GlyT2, regulates inhibitory glycinergic neurotransmission and represents a promising target for new analgesics. However, most GlyT2 inhibitors cause significant side effects, in part due to irreversible inhibition at analgesic doses. Here we develop a reversible inhibitor of GlyT2, RPI-GLYT2-82, and identify its binding site by determining cryo-EM structures of human GlyT2. We capture three fundamental conformational states of GlyT2 in the substrate-free state, and bound to either glycine, RPI-GLYT2-82 or the pseudo-irreversible inhibitor ORG25543. We demonstrate that RPI-GLYT2-82 dissociates from GlyT2 faster than ORG25543, providing analgesia in mouse neuropathic pain models without on-target side-effects or addiction liability. Our data provide a mechanistic understanding of allosteric inhibition of glycine transport, enabling structure-based design of non-opioid analgesics.

## Introduction

Chronic neuropathic pain affects 7-10% of the population yet only one in four of these patients respond to the gold standard treatment, pregabalin. Furthermore, because the efficacy of pregabalin is limited and accompanied by dose-dependent side effects, there is an over-reliance on opioid medications, exacerbating opioid use disorder and its associated mortality^1,2^. The rising incidence of neuropathic pain has therefore created an urgent need for the development of safe and effective analgesics that circumvent the opioid mechanism by targeting new pathways^2^.

A key feature of neuropathic pain is a reduction of glycinergic neurotransmission in the dorsal horn of the spinal cord – an adaptation that impairs our capacity to distinguish between painful and harmless stimuli^3^. Glycine is an inhibitory neurotransmitter that hyperpolarises postsynaptic neurons by binding to and opening Cl^−^-permeable glycine receptors. In the central nervous system, the concentration of extracellular glycine is tightly regulated by glycine transporters 1 and 2 (GlyT1 and GlyT2). GlyT1 is expressed at both excitatory and inhibitory synapses, whereas GlyT2 is exclusively localized to the presynaptic neurons of inhibitory synapses, primarily in the spinal cord^4^. GlyT2 thus plays a key role in regulating glycinergic neurotransmission by controlling glycine receptor activity on postsynaptic neurons and by recycling glycine for subsequent release from presynaptic neurons^5–8^. GlyT2 (encoded by *SLC6A5*) belongs to the solute carrier 6 (SLC6) subfamily of secondary active neurotransmitter/sodium symporters (NSSs) ^9,10^. NSSs are membrane proteins that operate via an alternating access mechanism, whereby transmembrane helices undergo conformational rearrangements during the transport cycle that alternately expose the substrate-binding site to each side of the membrane. In eukaryotic NSSs, binding of extracellular substrate and initiation of the transport cycle is dependent on a Na^+^ and Cl^−^ gradient, such that GlyT2 co-transports three Na^+^ ions and one Cl^−^ ion with each glycine^11–13^. Upon ion and substrate binding, access to the extracellular environment is occluded as the transporter transitions from an outward-facing to an inward-facing conformation, followed by release of solutes.

Several competitive and non-competitive SLC6 inhibitors have been developed that target either outward or inward-facing conformations of these transporters^14–25^. At glycinergic synapses, competitive GlyT2 inhibitors are likely to be ineffective due to the high concentrations of glycine in the cleft during synaptic transmission, whereas non-competitive inhibitors that target allosteric sites will be more effective^8^. Allosteric inhibitors are also more likely to result in subtype selectivity as substrate binding sites show little amino acid divergence across the NSS superfamily. Furthermore, because glycine is the only known substrate transported by GlyT2, the chemical space that can be explored for orthosteric inhibition is restricted^26^. Interestingly, ORG25543 (4-(benzyloxy)-N-{[1-(dimethylamino)cyclopentyl]methyl}-3,5-dimethoxybenzamide) is a potent, pseudo-irreversible, and non-competitive^27^ inhibitor of GlyT2 that provides analgesia in animal models of neuropathic pain^28–31^, without affecting acute or inflammatory pain thresholds^28^. In rat spinal cord slices, ORG25543 enhances tonic glycinergic neurotransmission and prolongs the decay time course of both miniature and evoked glycinergic inhibitory postsynaptic currents, suggesting that its effects are likely due to elevated extracellular glycine concentrations^32^. However, the compound did not progress to clinical use due to its narrow therapeutic index and significant on-target side effects, including tremors, seizures, and mortality at the maximal analgesic dose. These effects, which resemble the phenotype of the mouse *GlyT2^-/-^* knockout, are likely due to the high affinity and slow dissociation rate of ORG25543^28,29^. Indeed, siRNA knockdown of GlyT2 to 30% of wild-type (WT) levels results in prolonged analgesia without confounding side effects in mouse models of neuropathic pain^33^. In addition, the endogenous bioactive lipid N-arachidonoyl glycine and synthetic derivatives such as oleoyl-D-lysine cause partial inhibition of GlyT2 comparable to knockdown studies and provide analgesia without side effects. However, the poor pharmacokinetics of these bioactive lipids prevented progression to clinical trials^28,34,35^. Recently, an ORG25543-based benzamide, opiranserin (VVZ-149), reached a confirmatory phase 3 trial for the treatment of post-colectomy pain^36,37^, validating the importance of GlyT2 as a target for new pain therapeutics. Opiranserin, however, only shows moderately potent inhibitory activity on GlyT2^38^.

Here we synthesize a potency-tuned analogue of ORG25543, RPI-GLYT2-82 (1-benzoyl-N-((4-(dimethylamino)tetrahydro-2H-pyran-4-yl)methyl)indoline-5-carboxamide) and determine its suitability for development as a treatment for neuropathic pain. We determine the first structures of human GlyT2 (hGlyT2) – without substrate (inward-open conformation) and bound to either glycine (inward-occluded state), ORG25543 (outward-open state), or RPI-GLYT2-82 (outward-open state) – and confirm the allosteric action of both inhibitors. We further show that RPI-GLYT2-82 is efficacious at alleviating allodynia in both chronic constriction injury (CCI) and partial sciatic nerve ligation (PSNL) mouse models, without any side effects at the maximal analgesic dose. Our study thus reveals the molecular mechanism of allosteric inhibition of glycine transport, providing a framework by which new analgesics can be developed.

## Results

### RPI-GLYT2-82 is a non-competitive GlyT2 inhibitor

The narrow therapeutic index of ORG25543 has been attributed to its slow dissociation rate, suggesting that ORG25543-based compounds with improved reversibility may decrease on-target side effects and toxicity^29,39,40^. To identify such an allosteric inhibitor, we pursued structural modifications of the ORG25543 scaffold and evaluated the incorporation of ring-constrained systems, such as indoles and indolines, as viable substituents. These efforts culminated in the development of a potency-tuned analogue of ORG25543, RPI-GLYT-82, for further assessment (Supplementary methods).

We first characterized the inhibitory action of RPI-GLYT2-82 by expressing WT hGlyT2 (hGlyT2^WT^) in *Xenopus* oocytes and using the two-electrode voltage clamp technique to record glycine-induced transport currents in the presence of different concentrations of the compound. RPI-GLYT2-82 behaved as a non-competitive inhibitor (Fig. 1a, Supplementary Table 1) with a half-maximal inhibitory concentration (IC_50_) of 554 nM (Fig. 1b). This confirmed the lower inhibition potency of RPI-GLYT2-82 for hGlyT2^WT^ than ORG25543, which displayed an IC_50_ of 11.4 nM (Fig. 1c). Glycine-induced transport currents recovered to pre-inhibition levels within five minutes of RPI-GLYT2-82 removal, indicating a markedly faster transport recovery rate than ORG25543, which required 30 minutes for currents to be restored to only 12.5% of pre-inhibitor levels (Fig 1d, e, Supplementary Fig. 1). Like ORG25543, RPI-GLYT2-82 is selective for GlyT2 over GlyT1 (Fig. 1f). Together, these data show that RPI-GLYT-82 is a non-competitive GlyT2 inhibitor that binds with reduced potency compared to ORG25543. Interestingly, while the inhibition of transport by both compounds appears to be Na⁺-independent, RPI-GLYT2-82 exhibits slightly increased potency under lower Na⁺ concentrations, whereas inhibition by ORG25543 remains unaffected (Fig. 1g,h).

**Fig. 1.**
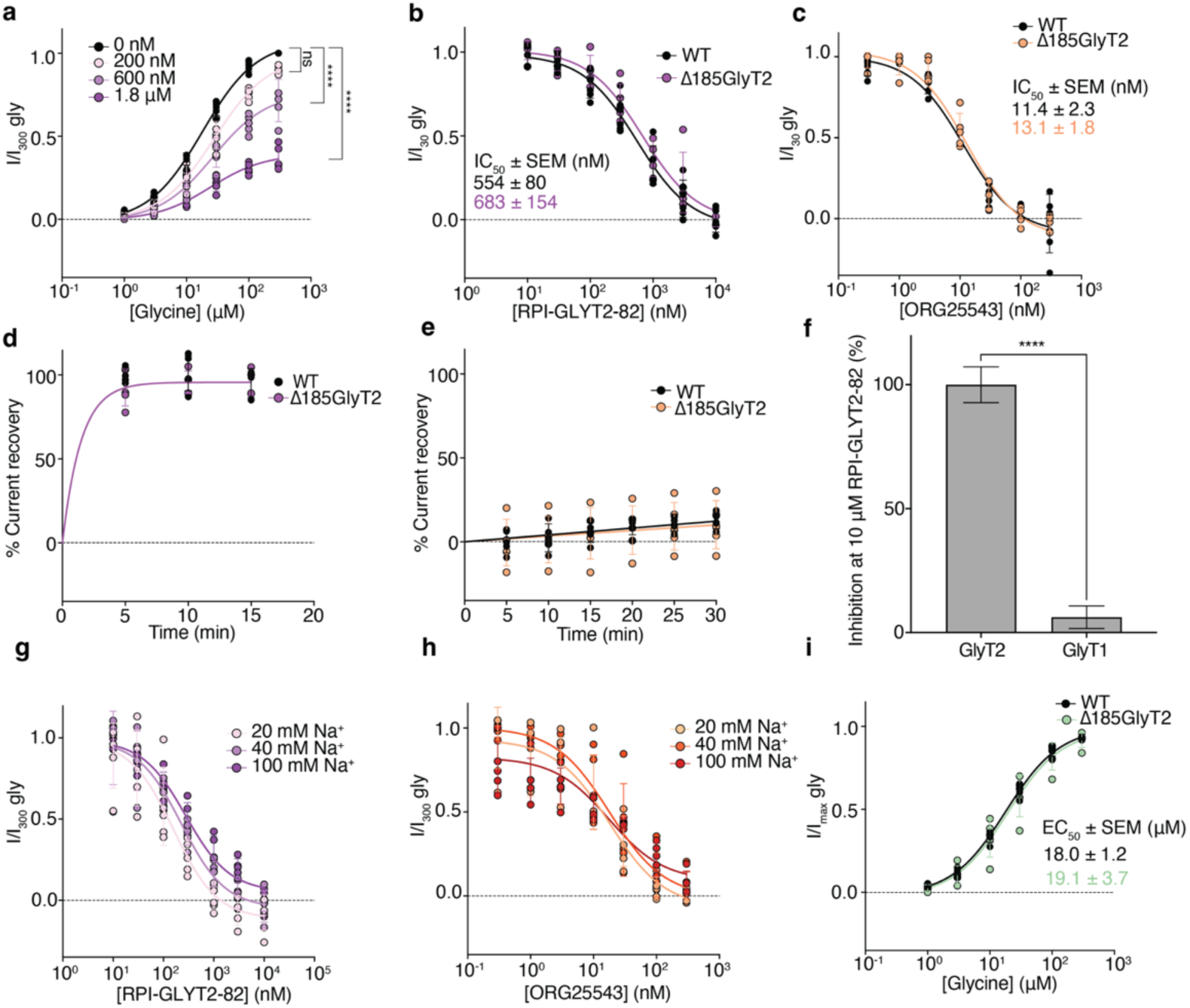
RPI-GLYT2-82 is a non-competitive inhibitor of hGlyT2. **a** Glycine concentration-dependent currents in the presence of different concentrations of RPI-GLYT2-82 showing non-competitive inhibition. There was no statistically significant difference in the glycine EC_50_ values across all four conditions (p = 0.49) but there was a statistically significant reduction in I_max_ between 0 nM RPI-GLYT2-82 (mean I_max_ ± SEM = 1.07 ± 0.01) and the I_max_ at 600 nM RPI-GLYT2-82 (mean I_max_ ± SEM = 0.76 ± 0.04) (p < 0.0001) and also the I_max_ at 1.8 µM RPI-GLYT2-82 (mean I_max_ ± SEM = 0.39 ± 0.02) (p < 0.0001). Data analysed using nonlinear regression (see Methods). **b** RPI-GLYT2-82 concentration-dependent inhibition of glycine transport currents. Current-responses were normalised to currents generated by 30 µM glycine. The difference between the hGlyT2^WT^ IC_50_ = 554 nM (95% CI: 412 to 747 nM) and the hGlyT2^Δ185^ IC_50_ = 683 nM (95% CI: 370 to 996 nM) is not statistically significant (p = 0.45) **c** ORG25543 concentration-dependent inhibition of glycine transport normalised to currents generated by 30 µM glycine. The difference between the hGlyT2^WT^ IC_50_ = 11.4 nM (95% CI: 8.0 to 16.4 nM) and the hGlyT2^Δ185^ IC_50_ = 13.1 nM (95% CI: 9.4 to 16.8 nM) is not statistically significantly (p = 0.60). **d** RPI-GLYT2-82 is reversible and washes out within 5-minutes. Data fitted to a one-phase decay curve (WT not fitted due to full reversal before the first time point). **e** ORG25543 has prolonged inhibitory effects with minimal restoration of glycine-induced currents 30-minutes post-ORG25543. Results normalised to currents generated by 30 µM glycine. **f** RPI-GLYT2-82 inhibits GlyT2 with minimal effect on GlyT1 showing only 6% inhibition at 10 μM (p < 0.0001). Mean inhibition ± SEM from 10 µM RPI-GLYT2-82 is plotted. **g** RPI-GLYT2-82 is more potent at lower Na^+^ concentrations. The IC_50_ at 20 mM Na^+^ (IC_50_ = 159 nM (95% CI: 81 to 238 nM)) is significantly lower than that at 100 mM Na^+^ (IC_50_ = 279 nM (95% CI: 182 to 376 nM)) (p = 0.04). **h** ORG25543 potency does not change with decreasing Na^+^ concentration. The IC_50_ at 20 mM Na^+^ (IC_50_ = 18 nM (95% CI: 6.8 to 29 nM)) is not significantly different than that at 100 mM Na^+^ (IC_50_ = 17.6 nM (95% CI: 5.5 to 30 nM)) (p > 0.9999). **i** Normalised glycine-dependent transport currents in hGlyT2^WT^- and hGlyT2^Δ185^-expressing *Xenopus laevis* oocytes showing comparable transport activity in the two constructs (p = 0.78). EC_50_, IC_50_, washout and competition experiments performed with *n* = 5 cells, from at least two different batches of oocytes. Error bars represent SD.

### hGlyT2 adopts a LeuT fold

To understand the mechanism by which ORG25543 and RPI-GLYT-82 inhibit GlyT2, we sought to determine the structures of hGlyT2 in the absence and presence of these inhibitors and the substrate glycine. Because the N-terminus of hGlyT2 is predicted to be largely unstructured and its deletion does not affect trafficking to the plasma membrane or glycine transport^41^, we created a construct lacking residues 2-185 (hGlyT2^Δ185^) to facilitate expression and purification. hGlyT2^Δ185^ exhibited a similar half-maximal effective concentration (EC_50_) for glycine transport, and similar IC_50_ values for both ORG25543 and RPI-GLYT2-82, to full-length hGlyT2^WT^ (Fig. 1b, c, i).

We purified hGlyT2^Δ185^ in mixed micelles containing lauryl maltose neopentyl glycol (LMNG), glyco-diosgenin (GDN), and cholesteryl hemisuccinate (CHS) at a ratio of 1.0:1.0:0.1 to generate monodisperse and homogenous hGlyT2^Δ185^ (Supplementary Fig. 2) for cryo-electron microscopy (cryo-EM) studies. This enabled structures of hGlyT2^Δ185^ in the absence of substrate and bound to glycine, ORG25543, or RPI-GLYT2-82 to be determined at 2.97 Å, 3.02 Å, 2.49 Å, and 2.79 Å resolution, respectively (Fig. 2a–f, Supplementary Fig. 3-6, Supplementary Table 2). The high-quality of the resulting density maps enabled unambiguous modelling of all 12 transmembrane (TM) helices, revealing unwinding of TMs 1 and 6 approximately halfway across the membrane – a characteristic feature of the substrate binding site in SLC6 transporters (Fig. 2, Supplementary Fig. 7-10). We were also able to model the intracellular and extracellular loops, except parts of extracellular loop 2 (EL2; residues 302 – 393) due to its high degree of flexibility (Fig. 2, Supplementary Fig. 7-10), as well as the position of bound ligands and ions. The conserved disulfide bridge between C311 and C320 in EL2 is present in the inhibitor-bound and substrate-free structures (Supplementary Fig. 7, 9-11). However, although we observed clear densities for C311 and C320 in glycine-bound hGlyT2^Δ185^, we did not detect a distinct density for the disulfide bridge, suggesting that the cysteine residues may be reduced in the substrate-bound state (Supplementary Fig. 8). The overall architecture adopted by hGlyT2 is characteristic of SLC6 transporters: an inverted pseudo-twofold symmetric organisation of TMs 1–5 relative to TMs 6–10 (Fig. 2, Supplementary Fig. 7-10), known as the LeuT fold due to its identification in the bacterial amino acid importer LeuT^42^.

**Fig. 2.**
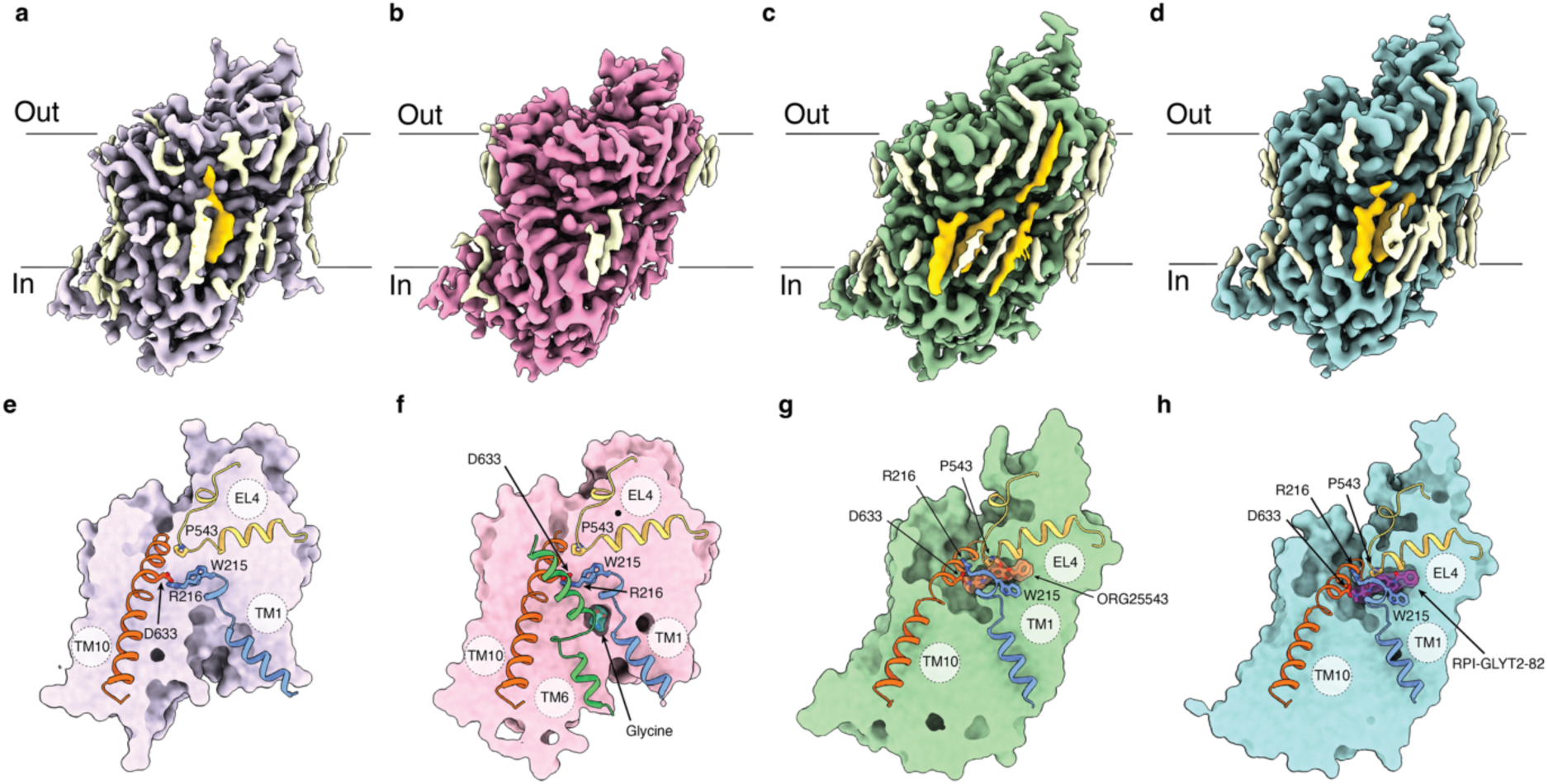
Substrate-free, glycine-bound, and inhibitor-bound states of hGlyT2. Cryo-EM density maps of hGlyT2^Δ185^ in **a** substrate-free state (purple map contour level = 0.04 in ChimeraX), bound to **b** substrate glycine (pink map contour level = 0.046), **c** ORG25543 (green map contour level = 0.055 in ChimeraX), and **d** RPI-GLYT2-82 (blue map contour level = 0.038 in ChimeraX). Lipid densities shown in light yellow, densities corresponding to sterols built into the models shown in dark yellow. **e-h** Surface representation of substrate-free, substrate-bound, and inhibitor-bound structures of hGlyT2^Δ185^ viewed parallel to the membrane. **e** Slice view of hGlyT2^Δ185^ in substrate-free form. Closed extracellular vestibule around W215 (blue) and open intracellular pathway are displayed. **f** Slice view of hGlyT2^Δ185^ showing glycine binding pocket (cyan, compound density shown at map contour level = 0.046 in ChimeraX) **g** Slice view of hGlyT2^Δ185^ showing ORG25543 binding pocket (orange, compound density shown at map contour level = 0.055 in ChimeraX). **h** Slice view of hGlyT2^Δ185^ showing RPI-GLYT2-82 binding pocket (dark purple, compound density shown at map contour level = 0.038 in ChimeraX). Residues W215 (TM1, blue), R216 (TM1, blue), P543 (EL4, yellow), and D633 (TM10, orange) are shown as sticks in **e-h**; TM6 shown in green in **g**.

GlyT2 and related SLC6 transporters are modulated by cholesterol^31,43–46^. Accordingly, we observed several cholesterol-like features in our hGlyT2^Δ185^ density maps, which were of sufficient resolution to be modelled in our substrate-free and inhibitor-bound structures (Supplementary Fig. 12). In these three structures, we observed a density consistent with a cholesterol (CHOL) molecule in an inner leaflet pocket formed by TM2, TM7, and TM11, equivalent to the previously reported CHOL2 site^17,20,21^. We identified an additional neighbouring density where we modelled a CHS molecule (CHOL5). These two molecules form hydrogen bonds with N674 (IL5). Interestingly, while the cholesterol at the CHOL2 site appears to be conserved, this asparagine residue is not conserved among NSS family members. In the *Drosophila* dopamine transporter (dDAT) and human serotonin transporter (hSERT) it is a glycine residue, in human DAT (hDAT) a serine, and in hGlyT1 and LeuT a proline (Supplementary Fig. 10, 11). We modelled a second CHS molecule adjacent to, but distinct from, the CHOL1 site (CHOL6) ^19,21^. This CHOL6 site lies parallel to and interacts with TM7. Above CHOL6, we modelled a cholesterol molecule in a novel site (CHOL7), formed by TM5, TM7, and EL4 (Supplementary Fig. 12). Two lipid densities were observed in CHOL6 and CHOL7 sites in a recent hDAT structure^20^. We further modelled a cholesterol molecule into a site that partially overlaps with CHOL3, and CHOL5 site proposed in molecular dynamics simulations of dDAT (named CHOL3’ in GlyT2)^17,18,47^. This cholesterol is embedded within the protein, at a methionine-rich site formed by TMs 3, 9, 10 and 12 (Supplementary Fig. 12), in contrast to its alignment with the membrane in other SLC6 structures.

We further examined the stability of sterols in the inhibitor-bound GlyT2 structures and found that most remained stably bound throughout the simulations (Supplementary Fig. 12d-l). However, the cholesterol molecule at the CHOL3’ site in the RPI-GLYT2-82-bound structure sampled multiple conformations while maintaining contact with the site, suggesting local flexibility in this region. Because this feature, along with other sterol molecules we modelled, is consistently present in all hGlyT2^Δ185^ structures in three different conformational states, we suggest that these sterols act primarily as important structural stabilisers of hGlyT2 rather than as regulatory elements. We did not observe any clear cholesterol-like densities at CHOL1 or CHOL4 sites^17^.

### Substrate-free hGlyT2 is in an inward-open state

Despite inclusion of the GlyT2 inhibitor oleoyl-D-lysine throughout purification, we did not observe any density corresponding to the inhibitor. The low solubility of the compound likely prevented it from remaining in sufficient concentration in solution to bind effectively. We therefore present this structure as the substrate-free state of hGlyT2^Δ185^ (Fig. 2, Supplementary Fig. 3, 7).

We captured substrate-free hGlyT2^Δ185^ in an inward-open conformation in which the extracellular gate is closed due to formation of a conserved salt bridge between residues R216 (TM1b) and D633 (TM10) (Cα-Cα distance of 9.8 Å). Solvation of TM5 and protrusion of TM1 into the micelle creates a large open cavity facing the intracellular milieu. The intracellular part of TM5 is unwound, just before the helix-breaking G419(X_9_)P429 (Gly(X_9_)Pro) motif, allowing water from the intracellular space to access the second Na^+^-binding site (Na2)^48^. The N-terminal segment of TM1 (TM1a) is bent away from the core of the protein, opening the pathway for solute to be released into the intracellular side. This splaying of TM1 disrupts the interaction between TM1a and the hydrophobic patch on the intracellular part of hGlyT2^Δ185^ that is present in other outward-facing or inward-occluded NSS structures^17,49,50^ as well as our outward-facing conformations of hGlyT2^Δ185^ (Fig. 2e, Supplementary Fig. 13).

We observed a clear density for Cl^−^ in the substrate-free cryo-EM map of hGlyT2^Δ185^. This Cl^−^ is coordinated by conserved residues Y233 (TM2), Q473 (TM6a), S477 (unwound region of TM6a), N509 (TM7), and S513 (TM7) at a mean coordination distance of 3.1 Å ± 0.4 (Fig. 3). We did not, however, observe any densities for Na^+^ at Na1, Na2, or the putative Na3 site, consistent with an inward-open conformational state of the transporter^16,50^ (Fig. 3).

**Fig. 3.**
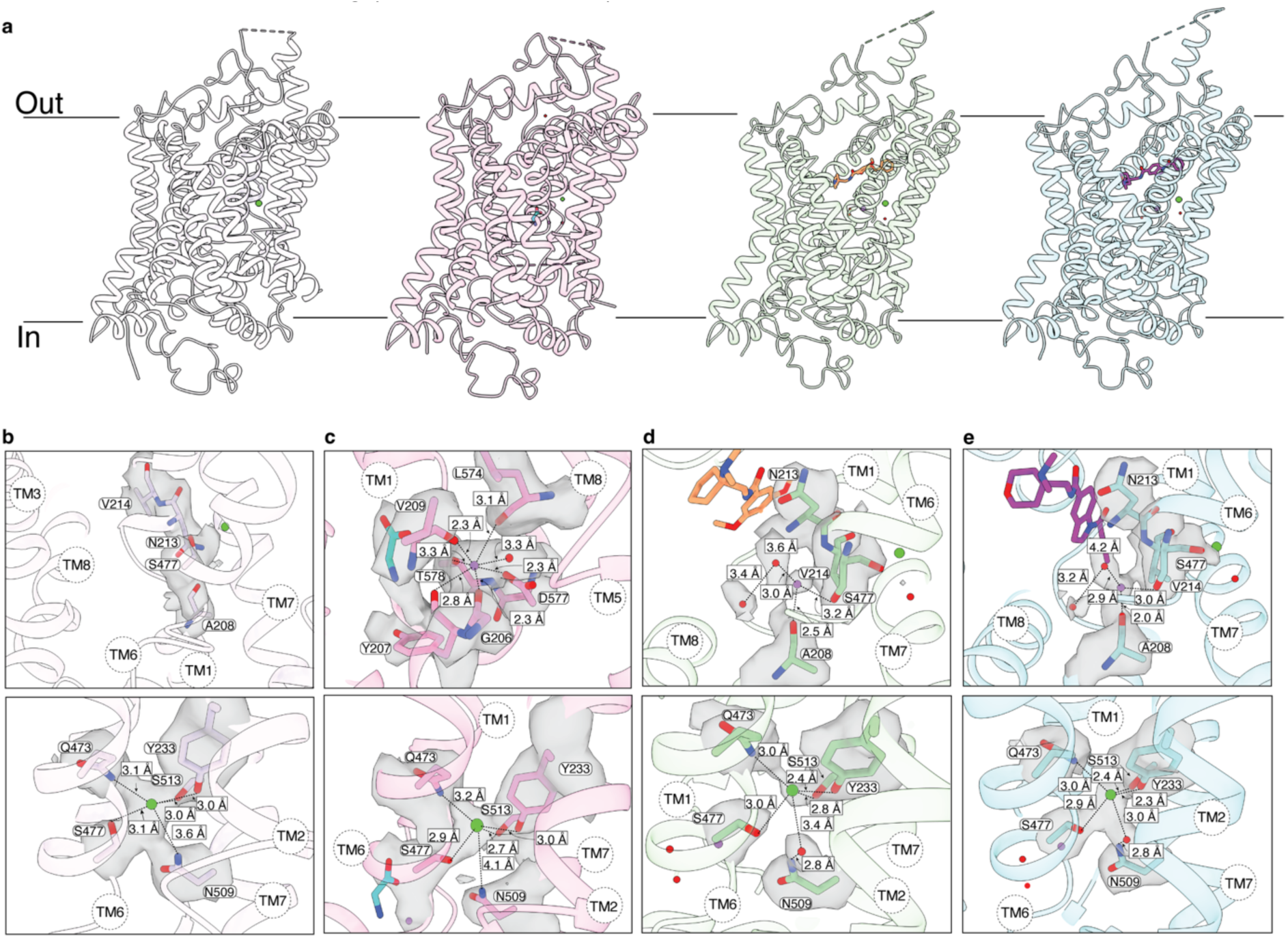
Overall architecture and ion-binding of hGlyT2^Δ185^. **a** Cartoon representation of substrate-free hGlyT2^Δ185^, glycine-bound hGlyT2^Δ185^, ORG25543-bound hGlyT2^Δ185^, and RPI-GLYT2-82-bound hGlyT2^Δ185^. **b** Upper panel is a magnified view of collapsed Na1 site, usual Na1 site-forming residues shown with corresponding densities (contour level = 0.030 in ChimeraX). Lower panel is of Cl^−^ (green) modelled in substrate-free hGlyT2^Δ185^ with coordinating residues (mean coordination distance of 3.1 Å ± 0.4) and densities shown (contour level = 0.030 in ChimeraX). **c** Upper panel is a magnified view of Na^+^ (purple) modelled into Na2 with coordinating residues (mean coordination distance of 2.9 Å ± 0.4) with densities (contour level = 0.046 in ChimeraX). Lower panel is a magnified view of Cl^−^ (green) modelled in glycine-bound hGlyT2^Δ185^ with coordinating residues (mean coordination distance of 3.2 Å ± 0.5) and densities (contour level = 0.046 in ChimeraX). **d** Upper panel is a magnified view of Na^+^ (purple) modelled into the density at Na1 with coordination residues (contour level = 0.050 in ChimeraX). ORG25543 shown in orange and water molecules shown in red (with corresponding densities). Lower panel is a magnified view of Cl^−^ (green) modelled in ORG25543-bound hGlyT2^Δ185^ with coordinating residues (mean coordination distance of 2.8 Å ± 0.3) and densities shown (contour level = 0.050 in ChimeraX). **e** Cartoon representation of RPI-GLYT2-82-bound hGlyT2^Δ185^. Upper panel is a magnified view of Na^+^ (purple) modelled into the density at Na1 with coordination residues (contour level = 0.045 in ChimeraX). RPI-GLYT2-82 is in magenta and water molecules in red (with corresponding densities). Lower panel shows Cl^−^ (green) modelled in RPI-GLYT2-82-bound hGlyT2Δ^185^ with coordinating residues (mean coordination distance of 2.9 Å ± 0.5) and densities (contour level = 0.045 in ChimeraX).

### Glycine-bound hGlyT2 is in an inward-occluded state

Glycine binds within a cavity at the centre of hGlyT2 formed by residues from TMs 1, 3, 6, and 8 (Fig. 2, 4a, b). The glycine-binding pocket is midway through the membrane bilayer and inaccessible from both extracellular and intracellular sides, indicative of an occluded state of the transporter. The extracellular gate residues, R216 and D633, form a salt bridge (Cα-Cα distance of 9.6 Å), which precludes access from the extracellular milieu. The guanidinium of R216 forms a cation-χ interaction with the phenyl group of conserved F476 (TM6a), which sterically blocks the extracellular pathway. The Cα of F476 is 11.3 Å from the Cα of conserved Y287 (TM3), which together cap the glycine binding site (Fig. 4b). The intracellular permeation pathway is occluded by TM1a, which has a more perpendicular arrangement to the membrane than in the substrate-free state (Fig. 2b, f, Supplementary Fig. 13). The distorted density observed in the intracellular half of TM5 (residues 415–419) (Supplementary Fig. 8) suggests an unwound region preceding the conserved helix-breaking Gly(X_9_)Pro motif, creating a solvent pathway from the intracellular milieu to Na2^47^, indicative of an inward-occluded state of glycine-bound hGlyT2.

**Fig. 4.**
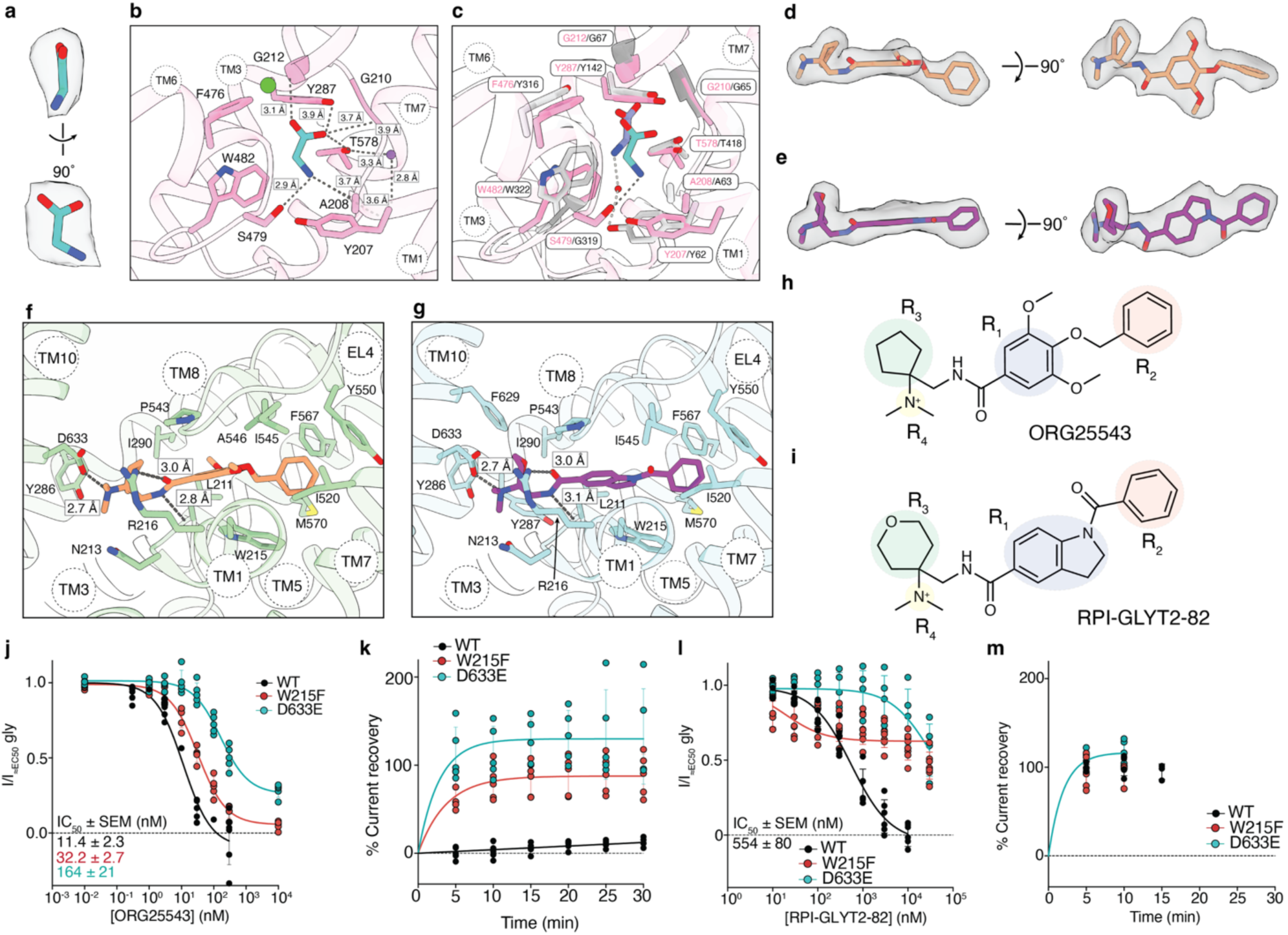
Glycine binds to the central site of hGlyT2, whereas ORG25543 and RPI-GLYT2-82 bind to an allosteric pocket. **a** Cryo-EM density of glycine (map contour level = 0.042 in ChimeraX). **b** Glycine (cyan) bound to the central site of hGlyT2. Interacting residues shown in sticks, H-bonds and ionic interactions shown as dashed grey lines. Na^+^ modelled into Na2 shown as a green circle, modelled Cl^−^ shown as a purple sphere, and modelled water molecule shown as a red circle. **c** Glycine binds in distinct poses in GlyT2 and GlyT1. Glycine-bound hGlyT2 (pink) overlayed with glycine-bound GlyT1 (PDB ID 8WFI, grey). The amine group of glycine (purple) forms a water-mediated interaction with GlyT1 Y62, the glycine (cyan) amino group forms a H-bond with GlyT2 S479. Interacting residues shown as sticks (hGlyT2 = pink, hGlyT1 = black). **d** Cryo-EM density of ORG25543 (map contour level = 0.055 in ChimeraX). **e** Cryo-EM density of RPI-GLYT2-82 (map contour level = 0.04 in ChimeraX). **f** ORG25543 (orange) bound to hGlyT2^Δ185^ (green). Interacting residues shown as sticks and H-bonds and ionic interactions shown as grey dashed lines. N213 is likely involved in a weak hydrogen bond interaction with the polarised C–H of the methyl group attached to positively charged ammonium nitrogen of R4 substituent. **g** RPI-GLYT2-82 (purple) bound to hGlyT2^Δ185^ (blue). Interacting residues shown as sticks, and H-bonds and ionic interactions shown as grey dashed lines. **h** Chemical structure of ORG25543 and key chemical substituents R1 (blue), R2 (orange), R3 (green), and R4 (yellow) outlined. **i** Chemical structure of RPI-GLYT2-82 and key chemical substituents (R1–R4) outlined. **j** ORG25543 concentration-dependant inhibition of glycine transport currents of WT hGlyT2, and mutants W215F (red) and D633E (blue). Data normalised to transport currents generated by the EC_50_ concentration of glycine for each transporter. hGlyT2^WT^ IC_50_ = 11.4 nM, hGlyT2^W215F^ IC_50_ = 32.2 nM, and hGlyT2^D633E^ IC_50_ = 164 nM. **k** Reversibility of ORG25543 inhibition of glycine transport currents mediated by hGlyT2^WT^, hGlyT2^W215F^, and hGlyT2^D633E^ mutants. **l** RPI-GLYT2-82 dose-response curves of WT hGlyT2, and mutants W215F (red) and D633E (blue). Data normalised to glycine transport currents generated by the EC_50_ concentration of glycine for each transporter. hGlyT2^WT^ RPI-GLYT2-82 IC_50_ = 554 nM. The IC_50_ values for hGlyT2^W215F^ and hGlyT2^D633E^ are greater than the maximal tested dose (> 10 µM). **m** RPI-GLYT2-82 is a reversible inhibitor of glycine transport by hGlyT2^WT^, hGlyT2^W215F^, and hGlyT2^D633E^ mutants. Experiments for **j** - **m** performed with n = 5 cells from at least two different batches of oocytes with error bars representing SD.

The carboxylate group of glycine forms hydrogen bonds with the backbone nitrogen atoms of G210 and G212 (TM1), the hydroxyl group of T578 (TM8), and more distantly, a weak hydrogen bond with the hydroxyl group of Y287 (TM3). Although the binding mode of glycine’s carboxylate group resembles that of other α-amino acid transporters, including closely related GlyT1, the positioning of the amino group differs^25^. In hGlyT2, the amino group of glycine appears to form a hydrogen bond with the side-chain hydroxyl of S479 (unwound region of TM6) and is further coordinated by the backbone carbonyl oxygens of Y207 and A208 on TM1, whereas in GlyT1, it adopts a different orientation to interact with the backbone of TM6 residues (Fig. 4c). Residue 479 is a glycine in GlyT1, and the corresponding mutation in GlyT2 (S479G) markedly increases the EC_50_ for glycine transport, confirming the importance of this residue for substrate transport by hGlyT2^26,31^. The binding pocket for glycine is further constrained by W482 in the conserved (G/A/C)ΦG motif (A481W482G483 in hGlyT2) on the non-helical region of TM6, which allows different sized ligands to be accommodated by the various SLC6 family members^51^. A conserved glutamate residue in TM2, E248, coordinates the AWG loop via a direct hydrogen bond with backbone nitrogen of A481 – an interaction that is conserved among SLC6 transporters^51^.

A density for Cl^−^ is clearly apparent in the cryo-EM map of glycine-bound hGlyT2^Δ185^, coordinated by the same residues as the substrate-free state (Y233, Q473, S477, N509 and S513) at a mean coordination distance of 3.2 Å ± 0.5 (Fig. 3, Supplementary Fig. 14). We also observed a prominent (13.5 r.m.s.d.) signal for Na^+^ at Na2, with distorted octahedral coordination by the side chain of D577 (TM8), the backbone carbonyl oxygens of G206 (TM1), Y207 (TM1), V209 (TM1), L574 (TM8), T578 (TM8), and a cytoplasmic water molecule (mean coordination distance of 2.9 Å ± 0.4), similar to previous inward-occluded NSS structures^48^ (Fig. 3c). We did not observe any densities for Na^+^ at Na1 or the putative Na3 site. The conserved S477 residue, equivalent to the Na^+^-coordinating S317 residue at Na1 in GlyT1, is rotated away from the substrate-binding site in glycine-bound hGlyT2, which may suggest that this inward-occluded conformation is in a transitional state.

### ORG25543 and RPI-GLYT2-82 lock hGlyT2 in an outward-open state

In the presence of either ORG25543 or RPI-GLYT2-82, the central substrate-binding site of hGlyT2^Δ185^ is accessible to solvent via the extracellular vestibule, indicating that the inhibitors lock the transporter in an outward-open conformation. The Cα-Cα distance between the two extracellular gating residues, R216 and D633, is 13.9 Å in both ORG25543- and RPI-GLYT2-82-bound structures, consistent with previous outward-facing NSS structures^17,18^. The cytoplasmic pathway is closed by the location of TM1a within the protein core, allowing the conserved residue W194 to plug a hydrophobic pocket formed by residues on TMs 1, 5, 6, and 7 on the intracellular face of GlyT2 (Fig. 2, Supplementary Fig. 11, 13).

In both inhibitor-bound structures, there is a clear density for a Cl^−^, which is coordinated by four of the same residues as those in both inward-facing confirmations (Y233, Q473, S477, and S513; mean coordination distance of 2.8 Å ± 0.3 and 2.9 Å ± 0.5 for ORG25543- and RPI-GLYT2-82 bound structures, respectively. However, Cl^−^ is not directly coordinated by the fifth residue N509 and instead interacts with this residue via a water molecule (Fig. 3d, e). We did not observe any densities for Na^+^ at Na2 or the putative Na3 site^12,13^, consistent with ORG25543^27^ and likely RPI-GLYT2-82, binding being Na^+^-independent (Fig. 1g,h). We observed a prominent (9.7 r.m.s.d.) signal for a non-protein density in the Na1 site of both inhibitor-bound structures. We initially assigned this density as a water molecule because one of the main Na^+^ coordinating residues required for the stable octahedral ion coordination geometry, N213 (TM1b), had rotated away from Na1 to interact with the respective inhibitor (Fig. 3d, e). Modelling of the protein complex with water in the Na1 site using molecular dynamics (MD) simulations showed that extracellular Na^+^ spontaneously bound to this site in four out of six replicate simulations and the site remained occupied by Na^+^ throughout 50% of the 3000 ns combined replicate simulations (Supplementary Fig. 15, Supplementary Table 3). As a result, we subsequently modelled Na^+^ at the Na1 site, coordinated by the backbone carbonyl oxygens of A208 (TM1) and S477 (TM6) as well as a water molecule, reminiscent of a loosely bound hydrated Na^+^ exposed to the extracellular space (Fig. 3 and Supplementary Fig. 15). The increased potency of RPI-GLYT2-82 at lower Na⁺ concentrations likely reflects reduced competition of the compound with Na⁺ for N213 coordination, under partially depleted Na⁺ conditions. In contrast, the potency of ORG25543 remains unchanged, likely due to its inherently high potency that renders it insensitive to local Na⁺ occupancy at the Na1 site (Fig. 1g,h).

### ORG25543 and RPI-GLYT2-82 occupy an allosteric site

The cryo-EM densities of ORG25543 and RPI-GLYT2-82 were of high quality allowing unambiguous assignment of the respective poses when bound to hGlyT2^Δ185^ (Fig. 4d, e). We modelled the inhibitors in an allosteric pocket sandwiched between TM1b, TM10, and EL4 that is lined with predominantly hydrophobic residues from TMs 1, 3, 5, 7, 8, 10 and EL4 (Fig. 4f, g). The scaffold of ORG25543 and RPI-GLYT2-82 has four key substituents (R1–R4) (Fig. 4h, i). In ORG25543 and RPI-GLYT2-82, respectively, these substituents are: the central benzamide phenyl ring and the central indoline (R_1_), benzyloxy, and benzoyl phenyl rings (R_2_), cyclopentyl and tetrahydropyran rings (R_3_), and the pendant ammonium in both compounds (R_4_). For both inhibitors, the central R_1_ group is stacked between W215 (TM1b) and P543 (EL4), and the distal R_2_ ring is caged by a network of residues including W215 (TM1b), I520 (TM7), Y550 (EL4), and F567 (TM8) as well as A546 (EL4) and M570 (TM8). The R_3_ rings are located in a relatively large, solvent-exposed pocket, and stabilized by the hydrophobic residues I290 (TM3), P543 (EL4), and F629 (TM10). The extracellular gate residues R216 and D633 engage in hydrogen bond and salt bridge interactions with both inhibitors (Fig. 4f, g), including a charged-reinforced hydrogen bond and a salt bridge between D633 and the R_4_ pendant ammoniums, and a hydrogen bond between the side chain of R216 and the carbonyl oxygen of R_1_. The allosteric pocket resembles that in MRS7292-bound DAT^20^, suggesting a common allosteric regulatory site in SLC6 transporters. However, this site is distinct from the vilazodone binding site in SERT^22^ (Supplementary Fig. 16).

MD-based r.m.s.d. analysis confirmed stable binding of ORG25543 to the allosteric site for 95.4% of combined simulation replicas. The central benzamide phenyl ring (R_1_) showed minimal fluctuation, due to stabilization by TM1b, EL4, and TM8 (Supplementary Fig. 17, 18, Supplementary Table 4), however the distal benzyloxy phenyl (R_2_) and solvated pendant ammonium (R_4_) had greater mobility (Supplementary Fig. 19). R_2_ frequently interacted with its surrounding residues (Supplementary Table 4, Supplementary Fig. 18) and despite higher fluctuations, R_3_ remained in a hydrophobic cavity flanked by Y286 and Y287 on TM3 (Supplementary Fig. 19).

Hydrogen bond analysis indicated stable interactions between the benzamide of ORG25543 and L211 as well as intermittent interactions with R216. The protonated amine (R_4_) of ORG25543 intermittently engaged in a charged-reinforced hydrogen bond with D633, in agreement with our cryo-EM data (Supplementary Table 4, Supplementary Fig. 18, 19). The probability density map showing the relative likelihood of bound hGlyT2 ligand positions (Supplementary Fig. 19) demonstrated that ORG25543 remained in a deep energetic minimum at the allosteric binding site with an occupancy of 95%. In comparison, RPI-GLYT2-82, adopted a less stable binding pose than ORG25543, exhibiting higher positional fluctuations within the first 50 ns of simulations. It remained in the binding site for only 48% of the combined simulation replicas and moved towards the central vestibule of GlyT2 for the remaining time (Supplementary Fig. 19). Like ORG25543, the R_2_–R_4_ substituents of RPI-GLYT2-82 exhibited the greatest positional fluctuations, and the central indoline of R_1_ formed contacts with the same set of residues that coordinate ORG25543’s R_1_ moiety (Supplementary Table 4, Supplementary Fig. 18, 19).

To analyse MD simulation trajectories of inhibitor-bound hGlyT2 complexes, we estimated binding enthalpy decomposition using endpoint free energy methods. This revealed the significant contribution of L211, W215, and D633 to ORG25543 binding and the 0.83–0.92-fold reduction in binding enthalpy for RPI-GLYT2-82 compared to ORG25543 (95% C.I.) (Supplementary Fig. 19). The per-residue energy decompositions further suggest the lower experimental IC_50_ for RPI-GLYT2-82 stems from an altered interaction enthalpy with L211, R216, and P543 (Supplementary Fig. 19). Together, these data show that both ORG25543 and RPI-GLYT2-82 occupy the same allosteric site, but that RPI-GLYT2-82 adopts a less stable binding pose.

To assess whether glycine can bind to inhibitor-bound hGlyT2 conformations obtained by cryo-EM, we performed 50 ns docking simulations. The observed exchange of solvent and ions between the extracellular space and the substrate-binding site, together with glycine’s mobility within the substrate binding cavity, suggests that glycine can access the ORG25543- or RPI-GLYT2-82–bound hGlyT2^Δ185^ from the extracellular solvent. These findings support a mechanism in which allosteric inhibitors such as ORG25543 and RPI-GLYT2-82 inhibit glycine transport by preventing conformational cycling, a process independent of glycine binding (Supplementary Table 5,6 and Supplementary Fig. 20).

### RPI-GLYT2-82 binds reversibly to hGlyT2

Having characterized the allosteric pocket to which ORG25543 and RPI-GLYT2-82 bind, we sought to determine the features responsible for reversible binding of RPI-GLYT2-82. Likely due to the important role of W215 in accommodating the R_1_ and R_2_ substituents of both inhibitors, a conservative mutation to phenylalanine resulted in a detectable decrease in the potency of inhibition of transport by ORG25543 (∼three-fold) and significant decrease in the potency of RPI-GLYT2-82 (IC_50_ > 30 µM) (Fig. 4j,k). Interestingly, although this W215F mutation had a relatively minor effect on potency, it significantly increased the rate of ORG25543 washout (Fig. 4l), similar to the effect of the mutation of the nearby Y550 (EL4) residue^31^. Likewise, mutation of the neighbouring D633 residue to glutamate significantly reduced the potency of inhibition for both ORG25543 (∼14-fold) and RPI-GLYT2-82 (> 18-fold) and accelerated ORG25543 washout (recovery from ORG25543 within five minutes), highlighting the critical role of this interaction for the potency of the inhibitors (Fig. 4j-l). Notably, washout of RPI-GLYT2-82 from both GlyT2^W215F^ and GlyT2^D633E^ was almost immediate, consistent with the reversible binding mode of the compound (Fig. 4m).

The pendant phenyl rings (R_2_) of both ORG25543 and RPI-GLYT2-82 snorkel deep into the binding pocket to engage in edge-to-face χ-χ interactions with W215 and Y550 (Fig. 4f, g). The R_2_ groups of both inhibitors are positioned orthogonally to their R_1_ core, however, as a benzamide, the pendant benzoyl phenyl ring of RPI-GLYT2-82 is likely to adopt a lower energy coplanar orientation with the carbonyl to which it is attached, reducing conformational flexibility. However, in its binding pose in the allosteric pocket, this RPI-GLYT2-82 pendant phenyl ring must twist nearly 90 degrees to adopt an orthogonal orientation, which likely induces torsional strain and is therefore energetically unfavourable^52^. The indoline benzamide carbonyl oxygen of RPI-GLYT2-82 also points toward a non-polar pocket (Fig. 4g), further contributing to destabilizing contacts. The R_3_ substituent of ORG25543 is a small hydrophobic cyclopentane ring that easily fits into a hydrophobic pocket in the binding site (Fig. 4f), potentially displacing any resident waters and interacting via van der Waals bonds. However, RPI-GLYT2-82 has a tetrahydropyran R_3_ substituent that is larger and more polar than the ORG25543 cyclopentane ring and its oxygen atom is not engaged in any significant binding interactions within the hydrophobic R_3_ pocket of the binding site (Fig. 4g). The increased size of this tetrahydropyran ring coupled with the lack of significant binding interactions with the polar tetrahydropyran oxygen may enable solvent water to “dissolve” the ring out of hydrophobic binding. Thus, the conformational inflexibility of R_2_, together with the larger and more polar R_3_ substituent, are likely responsible for the lower potency and improved washout kinetics of RPI-GLYT2-82 compared to ORG25543.

### RPI-GLYT2-82 provides analgesia in mouse models of neuropathic pain

Although ORG25543 provided analgesia in animal models of neuropathic pain, it caused significant side effects that were likely due to its high affinity and slow dissociation rate^28^. To test the utility of our reversible inhibitor, RPI-GLYT2-82, we conducted plasma and central nervous system pharmacokinetic (PK) profiling in mice and calculated unbound drug concentrations in plasma (*C*_u,p_) and brain (*C*_u,b_) at 30- and 120-minutes post-injection (Supplementary data 1). RPI-GLYT2-82 achieved a *C*_u,b_ of 597.89 nM at 30 minutes following a 50 mg/kg intraperitoneal (i.p.) dose (Supplementary Table 7) – approximately equal to its IC_50_ on hGlyT2, suggesting the compound is likely to achieve meaningful hGlyT2 occupancy *in vivo* (Fig. 1b, Supplementary Table 7).

Encouraged by these data, we tested RPI-GLYT2-82 in the CCI mouse models of neuropathic pain by measuring mechanical and cold allodynia in mice upon i.p. administration. When measuring mechanical allodynia, we observed dose-dependent anti-allodynia at both 50 and 100 mg/kg, with maximum efficacy between 90 minutes and 3 hours post-administration (Fig. 5a). This is comparable to the effect of 30 mg/kg ORG25543, which peaked between 1 and 2 hours post-administration^28^. Similarly, when measuring response by cold allodynia, peak anti-allodynia effects were observed within 2 hours at both 50 and 100 mg/kg with comparable efficacy to the control compound gabapentin (Fig. 5b). We further tested RPI-GLYT2-82 in the PSNL model by measuring its effects on mechanical allodynia and observed similar effects to the CCI model (Fig. 5c).

**Fig. 5.**
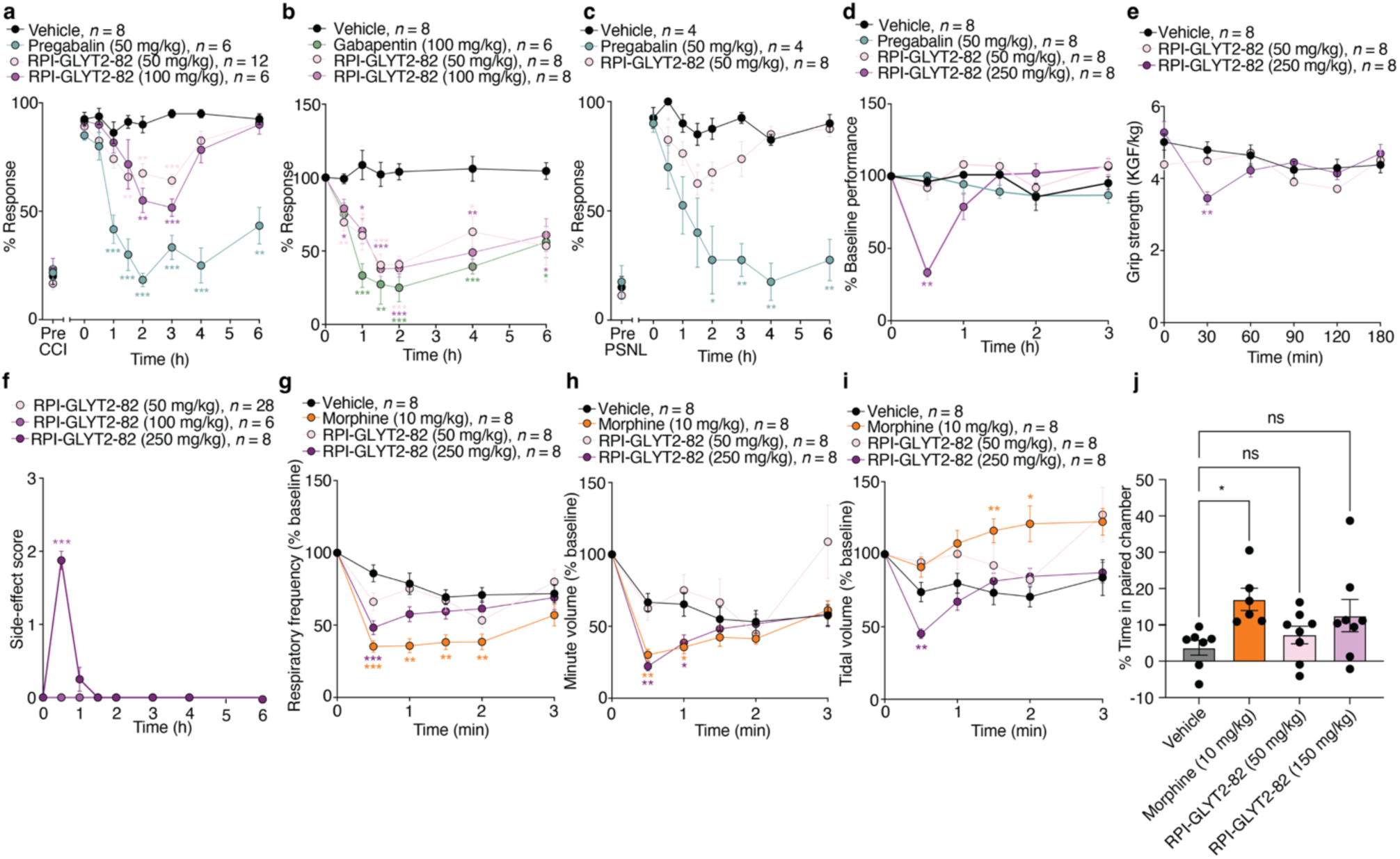
RPI-GLYT2-82 is anti-allodynic in mouse models of neuropathic pain. **a** RPI-GLYT2-82 reduced response rate to von Frey stimulus in CCI mice. Two-way ANOVA with Dunnett’s multiple comparison post hoc test conducted between mice administered vehicle and every other group. ANOVA showed significant effects of time (p < 0.001; F (5, 149) = 27.3) and treatment (p < 0.001; F (3, 28) = 48.3), and an interaction effect (p < 0.001; F (21, 196) = 6.30). Both 50 and 100 mg/kg (i.p.) doses demonstrated efficacy against mechanical allodynia, significant at 2 and 3 hours post-injection compared to vehicle (p < 0.01 & p < 0.001). The 50 mg/kg dose was also significant at 30 minutes (p < 0.01). Both doses peaked at 3 hours, with 30% and 42% reductions, respectively. **b** RPI-GLYT2-82 reduced response rate to acetone stimulus in CCI mice. Two-way ANOVA with Dunnett’s multiple comparison post hoc test was conducted between CCI model mice administered vehicle and every other group. ANOVA showed significant main effects of time (p < 0.0001; F (4, 93) = 25.8) and treatment (p < 0.0001; F (3, 26) = 21.8, along with an interaction effect (p < 0.0001; F (11, 93) = 4.4). Both 50 and 100 mg/kg (i.p.) doses demonstrated efficacy in reducing cold allodynia, significant between 0.5 and 6-hours postinjection compared to vehicle (p < 0.05-0.001). Gabapentin was also significant at 1-6 hours (p<0.05-0.001). All treatments peaked by the 2-hour time point, with 59%, 62% and 75% reductions, respectively. **c** RPI-GLYT2-82 reduced response rate to von Frey stimulus in PSNL mice. Two-way ANOVA showed significant main effects of time (p < 0.001; F (4, 46) = 12.7) and treatment (p < 0.001; F (2, 13) = 20.8), and an interaction effect (p < 0.001; F (14, 91) = 5.86). A 50 mg/kg (i.p.) dose reduced mechanical allodynia, significant at 90 minutes and 2-hours post-injection compared to vehicle (p < 0.05). This effect peaked at 90 mins, with 22.5% reductions. **d** RPI-GLYT2-82 caused a transient reduction in rotarod performance at 250 mg/kg. **e** RPI-GLYT2-82 produced transient deficits in the grip-strength test, indicating muscle weakness at 250 mg/kg but not at 50 mg/kg, 30 min post-injection which was recovered at the 60-minute time point. **f** RPI-GLYT2-82 produced transient sedation at maximal dose. **g–i** RPI-GLYT2-82 (50 mg/kg and 250 mg/kg) examined against positive morphine control (10 mg/kg) and vehicle (saline). Activity of compounds using whole-body plethysmography, measuring **g** respiratory frequency, **h** minute volume, and **i** tidal volume. Two-way ANOVA with Dunnett’s multiple comparison post-hoc test conducted between mice administered saline and every other group. **j** RPI-GLYT2-82 did not increase preference in a conditioned place preference paradigm. RPI-GLYT2-82 (50 and 150 mg/kg) compared to positive morphine control (10 mg/kg) and vehicle (5% solutol, 95% PBS). One-way ANOVA with Dunnett’s multiple comparison post-hoc test conducted between mice administered vehicle and every other group. Data shown as mean ± SEM. Significance is denoted as: *, p ≤ 0.05; **, p ≤ 0.01; ***, p ≤ 0.001. n = biological replicates.

Although ORG25543 reversed mechanical allodynia at a dose of 50 mg/kg, this coincided with near-complete loss of rotarod performance, seizures and in one case, lethality^28^. A 50 mg/kg analgesic dose of RPI-GLYT2-82 was not associated with adverse neuromotor effects or muscle weakness (Fig. 5d-f), and did not change respiration frequency, minute volume (MVb), or tidal volume (TVb) measured by whole-body plethysmography (Fig. 5g–i). A supramaximal 250 mg/kg dose (i.p.), which is five-fold higher than the maximally analgesic dose of RPI-GLYT2-82, produced an initial 61% reduction in baseline rotarod performance, which was restored over the 90-minute time course, and caused transient muscle weakness 30 min post-injection which recovered at the 60-minute time point (Fig 5d, e). Whilst the grip strength and rotarod experiments show mild, transient deficits in muscle strength and motor coordination at the supramaximal analgesic dose, they do not coincide with the anti-allodynic effects. Given the temporal separation of these motor side-effects from the anti-allodynia effects, they do not pose a confound in interpreting the motor response-dependent pain-like behaviors. The supramaximal 250 mg/kg dose of RPI-GLYT2-82 also caused transient changes in respiratory frequency and MVb in the first hour, and TVb in the first 30 minutes (Fig. 5g–i). These effects coincided with transient sedation, which was most apparent in the first 15–30 minutes and fully recovered by 90 minutes (Fig. 5d–f).

These *in vivo* data provide evidence that RPI-GLYT2-82 produces statistically significant dose dependent analgesia without the adverse neuromotor effects associated with ORG25543^28,33^. Interestingly, although the primary target of RPI-GLYT2-82 is GlyT2 (IC_50_ = 554 nM, Fig. 1b), it is also a less potent antagonist at the serotonin 2A (5-HT2A) receptor (IC_50_ = 1.9 µM, Supplementary data 1: Exhibit 14). This additional activity may contribute to its suppression of pain signals. Indeed, antagonism of 5-HT2A has previously been reported for other ORG25543-based GlyT2 inhibitors, including opiranserin^37,38^. RPI-GLYT2-82 also exhibits appreciable inhibitory activity at SERT (IC_50_ = 4.7 µM, Supplementary data 1: Exhibit 14), but is inactive at other off-target sites covered by our Cerep panel screen that are known to promote sedation (γ-aminobutyric acid type A (GABA_A_), histamine (H1), adenosine, adrenergic, and muscarinic receptors) (Supplementary data 1: Exhibit 11). Finally, because the rewarding properties of opioids limit their use in the clinic, we compared RPI-GLYT2-82 and morphine in a conditioned place preference (CPP) paradigm. Mice treated with morphine (10 mg/kg, i.p.) exhibited a significant increase in preference for the paired chamber (p = 0.0287) whereas mice treated with RPI-GLYT2-82 (50 and 150 mg/kg, i.p.) showed no significant preference (p = 0.78 and p = 0.15, respectively) (Fig. 5j). Collectively, our data show that RPI-GLYT2-82 provides a foundation for the development of a non-addictive medication for neuropathic pain.

## Discussion

Recent insights into the pathophysiological adaptations underlying neuropathic pain have provided opportunities to develop new classes of pain-relieving compounds that address limitations of the existing treatments by targeting new mechanisms. The glycine transporter GlyT2 is such a target due to its modulation of glycinergic signalling; however, a structural understanding of transporter inhibition has proved elusive. We have resolved the first structures of hGlyT2 in substrate-bound and substrate-free states as well as bound to two benzamine inhibitors: a potent pseudo irreversible inhibitor that failed in pre-clinical studies, ORG25543, and a novel, reversible inhibitor, RPI-GLYT2-82. We show that RPI-GLYT2-82 is anti-allodynic in both the CCI and PSNL mouse models of neuropathic pain, shows no addiction liability at up to three times the maximal analgesic dose, and is devoid of on-target side-effects. This is an essential milestone that has previously prevented many GlyT2 inhibitors from progressing past the pre-clinical stage.

We reveal that benzamide and related chemotype inhibitors access the transporter from the extracellular side to occupy an allosteric site in a crevice between TM1b and 10 that is capped by EL4, locking the transporter in an outward-open state. This allosteric cavity collapses in the substrate-bound inward-occluded and substrate-free inward-open states of hGlyT2 due to EL4 occupying the space (Fig. 6). In both these inward-facing structures, the TM5 segment is unwound, allowing solvation of the intracellular release pathway. In the absence of substrate, the hinge-like motion of TM1a exposes the core of the transporter to the intracellular side (Fig. 6). We observe a loosely bound Na^+^ in the Na1 site only in the inhibitor-bound structures of hGlyT2 and a Na^+^ in Na2 only in the substrate-bound state. Cl^−^ remains bound in outward-open, inward-open, and inward-occluded conformations of the transporter.

**Fig. 6.**
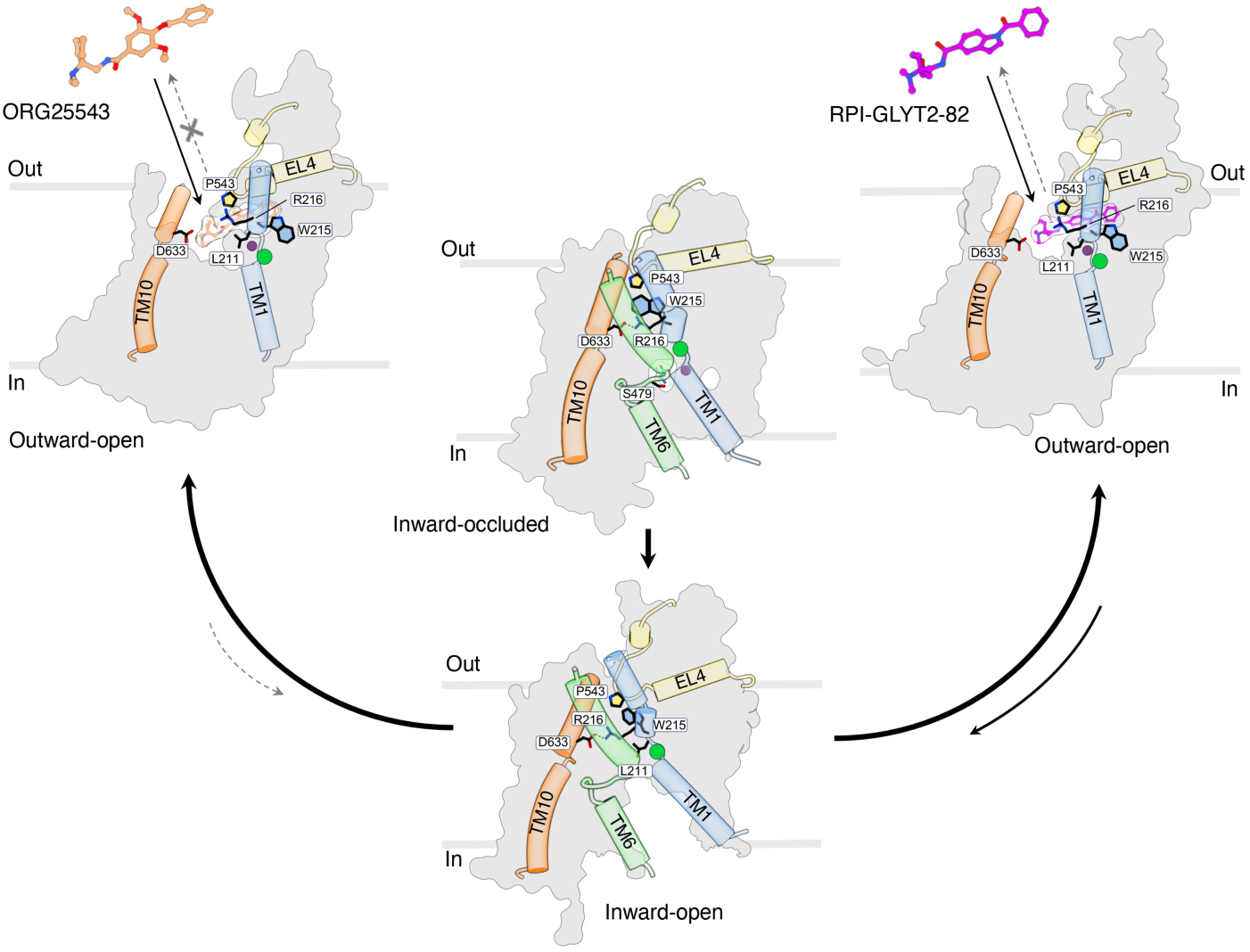
Glycine transport and allosteric inhibition of GlyT2. Schematic showing conformational differences between inhibitor-bound outward-open, glycine-bound inward-occluded and substrate-free inward-open states of GlyT2. ORG25543 (orange) and RPI-GLYT2-82 (magenta) lock GlyT2 in an outward-open conformation. Key binding site residues stabilizing the compounds, L211, W215, R216, P543, and D633, shown as sticks. The slow dissociation rate of ORG25543 prohibits the transporter cycling to an inward-open state. RPI-GLYT2-82 binding to GlyT2 is reversible allowing the transporter to sample other conformational states after release. Glycine is bound to GlyT2 in the inward-occluded state, S479 forming a key interaction to stabilize glycine in the central pocket. In the substrate-free inward-open state, the extracellular gate formed by R216 and D633 closes access to the extracellular side and W215 occupies the allosteric site. Cl^−^ (green circle) is bound in all states, Na^+^ (purple circle) in Na1 is bound only in the inhibitor-bound states, and Na^+^ in Na2 is observed only in the substrate-bound state. Transmembrane helices TM1, TM6, and TM10 shown in blue, green, and orange, respectively, and EL4 is shown in yellow.

In NSS transporters, two ligand-dependent networks coordinate closure of the extracellular pathway and opening of the cytoplasmic one. The first involves the substrate carboxyl group interacting with a tyrosine in TM3 (Y287 in GlyT2) and Na⁺ at the Na1 site; mutation of this tyrosine in the bacterial homologue LeuT prevents the substrate-induced outward-to-inward transition^53^. The second network forms a salt bridge between TM1 and TM10 (R216–D633 in GlyT2), stabilized by Cl⁻ binding through a conserved glutamine residue in the Cl^−^ binding site (Q473 in GlyT2). The interaction between Cl^−^ and Q473 in GlyT2, releases R216 to engage D633, sealing the extracellular gate^54,55^. In GlyT2, transport of Na⁺, Cl⁻, and glycine is tightly coupled, and only their combined binding drives full closure of the extracellular pathway and opening of the cytoplasmic side where the ions and substrate are released^56^. The conserved leucine residue on TM1 (L211 in GlyT2, L25 in LeuT) plays a key role in the inward-to-outward transition. The residue L211 acts as a gatekeeper, occupying the substrate pocket in the absence of ions to prevent leak during the return to the outward-facing state^57^ and vacating the pocket upon Na⁺ and Cl⁻ binding to prime the transporter for glycine binding and forward transport.

The allosteric inhibitors ORG25543 and RPI-GLYT2-82 lock GlyT2 in the outward-open state through two complementary mechanisms: (1) disrupting the extracellular gate by breaking the salt bridge involving R216 and D633, thereby preventing opening of the cytoplasmic pathway and transition toward the inward-facing state, and (2) constraining the conserved leucine residue (L211) outside of the substrate pocket, which further locks GlyT2 in the outward-open conformation (Fig. 4f,g, and 6). Immobilization of the critical gatekeeper residues blocks conformational cycling and traps GlyT2 in a non-transporting state, thereby inhibiting glycine transport.

Intriguingly, ORG25543 and RPI-GLYT2-82 form similar interactions with hGlyT2, suggesting that destabilizing interactions and the lower relative binding enthalpy of RPI-GLYT2-82 contribute to the reduced potency and increased reversibility of this compound. The increased size of the tetrahydropyran ring (R_3_) of RPI-GLYT2-82 likely imposes destabilising binding effects via steric clashes within the local and small hydrophobic pocket it occupies. In contrast, the smaller and more lipophilic cyclopentane ring of ORG25543 has a better fit in this pocket where it likely engages in van der Waals interactions, stabilising the binding pose and helping to anchor the compound within the binding site. These differences will result in a better binding affinity and longer residence time, relative to RPI-GLYT2-82. Although the structures capture stable binding modes of the inhibitors at cryogenic temperatures, our MD simulations provide insight into the dynamic nature of RPI-GLYT2-82 binding. Our simulations show a propensity for RPI-GLYT2-82 to partially enter the central vestibule of hGlyT2 and adopt a more favourable geometry compared to the pose it adopts in the cryo-EM structure, providing an additional explanation for the reversibility of RPI-GLYT2-82 binding. Exploiting this propensity could provide an opportunity to limit the on-target side effects of pseudo-irreversible inhibitors of GlyT2.

The allosteric binding pocket on the extracellular side of the transporter is a particularly interesting pharmacological target due to its sequence diversity among SLC6 transporters and could be exploited to develop highly selective inhibitors for a variety of neurological disorders. We have previously shown that the conservative mutation L425I in EL4 of the closely related transporter, GlyT1, is sufficient to impart sensitivity to the bioactive lipids that inhibit GlyT2^58^. This suggests that a similar allosteric binding site exists in GlyT1. Our results also advocate investigations of the existence of such allosteric sites on the closely related proline and ATB^0,+^ transporters as well as the more distantly related dopamine, serotonin, and GABA neurotransmitter transporters. Recent studies of MRS7292-bound DAT^20^ have revealed an extracellular allosteric site that differs from the GlyT2 site identified here, largely due to low sequence identity. Nonetheless, these two sites share notable structural features, suggestive of a conserved mechanism that may inspire new approaches to drug discovery for the SLC6 family of transporters.

Although RPI-GLYT2-82 provides analgesia in mouse models of neuropathic pain without side effects or addiction liability, it is not as efficacious at alleviating mechanical allodynia as the gold standard neuropathic pain drug, pregabalin (Fig. 5). However, RPI-GLYT2-82 shows equivalent efficacy at alleviating cold allodynia compared to gabapentin and exhibits higher potency (IC_50_ = 0.55 µM) for GlyT2 than opiranserin (IC_50_ = 0.86 µM)^38^. Further, the GlyT2 inhibitors ORG25543 and oleoyl-D-lysine have shown no effects on inflammatory and acute nociceptive pain thresholds and therefore the observed analgesic effects are likely limited to neuropathic pain^28^. Furthermore, because RPI-GLYT2-82 inhibits SERT and the 5-HT2A receptor, as well as GlyT2, it is unclear whether its analgesic effects are, in part, mediated by these other targets and whether sedation at five times the maximal analgesic dose is a result of alternate target occupancy. To overcome these limitations, the selectivity, occupancy and pharmacokinetics of allosteric GlyT2 inhibitors need to be optimized for analgesic efficacy. Given the insights gained from our cryo-EM reconstructions, structure-based design efforts can be rekindled to develop the next-generation of neuropathic pain treatments with improved *in vivo* analgesic efficacy based on optimized RPI-GLYT2-82 analogues. The structure activity- and structure-kinetic relationships presented here can also be used to correlate GlyT2 inhibitor binding profiles and transporter occupancy percentages with analgesic efficacy and therapeutic indices.

We believe the findings presented here reveal the potential that GlyT2 holds as a safe and effective non-opioid target for the treatment of neuropathic pain. The study should also spur re-evaluation of GlyT2 as a target for chronic pain by the broader drug discovery community, facilitating the identification of novel GlyT2 inhibitors that exhibit an optimal blend of inhibition profiles, transporter occupancy, and pharmacokinetics. Neuropathic pain is caused by a wide variety of insults and encompasses a large range of distinct pathological phenotypes. As such it is likely that a range of therapeutics will be required to alleviate the debilitating symptoms. The development of safer and more efficacious GlyT2 inhibitors has the potential to add to the array of therapeutic options as a stand-alone treatment or in combination with other analgesics. This study lays the groundwork for the development of non-opioid pain therapeutics with reduced risk of addiction and greater efficacy against neuropathic pain.

## Supporting information

Supplementary Dara

## Acknowledgments

We thank Shannon N. Mostyn for early contribution to producing the first hGlyT2 constructs. We thank Julian P. Storm and other members of the ASH lab for helpful discussions, and Dr. Roger Dawson for comments on the manuscript. The plasmid for expression of His-tagged HRV-3C protease was a kind gift from Dr. Eric R. Geertsma. We thank Lise Kristensen for access to the protein production facility, and Eva-Marie L. M. A. Pedersen for assistance with virus production, at the Department of Drug Design and Pharmacology, University of Copenhagen. The cryo-EM data was collected at the Core Facility for Integrated Microscopy (CFIM), Faculty of Health and Medical Sciences, University of Copenhagen, supported by the Novo Nordisk Foundation (grants NNF17SA0024386 and NNF22OC0075808). We acknowledge the support offered at the CFIM by Tilmann Pape and Nicholas Sofos. We thank Maria M. Garcia Alai for access to sample preparation and crystallization facility at EMBL Hamburg; and Angelica Struve Garcia, Lucas Defelipe, and David R. Carrillo for technical assistance. We are grateful to Assoc. Prof. David Chalmers for access to his *Silico* software package (http://silico.sourceforge.net). This project has received funding from the Lundbeck Foundation (R368-2021-522), the Novo Nordisk Foundation (NNF23OC0087107), Brødrene Hartmanns Foundation (23080143), and EU Interreg Öresund-Kattegat-Skagerrak project ‘Hanseatic Life Science Research Infrastructure Consortium’ (HALRIC, PP08) to A.S., the National Institute on Drug Abuse, USA, (NIH R01DA048879) to C.L.C. and R.J.V., (NIDA IRP ZIA000069) to M.M and Pain Foundation Ltd to K.R.A. K.E.A. was supported by a Postgraduate scholarship from the University of Sydney. Molecular dynamics simulations were supported by the Australian Government’s National Collaborative Research Infrastructure Strategy (NCRIS), with access to computational resources provided by the National Computational Infrastructure and Pawsey Supercomputing Research Centre through the National Computational Merit Allocation Scheme.

## Author contributions

C.L.C., R.J.V., and A.S. conceived and designed the project. C.L.C. and T.K.P. designed and synthesised RPI-GLYT2-82 compound. R.P.C.C. and A.S. established the protein expression and purification conditions. Expression, purification, cryo-EM sample preparations and data collections, processing, analysis and structure determination were carried out by R.P.C.C. with support from A.S. *In vitro* experiments were carried out by I.L. and R.P.C.C. with support from R.J.V. *In vivo* experiments were carried out by J.P-O, Z.J.F, A.E.T, K.E.A with support from and K.R.A., S.A.M., R.J.V., and M.M. The molecular dynamics studies were carried out by A.S.Q, and B.J.W-N, with support from M.L.O. R.P.C.C., R.J.V., and A.S. wrote the initial draft of the manuscript with contributions from all authors.

## Data and code availability

Atomic coordinates of hGlyT2^Δ185^ bound to ORG25543, RPI-GLYT2-82, glycine, or in substrate-free state have been deposited in the Protein Data Bank (PDB) under accession codes 9HUE, 9HUF, 9R1H, and 9HUG, respectively. The corresponding cryo-EM maps have been deposited in the Electron Microscopy Data Bank (EMDB) under accession numbers EMD-52409, EMD-52410, EMDB-53509, and EMD-52411, respectively. Molecular dynamics starting conformations, mdp files and topologies are available at https://github.com/OMaraLab/GlyT2_2024. This paper does not report original code. Any additional information required to reanalyse the data reported in this paper is available from the lead contact upon request.

## Methods

### Experimental model and subject details

DH5α (New England Biolabs), XL1-Blue Competent Cells (Agilent) and DH10EMBacVSV (Geneva Biotech) were cultured in lysogeny broth (LB) at 37 °C to amplify plasmids. Expi293F cells (ThermoFisher Scientific) were cultured in HE400AZ medium at 37 °C and 8% CO_2_. *Sf9* insect cells (Thermofisher Scientific) were cultured in Sf-900™ III SFM medium at 27 °C. *Xenopus laevis* oocytes were stored at 18 °C in frog Ringer’s solution (96 mM NaCl, 2 mM KCl, 1 mM MgCl_2_, 1.8 mM CaCl_2_, 5 mM HEPES, pH 7.5) supplemented with 2.5 mM sodium pyruvate, 0.5 mM theophylline, 50 μg/mL gentamicin and 100 μg/mL tetracycline.

### Animals

The CCP behavioural experiments were performed with adult male C57BL/6J mice, 8-12 weeks old, obtained from The Jackson Laboratory (Maine, USA). Mice were housed in groups of four and had free access to food and water. Housing rooms were temperature and humidity controlled and maintained on a 12 h light/dark cycle. Experiments were conducted during the light phase.

All other behavioural experiments undertaken with male and female C57BL/6 mice, sourced from the Animal Research Centre (WA, Australia) or Australian BioResources (NSW, Australia). Mice were 6 weeks of age upon arrival and housed in individually ventilated cages (IVC, Sealsafe Plus Green Line, Techniplast) with Smart Flow Air Handling Units (Techniplast), lined with corncob bedding. Mice were housed in groups of 4 in a holding room maintained on a 12-hour light-dark cycle (0545 to 1745) maintained at 21 ± 2 °C and relative humidity at 50 ± 5% with ad libitum access to food (Irradiated rat and mouse chow; Specialty Feeds, WA) and filtered water. Mice were given one week to habituate to the research facility and were then acclimatised to all aspects of the testing procedure including the observation room, handling and scuffing by the experimenter, and the relevant testing apparatus for at least three days prior to commencement of experiments. Animals were monitored at least twice a week for irregularities such as dehydration, skin conditions, and weight loss.

Pharmacokinetic studies were performed by Eurofins Discovery Partner with ICR mice. RPI-GLYT2-82 in-life plasma and BBB pharmacokinetic study was also conducted by Eurofins Discovery Partner with C57BL/6 mice. Male ICR mice weighing 30 ± 5 g and male C57BL/6 mice weighing 22 ± 2 g were provided by BioLasco Taiwan Co., Ltd. (under Charles River Laboratories Licensee). Animals were acclimated for 3 days prior to use and were confirmed to be in good health. The animals were housed in animal cages with a space allocation of 30 x 19 x 13 cm. All animals were maintained in a controlled temperature (20 - 24°C) and humidity (30% - 70%) environment with 12-h light/dark cycles. Free access to standard lab diet [MFG (Oriental Yeast Co., Ltd., Japan)] and autoclaved tap water were granted.

### Ethics statement

*Xenopus laevis* oocytes were supplied by Victor Chang Cardiac Research Institute and approved by the Garvin Institute/St Vincents Hospital Animal Ethics Committee (Animal Research Authority 23_11). Non-CCP behavioural experiments were performed under approval from the Animal Ethics Committee of the University of Sydney (AEC protocol numbers 2229, 1363 and 2146). CCP behavioural experiments were conducted in accordance with the Guidelines of the Animal Care and Use Committee of the Intramural Research Program, National Institute on Drug Abuse, National Institutes of Health (ASP protocol 23-NRB-43). Pharmacokinetics studies conducted by Eurofins Discovery Partner including housing, experimentation, and animal disposal were performed in general accordance with the “Guide for the Care and Use of Laboratory Animals: Eighth Edition” (National Academies Press, Washington, D.C., 2011) in the AAALAC-accredited laboratory animal facility. In addition, the animal care and use protocol was reviewed and approved by the IACUC at Pharmacology Discovery Services (PDS) Taiwan, Ltd.

### GlyT2 construct

The human GlyT2 complementary DNA sequence was codon optimised and synthetised by Azenta for expression in mammalian cell. The GlyT2 construct contains a N-terminal deletion of residues 2 to 185 (hGlyT2^Δ185^). The sequence of hGlyT2^Δ185^ followed by a G(GGGS)_3_G linker, a HRV 3C protease cleavage site, enhanced green fluorescent protein (eGFP), a twin Strep-Tag II tag and a decahistidine tag were cloned into a pCDNA3.1 vector for transient transfection in Expi293F human cells (ThermoFisher Scientific). For glycine-bound hGlyT2^Δ185^, the same construct and tags were cloned into pACEMam1 from the MultiBacMam Kit (Geneva Biotech) for baculovirus production in *Sf9* insect cells (ThermoFisher Scientific). The Expi293F cell stock was authenticated by ThermoFisher Scientific. The cell line was regularly tested for mycoplasma contamination and tested negative for contamination. No misidentified cell lines were used.

### Chemical Synthesis of RPI-GLYT2-82

The general procedure for the synthesis of RPI-GLYT2-82 is described in Supplementary methods.

### Expression of hGlyT2^Δ185^

The hGlyT2^Δ185^ protein was expressed in Expi293F cells cultured in HE400AZ expression medium (GMEP Cell Technologies, Japan) in 600 mL TubeSpin bioreactors, incubating in an orbital shaker at 37 °C, 8% CO_2_ and 175 rpm in a humidified atmosphere. The Expi293F cells (ThermoFisher Scientific; authenticated and free of mycoplasma contamination) were transfected at a density of 3.0 – 3.5 × 10^6^ cells per mL and a viability of above 97%. A 25 kDa linear polyethylenimine (LPEI) was used as the transfection reagent, at a GlyT2 DNA:LPEI ratio of 1:2 with a total plasmid concentration of 2.0 mg per mL of culture volume to transfect. The LPEI/DNA complexes were incubated in Opti-MEM™ I Reduced Serum Medium (ThermoFisher Scientific) for 15 minutes before adding to the cells. At approximately 17 hours post-transfection, cell cultures were supplemented with 10 mM sodium butyrate. The cells were harvested at approximately 44 h post-transfection at a viability of above 70% and stored at -70 °C until purification.

For glycine-bound hGlyT2^Δ185^, baculovirus was generated by transfecting *Sf9* insect cells. The bacmid was prepared by transforming the pACEMam1 vector containing the hGlyT2^Δ185^ construct into *Escherichia coli* Dh10Bac competent cells (Geneva Biotech). For transduction, 1.5 μg of purified bacmid was incubated in 100 μL of Sf-900^TM^ III SFM media and separately 0.1 mg of Cellfectin^TM^ II Reagent (ThermoFisher Scientific) incubated in 100 μL of Sf-900^TM^ III SFM media for 5 minutes at room temperature. Subsequently, the bacmid DNA and Cellfectin^TM^ II mixtures were combined and incubated for a further 20 minutes. The mixture was added dropwise to a well in a 6-well plate of attached *Sf9* insect cells at 0.45 × 10^6^ cells per mL (2 mL final volume). Following 4 hours of incubation at 27 °C, media was exchanged for fresh Sf-900^TM^ III SFM media supplemented with 2% heat-inactivated FBS (Gibco). P0 was harvested following 72-hour incubation at 27 °C by centrifuging at 650 × g for 20 minutes and collecting the supernatant. P1 and P2 were subsequently generated by transducing *Sf9* insect cells at 2.0 × 10^6^ cells per mL in Sf-900^TM^ III SFM media which was then supplements with 2% heat-inactivated FBS at time of harvest. For expression in Expi293F cells, hGlyT2^Δ185^ baculovirus was added at 1.5% (v/v) to the cells at a density of 3.0 – 3.5 × 10^6^ cells per mL in HE400AZ expression medium. At time of transduction, cell cultures were supplemented with 10 mM sodium butyrate. The cells were harvested at approximately 44 h post-transduction at a viability of above 70% and stored at -70 °C until purification.

### Purification of GlyT2

The cell pellets were thawed on ice and resuspended in 50 mM Tris-HCl (pH 7.5), 150 mM NaCl, 20% glycerol (v/v), 1 cOmplete^TM^ Protease Inhibitor Cocktail tablet per 50 mL of buffer (Roche), 2 mg of DNAse I and incubated on rotation for 30 minutes at 4 °C. The suspension was solubilised in 1% (w/v) n-dodecyl-β-D-maltopyranoside (DDM) supplemented with 0.1% cholesterol hemisuccinate (CHS) with 15-25 μM brain polar lipids extract (Avanti) for 45 minutes at 4 °C. Cell debris was removed by centrifugation at 140, 000 g, 4 °C for 20 minutes. To the cleared lysate, StrepTactin XT Sepharose chromatography resin (Cytiva) was added in batch and incubated overnight at 4 °C, with stirring.

The resin was washed with 10 column volumes (CV) of buffer containing 50 mM Tris-HCl (pH 7.5), 150 mM NaCl, 0.1% (w/v) GDN, 0.1% (w/v) LMNG, 0.02% (w/v) CHS and 15-25 μM brain polar lipids extract. The resin was subsequently washed with 20 CV of the same buffer supplemented with 10 mM MgCl_2_ and 2 mM adenosine 5′-triphosphate disodium salt hydrate (ATP). The resin was then washed with 10 CV of buffer containing 50 mM Tris-HCl (pH 7.5), 150 mM NaCl, 0.05% (w/v) GDN, 0.05% (w/v) LMNG, 0.01% CHS, and 15-25 μM brain polar lipids extract. The protein was eluted in the same buffer supplemented with 20% glycerol (v/v) and 50 mM biotin (pH 8). The protein was then treated with HRV-3C protease (in-house) for 3 hours at 10 °C to cleave the eGFP–twin-Strep-Tag-His tag, while being dialysed in buffer containing 50 mM Tris-HCl (pH 7.5), 150 mM NaCl, 0.003% (w/v) GDN, 0.003% (w/v) LMNG, 0.0006% (w/v) CHS and 20% glycerol (v/v). The eGFP–twin-Strep-Tag-His tag was removed by passing the sample through a nickel-nitrilotriacetic acid (Ni-NTA) HisTrap^TM^ HP column (Cytiva) and impurities were removed by anion exchange chromatography with a pre-packed 1 mL HiTrap^TM^ Q HP column (Cytiva). The sample was concentrated to above 2 mg per mL, using an Amicon Ultra-15 centrifugal filter unit (Merck) with a 100,000 dalton cut-off, flash frozen in liquid nitrogen and stored at -80 °C.

On the day of grid preparation, the sample was further purified and buffer exchanged by size exclusion chromatography in buffer containing 50 mM Tris-HCl (pH 7.5), 150 mM NaCl, 0.008% (w/v) GDN, 0.008% (w/v) LMNG, 0.0016% (w/v) CHS using a Shodex kW803 column connected to an Äkta Go purifier system. Fractions were concentrated to above 4 mg per mL and ultracentrifuged at 100,000 g for 30 minutes.

For purification of the ORG25543-bound hGlyT2^Δ185^ sample, cell pellets were resuspended in the buffer supplemented with 100 μM ORG25543 (Tocris), and 10 μM ORG25543 was maintained in all the buffers throughout the purification. 1 mM glycine was maintained in all buffers throughout the purification of the glycine-bound hGlyT2^Δ185^ sample. For the substrate-free GlyT2^Δ185^ sample, resuspension buffer was supplemented with 100 μM oleoyl-D-lysine and subsequent buffers were supplemented with 10 μM oleoyl-D-lysine throughout the purification. For the RPI-GLYT2-82-bound hGlyT2^Δ185^ sample, the protein was purified without inhibitor and was incubated with 100 μM RPI-GLYT2-82 for 45 minutes on ice prior to SEC on the day of grid preparation. The SEC buffer was supplemented with 100 μM RPI-GLYT2-82.

### Cryo-EM grid preparation and data collection

Immediately prior to grid preparation, the protein sample was supplemented with 0.01% (w/v) of fluorinated octyl maltoside (Anatrace). UltrAuFoil grids (R1.2/1.3 300 mesh, Quantifoil) were glow discharged (Leica EM ACE200) at 15 mA for 90 s. 3 μL of sample was applied to the grids, blotted for 5 s with a blot force of -3, or blotted for 4 s with a blot force of -4 before being plunge-frozen in liquid ethane using a Vitrobot mark IV (ThermoFisher) at 100% humidity and 4 °C in an environmental chamber. Movies were collected in counting mode at 165,000× magnification (300 kV), yielding a raw pixel size of 0.728 Å per pixel, using EPU 3.6 software (ThermoFisher) on a Titan Krios G2 (ThermoFisher) equipped with a Selectris X energy filter set to a slit width of 10 eV and a Falcon 4i direct electron detector (ThermoFisher). A defocus value range of –0.6 to –1.8 μm and a dose rate of around 10.2 electrons (e) per pixel per second were used, giving a total dose of 60 e/Å^2^ per movie (Supplementary Table 2).

### Cryo-EM data processing

The cryo-EM data sets were processed using cryoSPARC v.4.5.3^59^. A total of 12,374 movies were collected for the ORG25543-bound hGlyT2^Δ185^ sample, 16,094 for the RPI-GLYT2-82-bound hGlyT2^Δ185^ sample, 14,962 for glycine-bound hGlyT2^Δ185^, and 21,335 for the substrate-free hGlyT2^Δ185^ sample. The movies were motion corrected using Patch Motion Correction with default parameters. The contrast transfer function (CTF) parameters were estimated using Patch CTF Estimation and exposures with a CTF fit resolution of less than 3.6 Å were excluded during curation, yielding 11,378 exposures for ORG25543-bound hGlyT2^Δ185^, 14,629 for RPI-GLYT2-82-bound hGlyT2^Δ185^, 14,892 for glycine-bound hGlyT2^Δ185^, and 20,392 for the substrate-free state.

For the ORG25543-bound hGlyT2^Δ185^ sample, particles were picked by reference-free blob picker using a minimum and maximum particle diameter of 90 Å and 160 Å, respectively. A total of 2,807,000 particles were picked initially. Particles were then extracted with a box size of 300 pixels, Fourier cropped to 256 pixels and used to generate templates following a 2D classification, ab initio, heterogeneous refinement and non-uniform refinement^60^. Particles were then picked with the newly generated templates and extracted with a box size of 300 pixels (Supplementary Fig. 5). Following a 2D classification, a multi class ab initio and subsequent heterogeneous refinement were performed. A further heterogeneous refinement was performed using the selected particles. Suboptimal particles were removed by 3D classification and 2D classification. Selected particles were used for a one-class ab initio followed by a non-uniform refinement with initial low-pass resolution of 12 Å, deactivated dynamic mask start resolution and with the minimize over per-particle scale parameter on. Per-particle movement trajectories and empirical dose weights were estimated using reference-based motion correction followed by a final non-uniform refinement resulting in a 3D reconstruction of ORG25543-bound hGlyT2^Δ185^ map at 2.49 Å resolution, based on a Fourier shell correlation (FSC) cutoff of 0.143 with 283,838 particles.

For the RPI-GLYT2-82-bound hGlyT2^Δ185^ and substrate-free hGlyT2^Δ185^ sample, templates were generated from the ORG25543-bound hGlyT2^Δ185^ final map and used to pick particles. Subsequent data processing was similar to the ORG25543-bound hGlyT2^Δ185^ data processing pipeline (Supplementary Fig. 3, 6). Final non-uniform refinement for RPI-GLYT2-82-bound hGlyT2^Δ185^ and substrate-free hGlyT2^Δ185^ resulted in 3D reconstruction maps at 2.79 Å and 2.97 Å resolution with 201,519, and 180,698 particles, respectively (Supplementary Fig. 3, 6).

For glycine-bound hGlyT2^Δ185^, particles were picked by reference-free blob picker using a minimum and maximum particle diameter of 90 Å and 160 Å, respectively. A total of 3,363,288 particles were picked initially. Particles were then extracted with a box size of 300 pixels, Fourier cropped to 256 pixels and used to generate templates following a 2D classification, ab initio and heterogeneous refinement (using ORG25543-bound hGlyT2^Δ185^ map with decoy classes). Particles were then picked with the newly generated templates and extracted with a box size of 300 pixels and Fourier cropped to 256 pixels. Following a 2D classification, the set of 1,140,447 particles was split into for random batches. The map used to generate templates was further refined by several rounds of heterogeneous refinement, ab initio, 3D classification and non-uniform refinement that generated a 3.51 Å map from 267,885 particles. These 267,885 particles were used and combined individually with the four sets of particles and used to generate 2 class ab initio models. The four sets were combined to two sets by combining particles from the best ab initio class and removing duplicates by 2D classification. This was repeated to result in one combined set of particles that was used to generate a 4.01 Å map from 473, 491 particles following several rounds of heterogeneous refinement and ab initio and a non-uniform refinement of the best class. A molmap focus mask was generated using the substrate-free hGlyT2^Δ185^ model and used in a 3-class 3D classification with a filter resolution of 3 Å and a 5000 O-EM batch size (per class). The particles generating the best class were reextracted at 300 pixels. A 2-class ab initio was followed by several rounds of non-uniform refinement with deactivated dynamic mask start resolution and with the minimize over per-particle scale parameter on and heterogeneous refinement. Reference-based motion correction was then followed by a final non-uniform refinement and finally a local refinement resulting in a 3D reconstruction of glycine-bound hGlyT2^Δ185^ map at 3.02 Å resolution, based on a Fourier shell correlation (FSC) cutoff of 0.143 with 162,255 particles (Supplementary Fig. 4).

### Model building and refinement

The initial hGlyT2^Δ185^ structure model was derived using AlphaFold2 (AF-Q9Y345-F1)^61^. The predicted model was manually fitted into the cryo-EM map in UCSF ChimeraX v1.8^62^ and was initially refined using Namdinator^63^. The models were refined using Phenix v1.21.1-5286^64^ followed by visual examination and manual rebuilding in Coot^65^ v0.9.8.6 and ISOLDE v1.6^66^. The initial Ramachandran outliers were corrected by manual rebuilding and local real-space refinement in Coot. The model was refined in real space using Phenix with the following settings: secondary structure, Ramachandran restraints, global minimization, group atomic displacement factors (ADP) refinement, 5 macro cycles. ISOLDE was then used to simulate the entire model at the start allowing the energy minimiser to alleviate clashes. The remaining Ramachandran and rotamer outliers were corrected using more localised simulations where real-time validation in ISOLDE enabled us to investigate the results of every change directly on the model. Coordinates were written out for a final refinement in Phenix using the input model restraints for the real space refinement. The ligand-restraint files for refinement were generated by phenix.elbow. Figures were prepared using ChimeraX v1.8^62^.

The final models of ORG25543-bound hGlyT2^Δ185^ and RPI-GLYT2-82-bound hGlyT2^Δ185^ lack the initial methionine residue, residues 335–364 in EL2, and the last 13 residues of the C terminus. The final model of substrate-free hGlyT2^Δ185^ lacks residues 185–194 from the N terminus, residues 330–364 in EL2, and the last 12 residues of the C terminus. The final model of glycine-bound hGlyT2^Δ185^ lacks residues 185–194 from the N terminus, residues 327–365 in EL2, and the last 14 residues of the C terminus. Refinement and validation statistics are presented in Supplementary Table 2.

### Expression and functional analysis of GlyT2^WT^, GlyT2^Δ185^, and mutant GlyT2 in *Xenopus laevis* oocytes

Stock solutions of 50 mM ORG25543, and 20 mM RPI-GLYT2-82 were dissolved in dimethyl sulfoxide (DMSO) and diluted in frog Ringer’s solution (96 mM NaCl, 2 mM KCl, 1 mM MgCl_2_, 1.8 mM CaCl_2_, 5 mM HEPES, pH 7.5) to desired concentrations. Final solutions contained less than 0.001% DMSO, a concentration that had no effect on transporter function.

Human GlyT2a wild type (herein referred to as GlyT2) and codon optimised hGlyT2^Δ185^ complementary DNA were subcloned into the plasmid oocyte transcription vector (pOTV). Site-directed mutagenesis was performed using traditional PCR techniques and sequenced by the Australian Genome Research Facility (Sydney, Australia).

Purified plasmid DNAs were linearised with SpeI (New England Biolabs, Cat#R3133S). Complementary RNA was transcribed by the T7 RNA polymerase using the mMESAGE mMACHINE T7 kit (Ambion, Cat#AM1344).

Stage V *Xenopus laevis* oocytes were detached from the lobe by digestion with 2 mg/mL collagenase A (Boehringer, Mannheim, Germany). Defolliculated stage V oocytes were injected with 20 ng of cRNA encoding hGlyT2^WT^, hGlyT2^Δ185^, or mutant GlyT2 (Drummond Nanoinject, Drummond Scientific Co., Broomall, PA, USA). The oocytes were stored at 18 °C in frog Ringer’s solution, which was supplemented with 2.5 mM sodium pyruvate, 0.5 mM theophylline, 50 μg/mL gentamicin and 100 μg/mL tetracycline for 3-6 days, until transporter expression was adequate for measurement using the two-electrode voltage clamp technique. Experiments were performed with at least *n* = 5, from cells precured from at least two individual *Xenopus laevis*.

Whole-cell glycine transport currents were recorded at -60 mV using a Geneclamp 500 amplifier (Axon Instruments, Foster City, CA, USA) with a Powerlab 2/25 chart recorder (ADInstruments, Sydney, Australia). LabChart version 8 software (ADInstruments, Sydney, Australia) was used to process and visualise current traces.

Glycine concentration dependent transport currents were measured for GlyT2^WT^ and hGlyT2^Δ185^ and the mutant GlyT2s (*n* = 5) and the mutant GlyT2s (*n* = 5) and fit to the modified Michaelis Menton equation using GraphPad Prism (version 4.0, GraphPad Software, San Diego, USA):

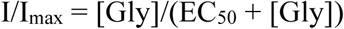

where, I is the transport current generated by each glycine concentration (1-300 µM), I_max_ is the calculated maximal current and EC_50_ is the concentration of glycine that generates a half maximal current.

The potency of inhibition of hGlyT2^WT^, hGlyT2^Δ185^, or mutant GlyT2 was measured by applying the EC_50_ glycine concentration for each transporter and then in the presence of the EC_50_ for glycine with increasing concentrations of inhibitors ORG25543 (0.3–300 nM) and RPI-GLYT2-82 (0.01–10 μM). Currents were fit to the equation:

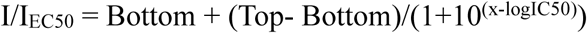

where I is current (nA) generated by EC_50_ for glycine in the presence of the indicated concentration of the inhibitor, and I_EC50_ is the current generated by the EC_50_ (μM) glycine concentration in the absence of inhibitor. Top and Bottom are the maximal and minimal (≥ 0) plateau responses respectively, x is the concentration of inhibitor (nM), and IC_50_ is the concentration that causes 50% maximal inhibition.

The sodium dependence of inhibition by ORG25543 and RPI-GLYT2-82 was measured as described above but with 300 μM glycine to ensure measurable transport currents were obtained at low sodium concentrations. Choline chloride was used as a substitute salt when altering sodium concentrations. The mechanism of inhibition of transport was determined by measuring glycine concentration-dependent transport currents with the co-application of RPI-GLYT2-82 at concentrations < 3-fold IC_50_, IC_50_, and > 3-fold IC_50_.

Reversibility from inhibition was determined by using the previously determined IC_50_ of ORG25543 or RPI-GLYT2-82 (or a maximal dose of 10 μM in cases where the IC_50_ is greater than 10 μM). The glycine EC_50_ for hGlyT2^WT^, hGlyT2^Δ185^, or mutant hGlyT2 was applied and then the IC_50_ of the inhibitor was co-applied until the current plateaued. The cell was then continuously perfused with frog Ringer’s solution to wash out the inhibitor from the bath solution. Every 5 min, glycine (EC_50_) was re-applied for 1 min, and repeated for a total wash period of 30 min or until complete transporter recovery. Washout kinetics were fitted using the equation below:

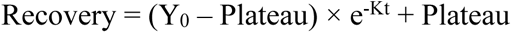

Where recovery is the amount of transport-dependent current restored as a percentage of pre inhibitor levels (%), Y_0_ is the point which crosses the Y axis (≥ 0) and increases to the Plateau (≤ 100) in one phase, K is the rate constant (1/time), and t is time (mins).

### *In vivo* characterisation

#### Drug preparation

Pregabalin (50 mg/kg) and RPI-GLYT2-82 were dissolved in a vehicle of 5% Solutol in PBS. For PSNL experiments, a vehicle of 10% Solutol and 1% DMSO in 0.9% saline was used. For CCI acetone experiments, gabapentain (100 mg/kg) and RPI-GLYT2-82 were dissolved in a vehicle of 10% Cremophor and 5% Tween80 in 0.9% saline. All solutions were prepared on the morning of experimentation in 3-5 mL glass v-vials immediately following baseline behavioural measurements. Mice were randomly assigned to a drug group by a second researcher using randomisation software (random.org), and the experimenter was blind to group allocations until after data collection. Compounds were administered via a single 200 μL i.p. injection into the lower right quadrant of the abdomen using 29-G insulin needles.

#### Chronic constriction injury and peripheral sciatic nerve ligation models of neuropathic pain

Neuropathic pain models were established via nerve injury to the left sciatic nerve via chronic constriction injury (CCI)^67^ via 2 mm polyethylene cuff or two 6-0 chromic gut ligatures placed 2 mm apart or partial sciatic nerve ligation (PSNL) via silk suture tightly constricting 1/3 – 1/2 of the nerve proximal to the trifurcation^68^. Mice were allowed to recover for 14 days after surgery prior to behavioural testing.

#### Mechanical allodynia assessment

Changes in sensitivity to a non-noxious mechanical stimulus was assessed by using a modified von Frey test. A single calibre of nylon filament that elicited approximately 15% response rate in naïve animals (0.4 g force) was applied to the plantar surface of the left hind paw until it just bent. Responses were defined as brisk withdrawal, shaking, or licking of the hind paw. Mechanical allodynia was assessed prior to surgery (baseline) and on the day of testing prior to drug administration (t = 0) and at 0.5, 1, 1.5, 2, 3, 4, and 6 hours post injection. Exclusion criteria were response rates > 30% at baseline or < 70% on the test day at t = 0. Data are presented as the percentage of responses out of 10 applications of the filament.

#### Cold allodynia assessment

Changes in sensitivity to a cold stimulus was assessed by using the acetone test^69^. Cold allodynia was assessed by counting the number of pain-like responses (rapid flinching, shaking or licking of the hindpaw) to evaporative cooling induced application of acetone (20 µl) on the plantar surface of the operated hind paw over a 30 second period. Cold allodynia was assessed prior to surgery (baseline) and on the day of testing prior to drug administration (t = 0) and at 0.5, 1, 1.5, 2, 4, and 6 hours post injection. Data are presented as the percentage of maximal possible responses. Data from n = 1 mouse was exclude as it did not display cold allodynia compared to pre-CCI in baseline testing.

#### Rotarod

Motor coordination was assessed in naïve mice using an accelerating Rotarod, using large cylinders (69.5-mm diameter, 99-mm width, 27-cm drop to landing platform) with speed increasing from 4 rpm to 20 rpm over 180 seconds. Mice underwent 3 days of training with 3 training sessions per day at 30-minute intervals. Mice were allowed 5 min of acclimatisation on cylinders at the start of each training day. On test day, mice were tested at t = 0 and then 30, 60, 90, 120, and 180 minutes after drug administration. Data were normalised to % baseline.

#### Grip strength

Grip strength was assessed using a grip strength meter (Columbus Instruments; Columbus, Ohio) consisting of a force transducer with a triangular horizontal grip bar. Forelimb strength was assessed by gently holding mice by the base of the tail and lowering the animal toward the grip bar so that the forelimbs gripped the horizontal bar. The peak tension as the mouse was pulled away from the grip bar was recorded. The measurement was repeated 5 times over 2 minutes, and the average of 5 recordings was normalised to the animal’s body weight, expressed as kilogram-force per kilogram (KGF/kg).

#### Side effect scoring

Throughout any given behavioural assessment, all animals were also monitored for adverse clinical events using a numerical rating scale. The measures assessed included hypoxia (blanched ears, abnormal colouration), sedation (unconsciousness), activity (unresponsive, inactive), hypothermia (piloerection or shivering), pain/discomfort (arching back, vocalisation, paw withdrawal, piloerection, respiratory depression, orbital tightening), hydration (sunken face), tone (weakness, poor coat), convulsions (clonic seizures, rearing), and grimace/discomfort (orbital tightening). Adverse events were recorded at each time point, observed when placed in the testing apparatus, and scored on a scale of 0 – 3 where 0 = none, 1 = mild, 2 = moderate, and 3 = severe. Scores were averaged for each treatment group.

#### Respiratory depression assessment

Respiratory function was assessed in naïve, uninjured mice via a whole-body plethysmography apparatus consisting of two parallel chambers connected to a pressure transducer (Buxco, DSI Instruments, USA). Mice were acclimatised to the chambers for 15 minutes on the two days preceding the experiment. Chambers were covered with a lab coat and divided with an opaque barrier to prevent any unwanted visual stimulation. On the day of experimentation, mice were acclimatised for 20 minutes, the last 5 of which were taken as the baseline. Mice were briefly removed from the chamber for injection, before being returned to the chamber. Respiratory function was then measured for a period of 5 minutes at 30, 60, 90, 120, and 180 minutes post-injection. Respiratory frequency, tidal volume (TVb) and minute volume (MVb) were recorded on the accompanying computer software (Buxco, FinePointe, DSI Instruments).

#### Conditioned Place Preference (CPP)

The CPP apparatus (Stoelting Co.) consisted of one middle (10 cm × 20 cm) and two outer chambers (18 cm × 20 cm). The middle chamber had bare walls with a smooth floor. One outer chamber had stripes on the walls with a smooth floor and the other had circles on the walls with a textured floor. CPP consisted of three stages: pretesting, conditioning, and testing. Pretesting occurred on the first two days. Mice were placed in the middle chamber and allowed to freely explore the entire apparatus for 20 minutes. Time spent in each chamber was recorded using ANY-maze software (Stoelting Co.) and averaged across the two pretest days to determine baseline preferences. For the next eight days mice received intraperitoneal injection of test drugs (morphine, RPI-GLYT2-82) or vehicle on alternate days. Conditioning sessions lasted 40 minutes during which mice only had access to one of the outer chambers. Drugs were administered in the animals least preferred chamber identified during pretesting as this biased approach has been shown to produce reliable CPP with morphine in mice^70^. The CPP test was performed on day eleven using the same conditions as pretesting. Preference score was calculated as the percent change in time spent in the drug-paired chamber on the test day compared to pretesting.

#### Statistical analyses

To compare the glycine EC_50_ values and the inhibitor IC_50_ values between hGlyT2^WT^ and hGlyT2^Δ185^, an unpaired two-tailed T-test was conducted. A n unpaired two-tailed T-test was also conducted to compare inhibition by RPI-GLYT2-82 between hGlyT2^WT^ and hGlyT1^WT^. To compare EC_50_ values and I_300_ values at different RPI-GLYT2-82 concentrations, Brown-Forsythe ANOVA with Dunnett’s T3 multiple comparisons test and a one-way ANOVA with Dunnett’s multiple comparisons test was conducted respectively. To compare inhibitor sodium dependence at 20, 40, and 100 mM, a two-way ANOVA with Tukey’s multiple comparisons test was conducted.

CCI mechanical allodynia, PSNL mechanical allodynia and CCI cold allodynia were analysed by two-way ANOVA with Dunnett’s multiple comparisons test. Side effect scoring and grip strength data were analysed A repeated measures two-way ANOVA with Greisser-Greenhouse correction and Tukey’s multiple comparisons was conducted to determine p values. Respiratory frequency, tidal volume (TVb) and minute volume (MVb) data were averaged and normalised to percentage (%) of baseline and analysed via two-way ANOVA with Dunnett’s multiple comparisons test. To determine CCP p values, a ROUT test (Q = 10%) was used to remove outliers before performing an ordinary one-way ANOVA with Dunnett’s multiple comparisons test to analyse differences between vehicle and drug treated groups.

#### Molecular dynamics simulations

Molecular dynamics simulations of membrane-embedded GlyT2–inhibitors were initiated from the GlyT2 structures with bound inhibitors using the GROMACS 2023 MD package in conjunction with the GROMOS 54a7 forcefield^71^. The GROMOS 54a7 force field was chosen as it has been specifically parameterised to reproduce the experimental solvation free enthalpy and partition coefficients between polar and nonpolar environments for a range of chemical compounds and is ideally suited for lipid and membrane protein simulations. As the cryo-EM model contained a discontinuity between 330 and 364, acetyl and NH_2_ caps were applied to avoid the introduction of inappropriate charges. The disulfide bond (C311 and C320 (EL2)) was explicitly included. The protonation state of the side chains of ionizable residues was assigned according to the pKa predicted using PROpKa version 3.5.1^72,73^.

Parameters for cholesterol hemisuccinate, ORG255432 and RPI-GLYT2-82 were generated using the Automated Topology Builder 3.069 (CHS molid 1736730, ORG25543 molid 1727769, RPI-GLYT2-82 molid 1732579)^74^. The pKa of the ORG25543 and RPI-GLYT2-82 amide groups were estimated with molgpka^75^ (ORG25543 pKa 8.4, RPI-GLYT2-82 pKa 8.4), and the compounds were simulated with the pendant ammonium protonated.

Proper dihedrals of benzamide groups were refined using a tool that will be included in the next release of the ATB^74^ (ORG25543 R_1_–R_3_ benzamide k_Φ_ = 18.31 kJ/mol/deg, RPI-GlyT2-82 R_1_–R_2_ benzamide k_Φ_ = 22.28 kJ/mol/deg, RPI-GlyT2-82 R_1_–R_3_ benzamide = 21.75 kJ/mol/deg). Benzamide dihedrals were fitted to the difference between the energy obtained by DFT (wB97X/6-31Gd) and energy minimisation with the ATB force field for dihedral angles constrained to 25 values between - 180 and 180 degrees.

Two simulation systems were prepared, one using the GlyT2/ORG25543 cryo-EM complex, and a second using the GlyT2/RPI-GLYT2-82 complex. The composition of all MD simulation systems is given in Supplementary Table 8. The cryo-EM structures of each GlyT2 complex, containing ligand, bound chloride and bound sterols, was embedded in an 80% POPC and 20% cholesterol lipid bilayer created using the MemGen webserver^76^. Each system was solvated with simple point charge (SPC) water, chosen for compatibility with the GROMOS 54a7 forcefield, and 150 mM NaCl for physiological relevance^77^. Counterions were added to ensure overall charge neutrality. All simulations were carried out under periodic boundary conditions. Systems were energy minimized using the steepest descent algorithm then equilibrated with decreasing harmonic restraints on the protein, bound ligand, and bound sterols in five sequential 2 ns simulations with a 2 fs timestep (1000 kJ/mol/nm^2^, 500 kJ/mol/nm^2^, 100 kJ/mol/nm^2^, 50 kJ/mol/nm^2^, 10 kJ/mol/nm^2^). Solute hydrogen masses were repartitioned to access longer simulation timesteps^78^ then the repartitioned systems were equilibrated with an additional 2 ns simulation using a 4 fs timestep, with 10 kJ mol^−1^ nm^−2^ harmonic restraints on the protein, ligand, and bound sterols.

The coordinates from the final frame of each equilibrated system were used as the starting configuration for 500 ns production simulations, using a 4 fs timestep. To increase the statistical sampling of the GlyT2/ligand conformation space and ensure convergence of the docked compounds, non-biased production simulations of each system were performed multiple replicate simulations (GlyT2/ORG25543 n= 6, GlyT2/RPI-GLYT2-82 n= 3). Random starting velocities were assigned at the start of each replicate simulation according to a Boltzmann distribution. The temperature was maintained at 310 K using the Bussi-Donadio-Parrinello velocity rescale thermostat and a coupling constant of 0.1 ps^79^. The pressure was maintained at 1 bar using semi-isotropic pressure coupling with the Parrinello--Rahman barostat^80^ using a 5 ps coupling constant and an isothermal compressibility of 4.5 × 10^-5^ bar ^-1^. SETTLE^81^ was used to constrain the geometry of water molecules and LINCS^82^ was used to constrain the covalent bond lengths of the solute. The electrostatic interactions were calculated using the Particle Mesh Ewald summation, and non-covalent interactions were determined via the Verlet scheme with a 1.4 nm cut-off.

Simulations were visualized using VMD v1.9.4a5575^83^. R.m.s.d. and r.m.s.f. analysis was performed on frames collected at 1 ns intervals, using gromacs tools. Energy landscapes were calculated on frames collected at 0.1 ns intervals with PLUMED version 2.9^84,85^. Per residue contacts and hydrogen bonding frequency analyses were performed on frames collected at 1 ns with VMD v1.9.4a5575^83^. Sodium density was performed on frames collected at 1 ns with MDAnalysis version 2.7.0^86^. Molecular Mechanics-Poisson Boltzmann/Surface Area (MM-PB/SA) was used for binding enthalpy decomposition^87,88^. Per ligand enthalpic contributions to binding were calculated with *g_*mmpbsa^89,90^ on the first 50 ns of each replicate simulation, with frames spaced by 1 ns.

#### Analysis of MM-PBSA binding enthalpy

MM-PBSA calculations were performed using the one-trajectory method^87,88,91^ on the first 50 ns of MD simulation time for GlyT2/ORG5543 and GlyT2/RPI-GLYT2-82 systems, where the ligand remained in its experimentally determined pose. Binding entropy changes were not calculated. Frames were extracted at 1 ns intervals, yielding 50 frames per simulation for analysis.

The *g_mmpbsa*^89^ module of GROMACS was used to compute binding enthalpies with a grid spacing of 0.5 Å, a solvent probe radius of 1.4 Å, γ = 0.03 kJ/mol/Å^2^, and an external solute (saline) dielectric constant of 80. To address variations in internal solute dielectric across the full protein, the analysis was restricted to the ligand (experimental pose) and its binding site residues: 211, 212, 213, 215, 216, 286, 287, 290, 433, 520, 541, 542, 543, 545, 546, 547, 550, 567, 570, 571, 574, 629, and 633.

Focusing on the binding site enabled MM-PBSA calculations using a consistent internal solute dielectric value, eliminating contributions from distal residues to the binding site with a different solute dielectric value.

Electrostatic analysis using the APBS webserver^92^ revealed binding site surface electric potentials ranging from approximately -15 to -30 kT/e for most of the binding site. This variation in surface electric potential afforded a solute dielectric range of 10-20 across the GlyT2 binding site. The conversion from surface electric potential to solute dielectric was done using Equation **1**^93,94^. Here, *ϕ* is the surface electrostatic potential (in units of kT/e) from the APBS webserver^92^, *q* is the charge of an electron, *ξ*_0_ is the vacuum permittivity constant, *ξ_s_* is the internal solute dielectric for MM-PBSA, and *r* is set to the interaction cut-off distance from the MD simulations (r = 1 nm). Specifically, *ξ*_0_ is scaled by *ξ_s_*, such that *ξ*_0_*ξ_s_* accounts for the overall permittivity of the medium. Equation **1** is reduced to the relationship from Coulomb’s law when *ξ_s_* = 1.

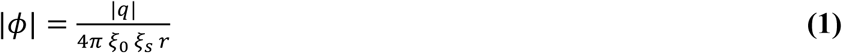

We chose ***ξ***_***s***_= 10 for two reasons: A) Equation **1** afforded values of ***ξ***_***solute***_between ∼10-20; and, B) for ***ξ***_***s***_values between 10-20, the resultant MM-PBSA calculation with ***ξ***_***solute***_= 10 yielded a ratio of RPI-GLYT2-82/ORG25543 binding enthalpy (0.83-0.92, 95% C.I.) that was the most consistent with the experimental log(IC_50_) ratio of RPI-GLYT2-82/ORG25543 (0.79). Specifically, ***ξ***_***solute***_= 10 gave the best qualitative agreement with experiment. Binding enthalpies and per-residue enthalpy contributions were reported as mean values ± the 95% confidence interval in energy (Supplementary Fig. 19). We did not report per-residue binding energy benefits/penalties > 2 kT (5.15 kJ/mol).

#### Substrate docking

Docking was performed on the GlyT2 structure obtained from cryo-EM, with the inhibitor (ORG25543 or RPI-GLYT2-82), the cryogenic sodium ion, and the cryogenic chloride ion placed in their respective binding sites. Zwitterionic glycine was docked into the GlyT2/inhibitor/ions complex using autodock vina^95^. This structure was placed in a 80% POPC/20% CHOL bilayer, with cryogenic cholesterol and cholesterol hemisuccinate placed in their cryo-EM binding sites. The membrane embedded complex was solvated with SPC water containing 150 mM NaCl, with additional ions added to neutralize the overall system charge. To evaluate the possibility of glycine binding to the cryo-EM complexes, these solvated systems were used for a series of five replicate simulations.

Each replicate was equilibrated separately through a series of 2 ns simulations with decreasing position restraints on the protein backbone, cryogenic ions, cryogenic sterols, and docked glycine (fc 1000, 500, 100, 50, 10) with a 2 fs timestep. Following position restrained EQ, solute hydrogen masses were repartitioned to access longer simulation timesteps, then the repartitioned systems were equilibrated with an unrestrained 2 ns simulation using a 4 fs timestep. Coordinates and velocities of the equilibrated systems were used for 50 ns production simulations performed under the conditions described above.

