## Supplementary Dara for "A reversible allosteric inhibitor of GlyT2 alleviates neuropathic pain without on-target side effects"

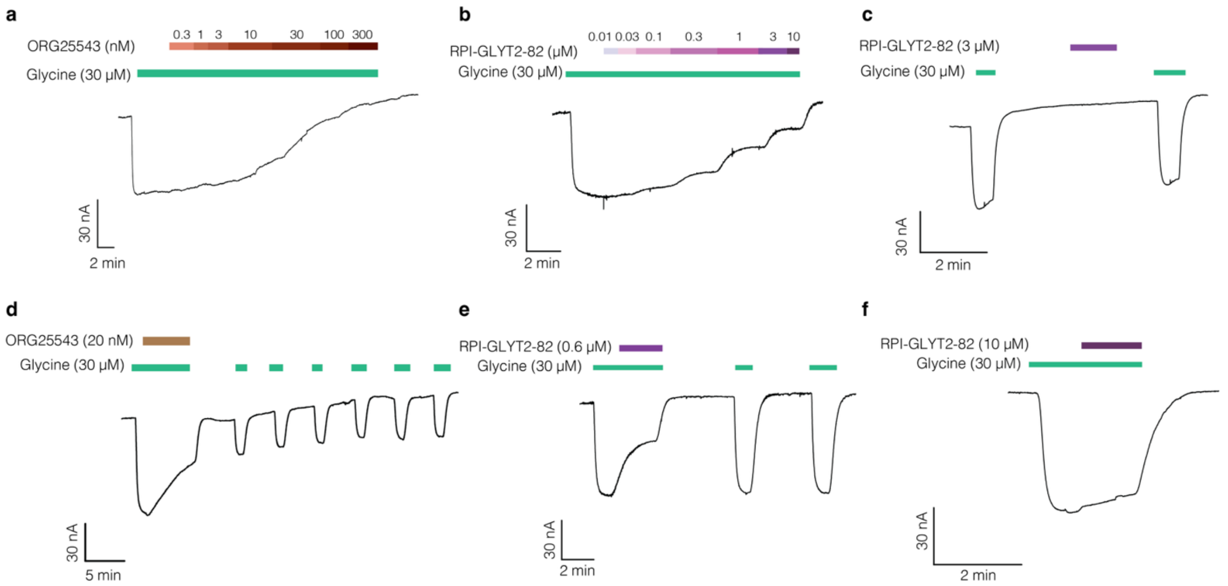

**Supplementary Fig. 1| Electrophysiological characterization of ORG25543 and RPI-GLYT2-82.** Transport activity was characterized by two-electrode voltage clamp electrophysiology with *Xenopus laevis* oocytes expressing hGlyT2$^{WT}$, hGlyT2$^{\Delta185}$ or wild-type hGlyT1. Representative hGlyT2 expressing cells electrophysiology traces of **a** ORG25543 dose-response **b** RPI-GLYT2-82 dose response, **c** RPI-GLYT2-82 activity in the absence glycine, **d** reversibility characterization of ORG25543 and **e** reversibility characterization of RPI-GLYT2-82. **f** Representative trace of hGlyT1 expressing oocyte perfused with RPI-GLYT2-82 and glycine.

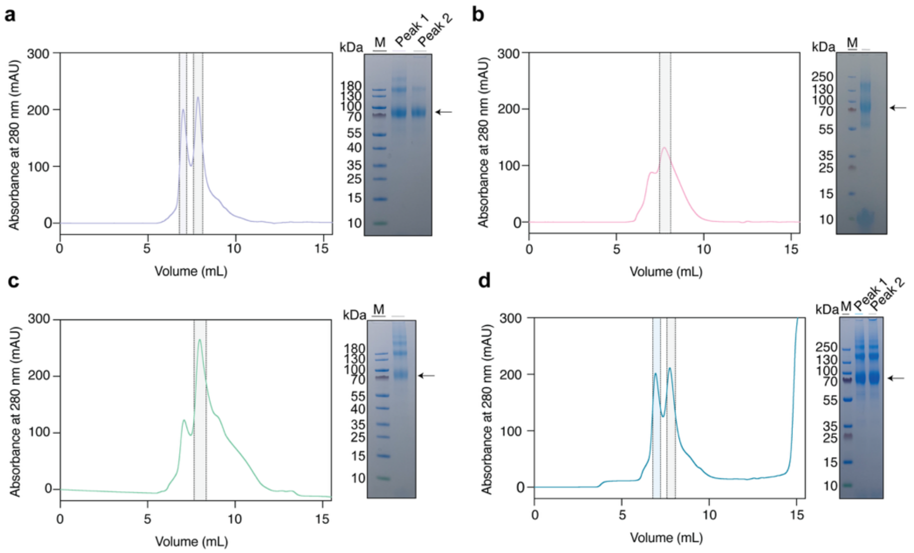

**Supplementary Fig. 2| Purification of GlyT2$^{\Delta185}$.** Chromatogram of absorbance at 280 nm of purified hGlyT2$^{\Delta185}$ from a Shodex KW-803 gel filtration column and corresponding SDS-PAGE gel analysis of pooled fractions. hGlyT2$^{\Delta185}$ was purified by size exclusion chromatography (SEC) in 50 mM Tris pH 7.5, 150 mM NaCl, 0.008 % (w/v) GDN, 0.008 % (w/v) LMNG and 0.0016 % (w/v) CHS with **a** oleoyl-D-lysine (10 µM), **b** glycine (1 mM)**,** **c** ORG25543 (10 µM) or **d** RPI-GLYT2-82 (100 µM). The protein band at ~ 70 kDa represent GlyT2$^{\Delta185}$ and is indicated by an arrow and the ladder is indicated by M. In the case of **a** and **d** peak 1 pooled sample are indicated by coloured shading (purple and blue, respectively) and peak 2 samples are indicated by grey shading. In both cases, peak 2 was pooled and used for cryo-EM studies.

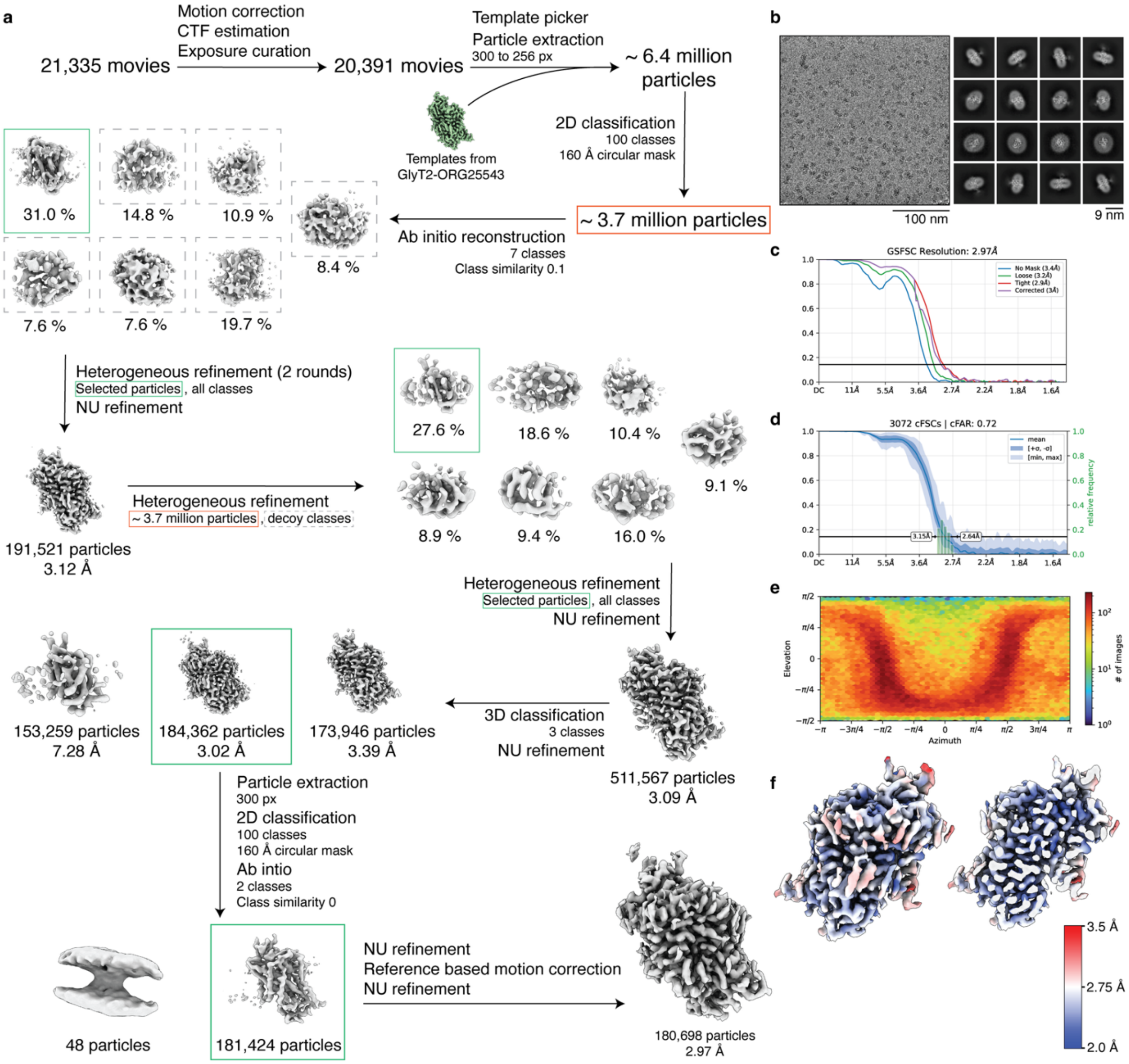

**Supplementary Fig. 3| Cryo-EM data processing and analysis of substrate-free hGlyT2$^{\Delta185}$.** **a** Cryo-EM workflow. In brief, 6,458,661 particles were picked from 20,391 movies using a low-pass filtered ORG25543-bound hGlyT2$^{\Delta185}$ map as a template. Following a 2D classification, 3,759,887 particles were used to generate 7 ab initio classes and after two rounds of heterogenous refinement and a non-uniform refinement, a 3.12 Å was generated. This map was used in a heterogenous refinement with 3,759,887 particles. Following a 3D classification, non-uniform refinements, 2D classification ab initio reconstruction, and reference-based motion correction, the final map was refined to 2.97 Å. **b** Representative motion-corrected micrograph and representative 2D class averages. **c** The gold standard Fourier shell correlation (GSFSC) curve with FSC= 0.143 cut-off represented by a black horizontal line. **d** The conical FSC curve. **e** Angular sampling of the final reconstruction. **f** Local resolution map.

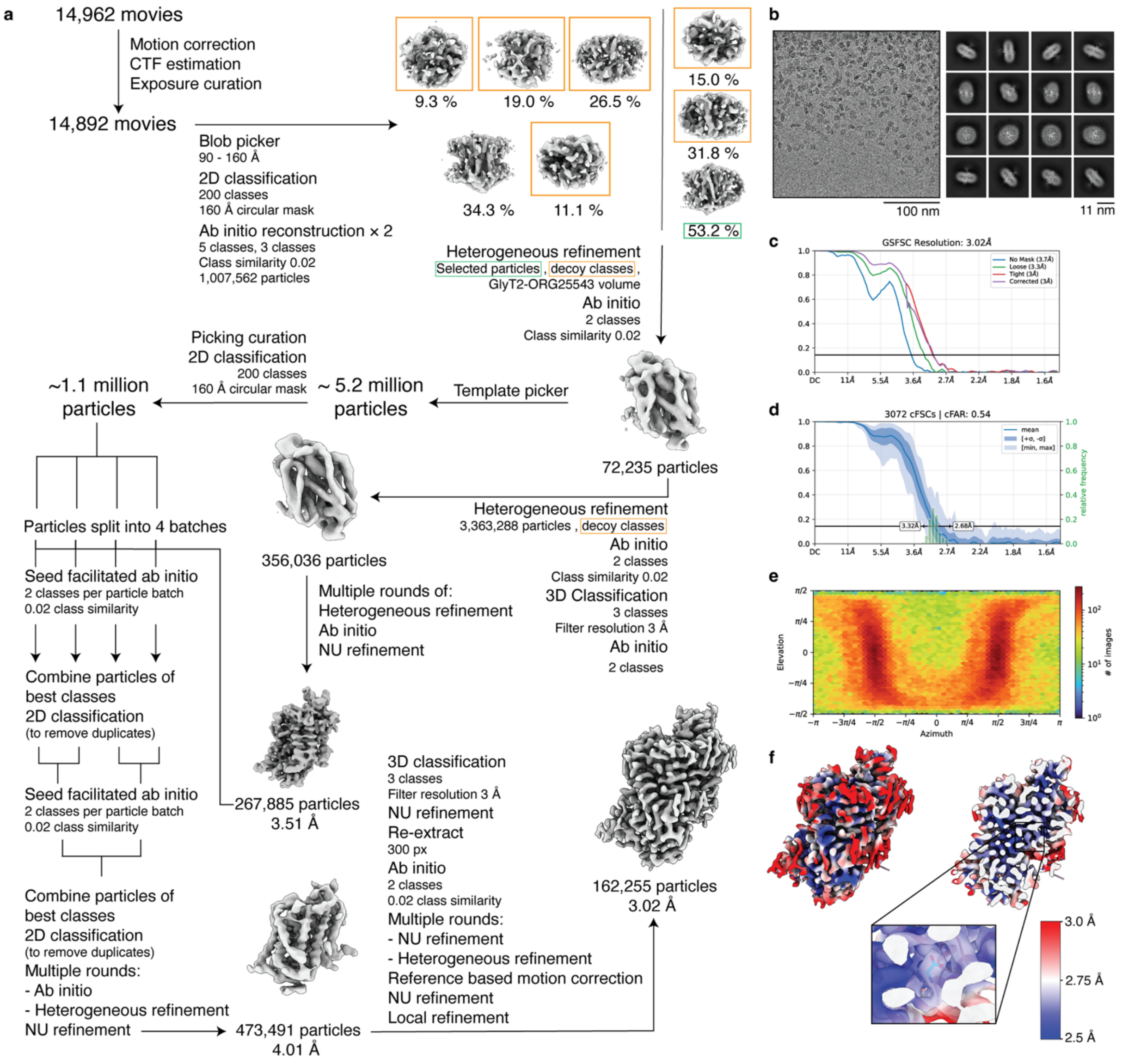

**Supplementary Fig. 4| Cryo-EM data processing and analysis of glycine-bound hGlyT2$^{\Delta185}$.** **a** Cryo-EM workflow. In brief, 3,363,288 particles were picked from 14,892 movies using blob picker. Following a 2D classification, 1,007,562 particles were used to generate 5 and 3 ab initio classes. Heterogeneous refinement was used with ORG25543-bound hGlyT2$^{\Delta185}$ map and decoy classes, and a 2-class ab initio was generated from the best class. The volume was further processed by heterogeneous refinement, ab initio, 3D classification and non-uniform refinement to generate a 3.51 Å map. Particles were then used to seed four independent ab initio processes from template picked particles. Best classes were combined down to two ab initio processes and finally down to one three class ab initio. The best map was further processed by multiple rounds of ab initio, heterogeneous refinement and non-uniform refinement. A focus masked 3D classification was performed followed by non-uniform refinement and heterogeneous refinement and a local refinement generated the final map at 3.02 Å. after two rounds of heterogenous refinement and a non-uniform refinement, a 3.12 Å was generated. **b** Representative motion-corrected micrograph and representative 2D class averages. **c** The gold standard Fourier shell correlation (GSFSC) curve with FSC= 0.143 cut-off represented by a black horizontal line. **d** The conical FSC curve. **e** Angular sampling of the final reconstruction. **f** Local resolution map.

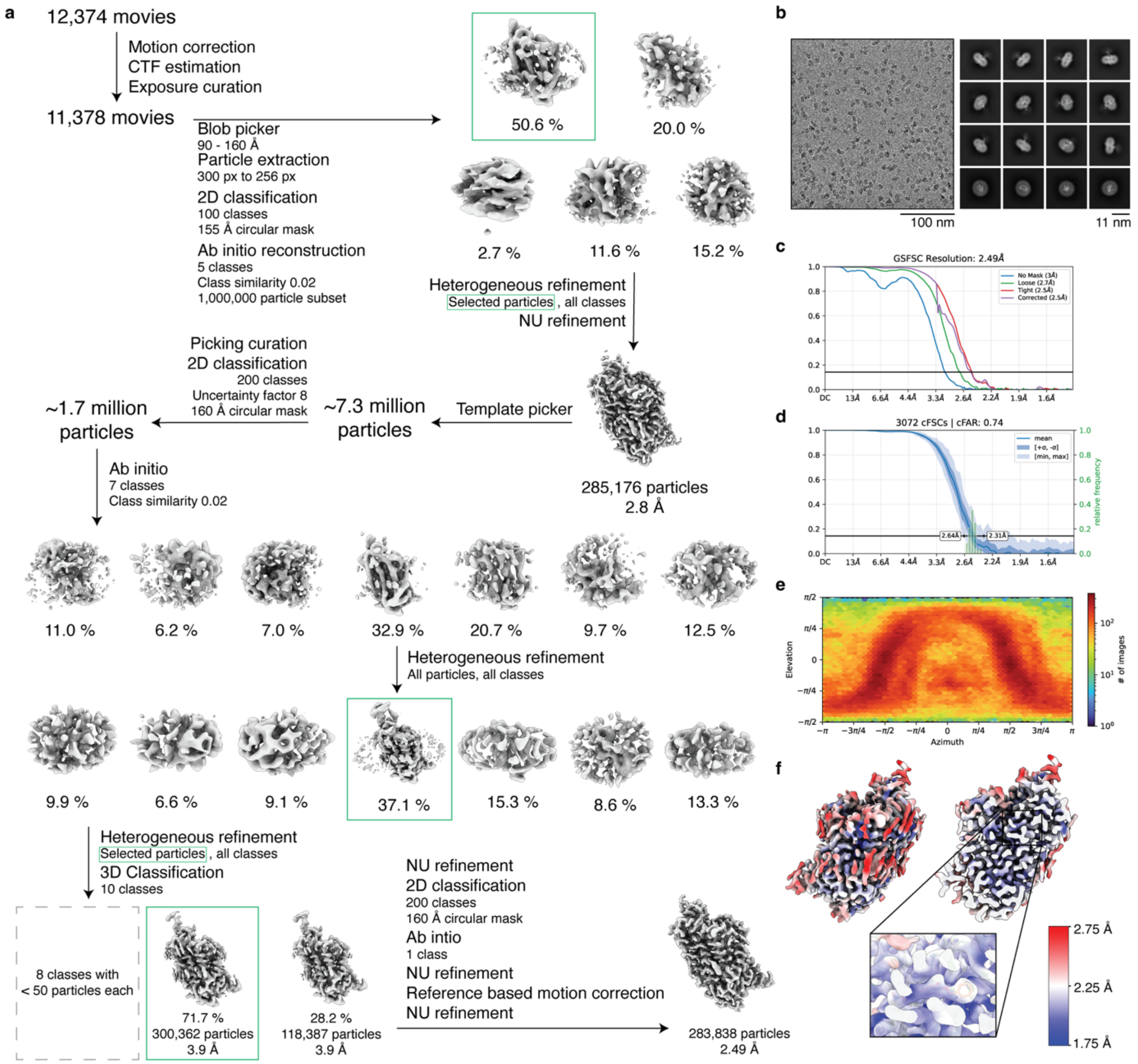

**Supplementary Fig. 5| Cryo-EM data processing and analysis of ORG25543-bound hGlyT2$^{\Delta185}$. a** Cryo-EM workflow. In brief, 11,378 movies were selected after manual curation. 2,807,000 particles were initially picked from which 1,123,791 particles were selected to generate 5 ab initio models. The best model was then used as a template to pick particles. Following a 2D classification, selected particles were used to generate 7 ab initio classes and after two rounds of heterogenous refinement and a 3D classification, a 3.9 Å map was generated. The final 2.49 Å map was generated after several rounds of non-uniform refinement, 2D classification, a reference-based motion correction, and a final non-uniform refinement. **b** Representative motion-corrected micrograph and representative 2D class averages. **c** The gold standard Fourier shell correlation (GSFSC) curve with FSC= 0.143 cut-off represented by a black horizontal line. **d** The conical FSC curve. **e** Angular sampling of the final reconstruction. **f** Local resolution map.

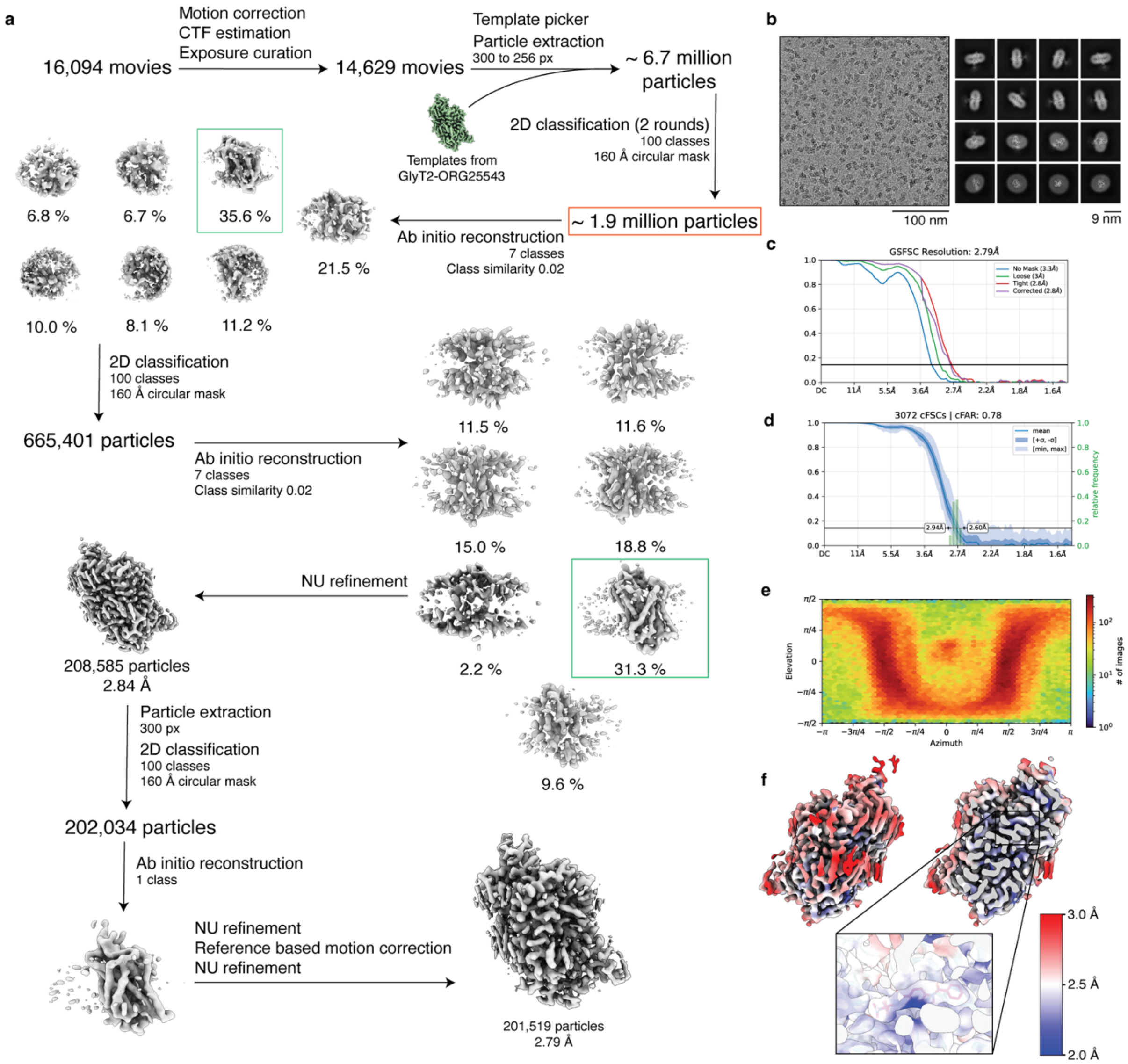

**Supplementary Fig. 6| Cryo-EM data processing and analysis of RPI-GLYT2-82-bound hGlyT2$^{\Delta185}$.** **a** Cryo-EM workflow. In brief, 14,629 movies were selected after manual curation. A low-pass filtered ORG25543-bound hGlyT2$^{\Delta185}$ map was used as a template to initially pick 6,700,966 particles. Following two rounds of 2D classification, 7 ab initio models were generated. The best model was then refined to a 2.84 Å map. Following another round of 2D classification, ab initio reconstruction, non-uniform refinement, and a reference-based motion correction, the final map was refined to 2.79 Å map. **b** Representative motion-corrected micrograph and representative 2D class averages. **c** The gold standard Fourier shell correlation (GSFSC) curve with FSC= 0.143 cut-off represented by a black horizontal line. **d** The conical FSC curve. **e** Angular sampling of the final reconstruction. **f** Local resolution map.

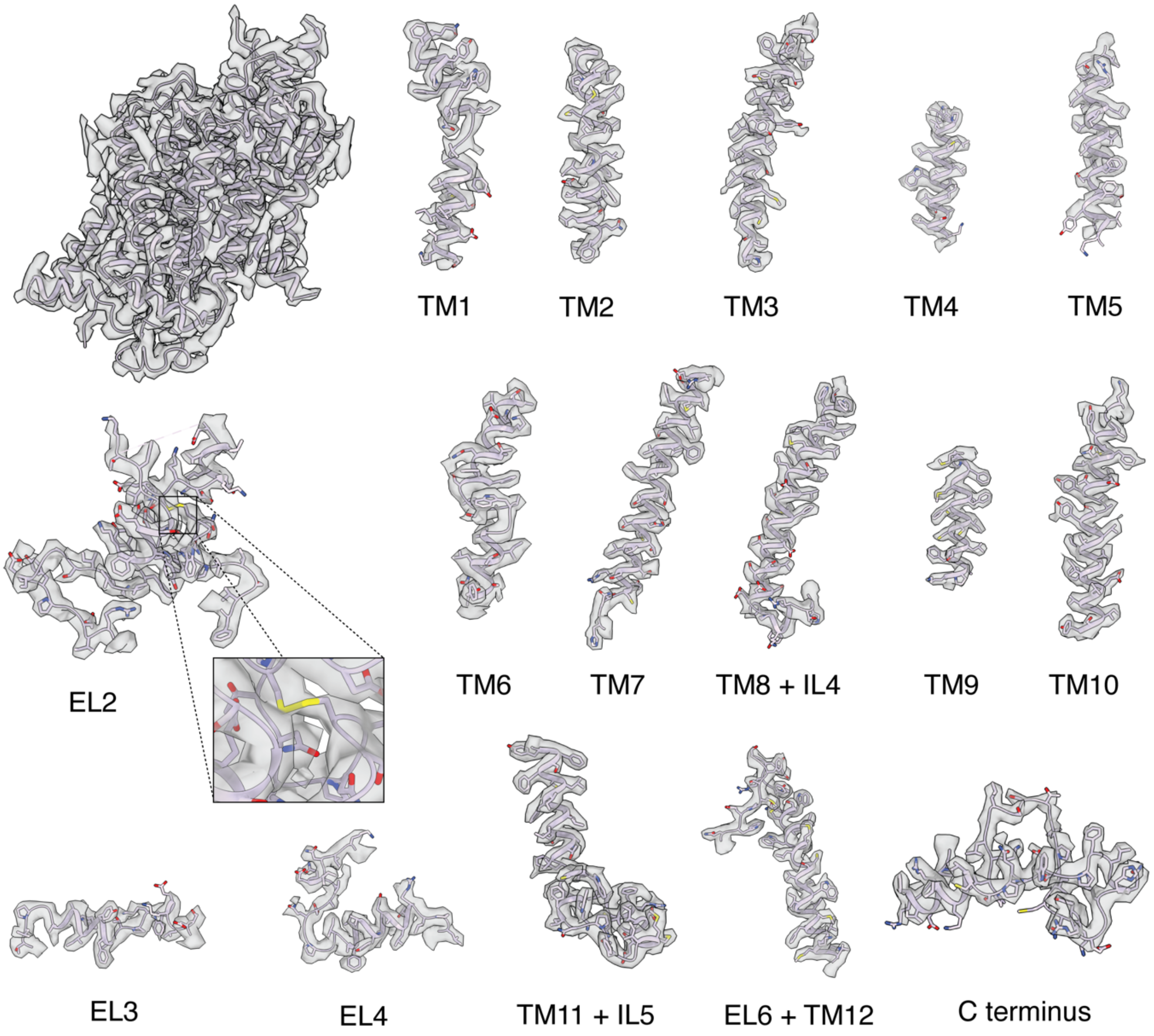

**Supplementary Fig. 7| Cryo-EM density of substrate-free hGlyT2$^{\Delta185}$.** The overall structure of substrate-free hGlyT2$^{\Delta185}$ (top left, PDB ID 9HUG) and magnified density views of transmembrane helices (contour level = 0.04 in ChimeraX), EL2 (with the conserved NSS disulfide bridge), as well as EL3, EL4 and the C terminus are shown.

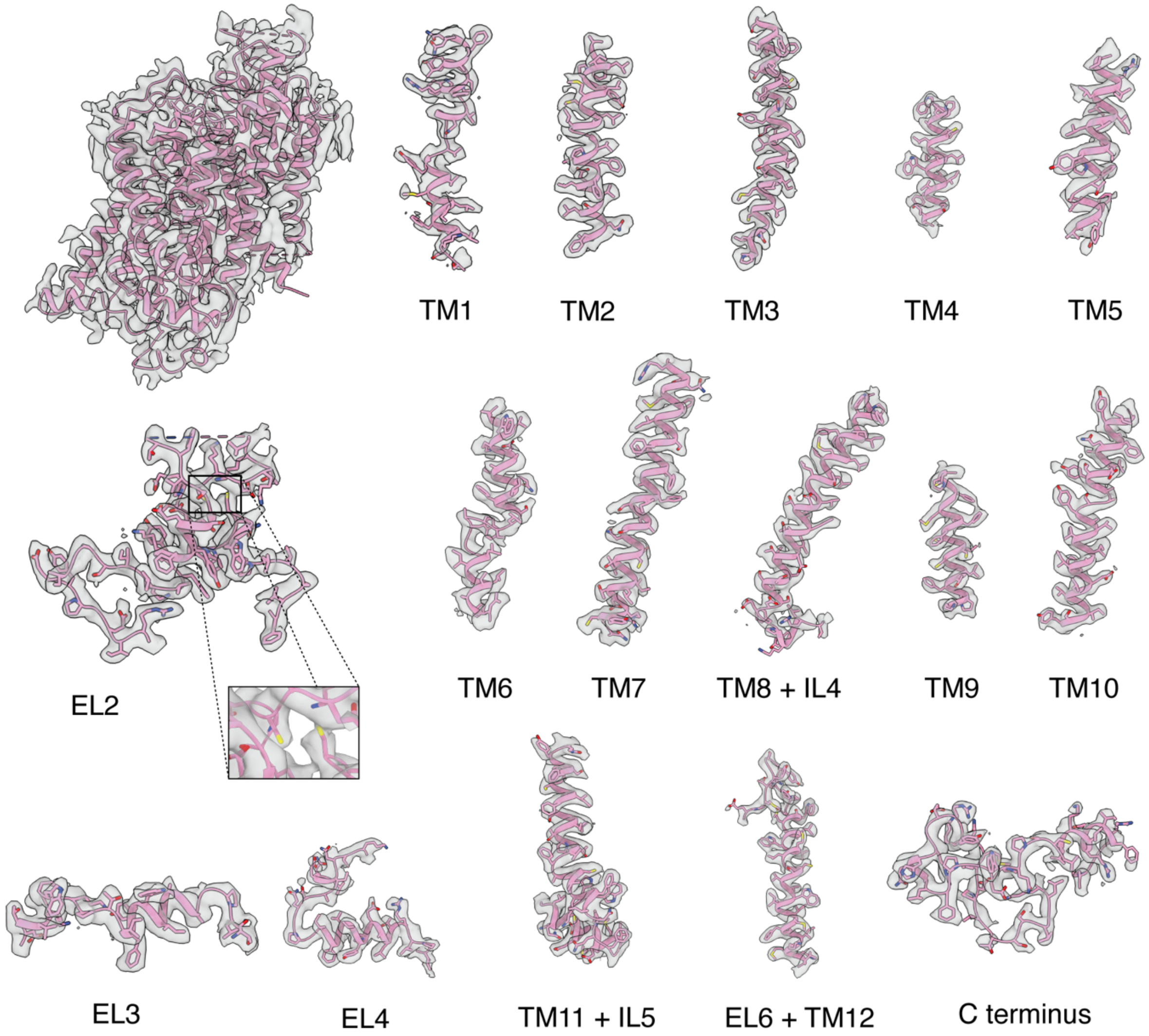

**Supplementary Fig. 8| Cryo-EM density of glycine-bound hGlyT2$^{\Delta185}$.** The overall structure of glycine-bound hGlyT2$^{\Delta185}$ (top left, PDB ID 9R1H) and magnified density views of transmembrane helices (contour level = 0.046 in ChimeraX), EL2 (with the conserved NSS disulfide bridge reduced), as well as EL3, EL4 and the C terminus are shown.

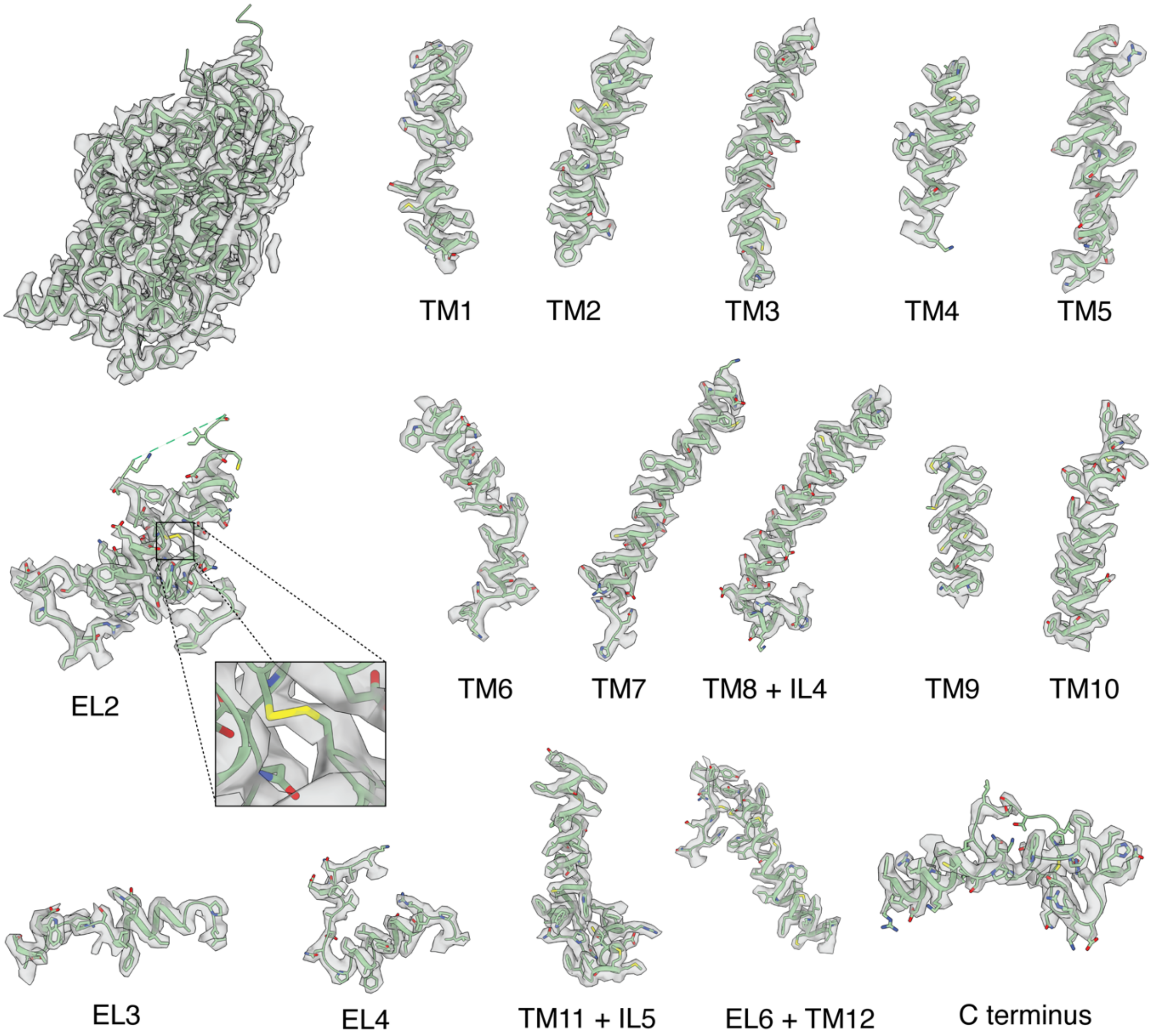

**Supplementary Fig. 9| Cryo-EM density of ORG25543-bound hGlyT2$^{\Delta185}$.** The overall structure of ORG25543-bound hGlyT2$^{\Delta185}$ (top left, PDB ID 9HUE) and magnified density views of transmembrane helices (contour level = 0.055 in ChimeraX), extracellular loop (EL) 2 (with the conserved NSS disulfide bridge), as well as EL3, EL4 and the C terminus are shown.

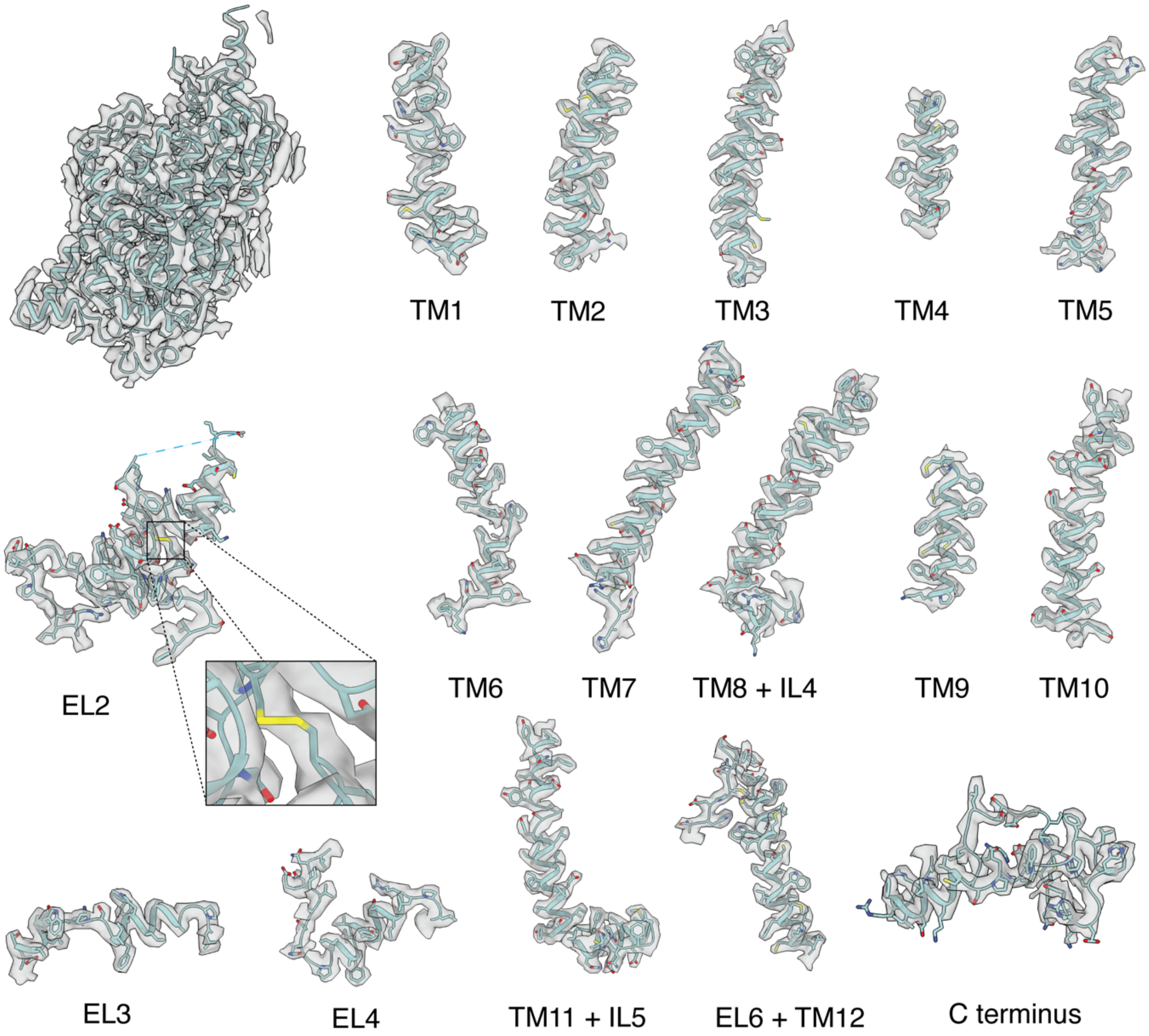

**Supplementary Fig. 10| Cryo-EM density of RPI-GLYT2-82-bound hGlyT2$^{\Delta 185}$.** The overall structure of RPI-GLYT2-82-bound hGlyT2$^{\Delta 185}$ (top left, PDB ID 9HUF) and magnified density views of transmembrane helices (contour level = 0.038 in ChimeraX), EL2 (with the conserved NSS disulfide bridge), as well as EL3, EL4 and the C terminus are shown.

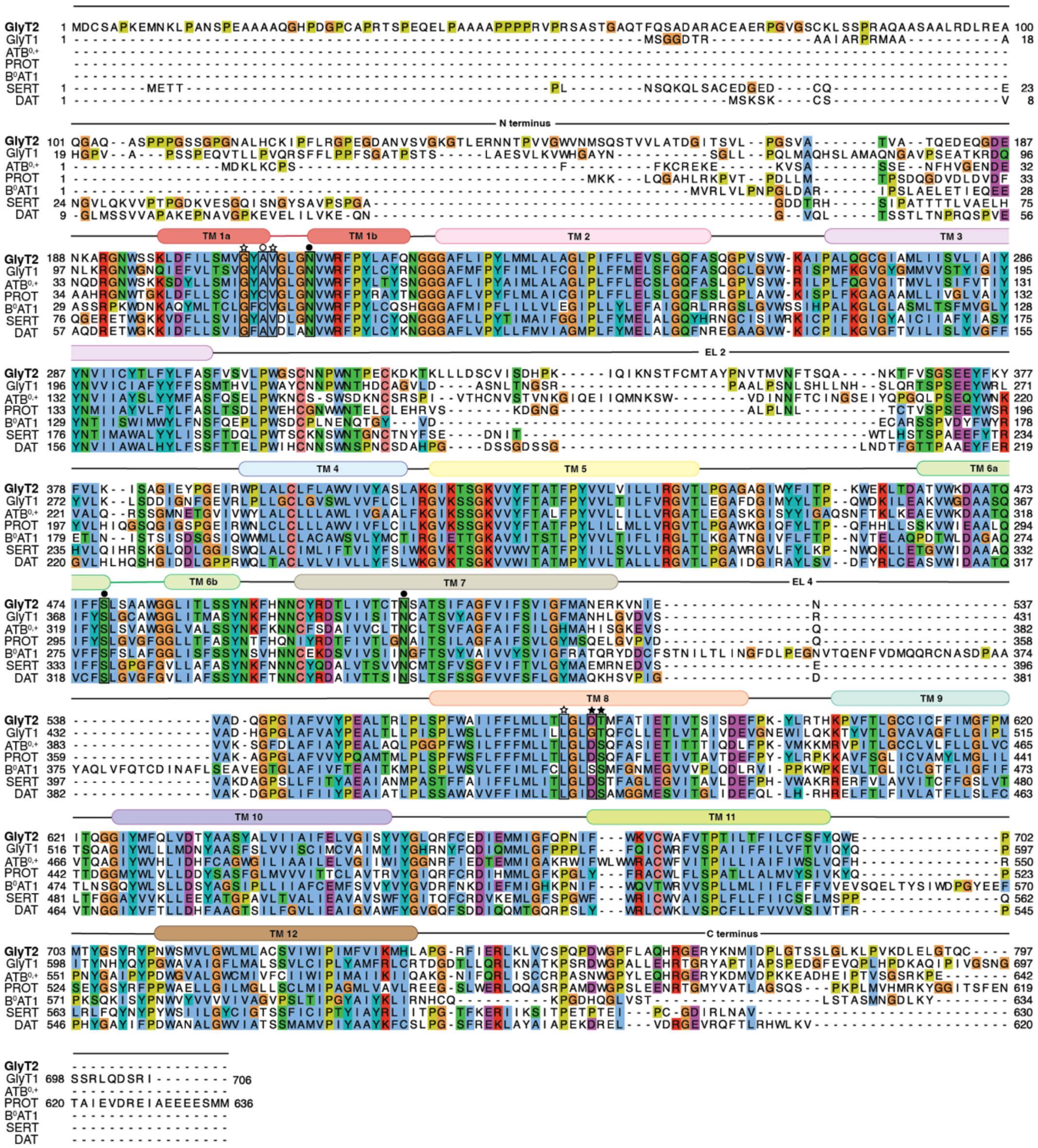

**Supplementary Fig. 11| Sequence alignment of selected SLC6 members.** ClustalW alignment of hGlyT2 (UniProt Q9Y345), hGlyT1 (UniProt P48067), hATB$^{0,+}$ (UniProt Q9UN76), hPROT (UniProt Q99884), hB$^{0}$AT1 (UniProt Q695T7), hSERT (UniProt P31645) and hDAT (UniProt Q01959) were aligned. Transmembrane (TM) helices and extracellular loops (EL) 2 and 4 are indicated above the alignment. Indicated in black boxes are the residues contributing to sodium binding in the conserved Na1 (circles) and Na2 (stars) sites. Side chain interactions are shown with a filled circle/star, and backbone interactions are shown with an empty circle/star. Colouring was generated using Jalview 2.11.4.1.

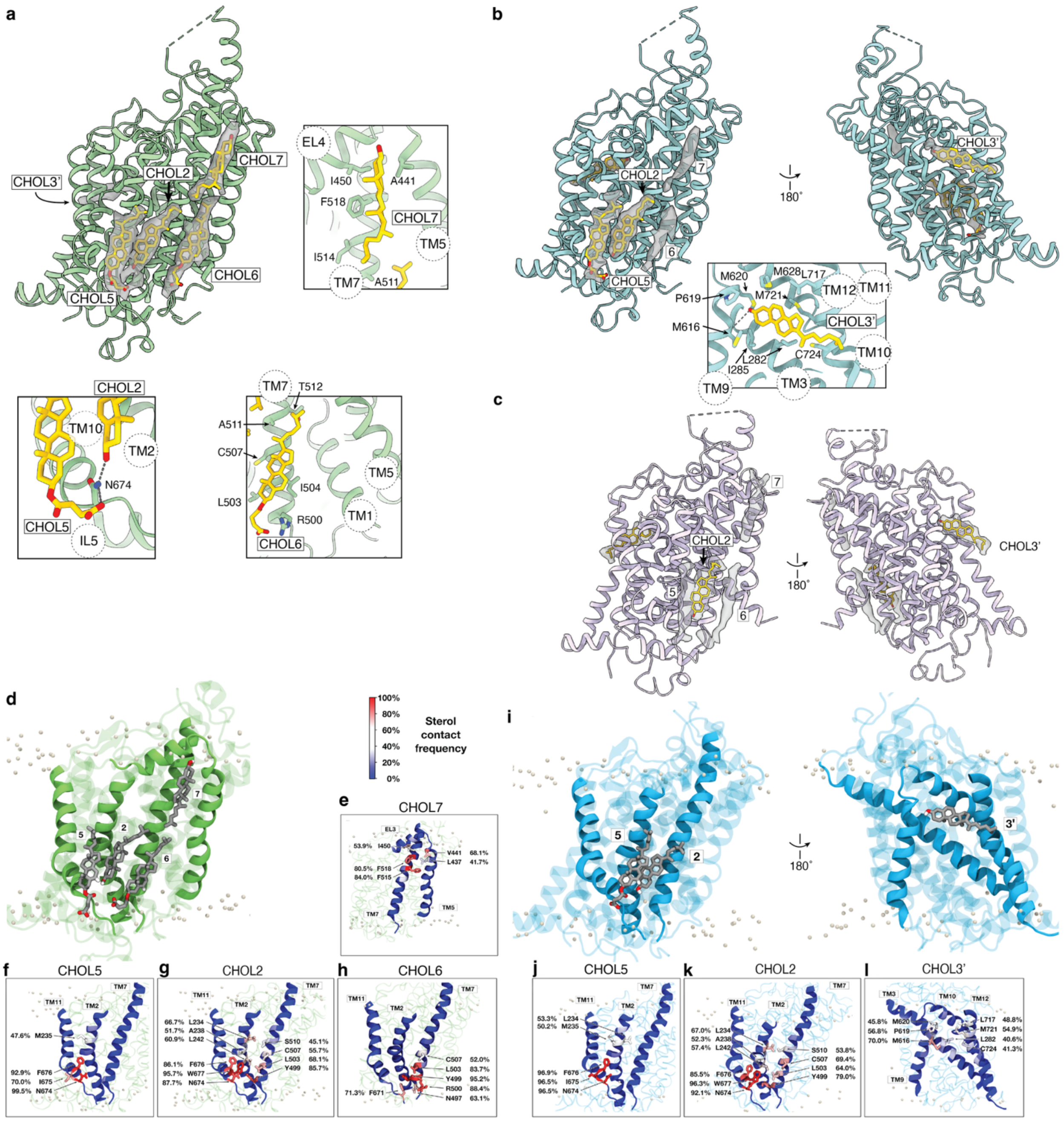

**Supplementary Fig. 12| Novel and conserved sterols bound to GlyT2. a** Two cholesterol molecules (CHOL2 and CHOL7) and two CHS molecules (CHOL5 and CHOL6) were modelled (yellow sticks) into nonprotein densities (grey) in ORG25543-bound hGlyT2$^{\Delta185}$ (map contour level = 0.025 in ChimeraX). The CHS molecule at site 6 is positioned adjacent to the conserved CHOL1 but occupies a distinct location and does not interact with TM5. **b** Sterols were modelled into CHOL2 (cholesterol), CHOL5 (CHS) and CHOL3' (cholesterol) (yellow sticks) sites of RPI-GLYT2-82-bound hGlyT2$^{\Delta185}$. Densities for sites 6 and 7 are shown but the corresponding sterols were not modelled in this structure (map contour level = 0.038 in ChimeraX). Hydrogen bonds are shown as grey dashed lines. **c** Sterols were modelled into CHOL2 (cholesterol) and CHOL3' (cholesterol) in substrate-free hGlyT2$^{\Delta185}$. Densities are also shown for sites 5, 6 and 7 but the corresponding sterols were not modelled in the structure (map contour level = 0.04 in ChimeraX).

**d-l** MD simulations were conducted with sterols placed in the binding sites identified in the cryo-EM structures. **d-h** Frequency of contact between GlyT2 residues and sterols from ORG25543-bound hGlyT2$^{\Delta185}$ cryo-EM sites. **d** All four sterols remain in their cryo-EM sites for the majority of the 3000 ns combined replicate simulation, as shown by high frequency contacts with residues identified in the ORG25543-bound hGlyT2$^{\Delta185}$ cryo-EM structure. **e-h** CHOL5 (CHS), CHOL2 (cholesterol) and CHOL6 (CHS) remain in close contact with each other through the simulation, with the highest frequency sterol/GlyT2 contacts occurring with polar and aromatic residues adjacent to the intracellular membrane interface. **i-l** Frequency of contact between GlyT2 residues and sterols from RPI-GLYT2-82-bound hGlyT2$^{\Delta185}$ cryo-EM sites. **i** MD simulations were conducted with sterols placed in the binding sites identified in the RPI-GLYT2-82-bound hGlyT2$^{\Delta185}$ cryo-EM structure. **j,k** CHOL2 (cholesterol) and CHOL5 (CHS) remain in their cryo-EM sites for the majority of the 1500 ns combined replicate simulation, demonstrated by high frequency contacts with residues identified in the cryo-EM structure. Again, the highest frequency sterol/GlyT2 contacts occurring with polar and aromatic residues adjacent to the intracellular membrane interface. **l** Cholesterol remains in contact with the CHOL3' site across the three replicates but does not remain bound in original pose, instead sampling conformations in the vicinity of the CHOL3' site. Residues are coloured by frequency of contact, defined as any residue atom lying within 4 Å of any sterol atom.

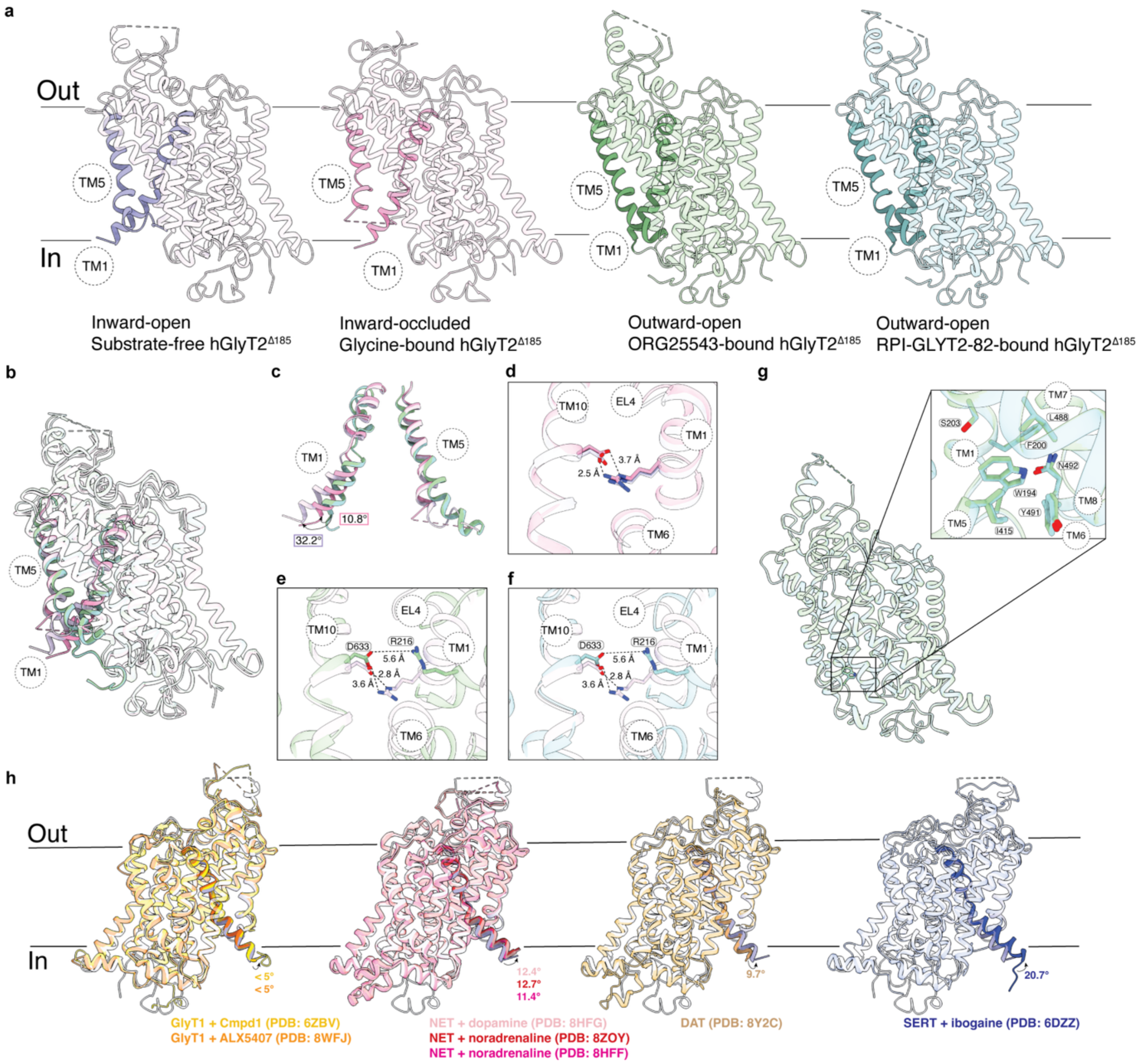

**Supplementary Fig. 13| Inward-open, inward-occluded and outward-open conformational states of hGlyT2$^{\Delta185}$. a** Substrate-free hGlyT2$^{\Delta185}$(purple) adopts an inward-open conformation, glycine-bound hGlyT2$^{\Delta185}$ (pink) adopts an inward-occluded conformation and ORG25543-bound hGlyT2$^{\Delta185}$ (green) and RPI-GLYT2-82-bound hGlyT2$^{\Delta185}$ (blue) adopt outward-open conformations. **b** Overlay of the four hGlyT2$^{\Delta185}$ models with TM1 and TM5 highlighted. c In the inward-open conformation, TM1 is shifted by 32.2° away from the core of the transporter compared to the outward-open conformation, and in the inward-occluded conformation it is shifted 10.8° away, as measured by the Cα-Cα angle of L211 (as one fixed point) and S196 on each structure (as the two moving points). The intracellular portion of TM5 is unwound in the inward-open and inward-occluded conformations but α-helical in the outward-open conformations. **d-f** The extracellular gate forming residues, R216 and D633, and the side-chain distances are shown as a comparison of **d** glycine-bound hGlyT2$^{\Delta185}$ (pink), **e** ORG25543-bound hGlyT2$^{\Delta185}$ structure (green) and **f** RPI-GLYT2-82-bound hGlyT2$^{\Delta185}$ structure (blue), with the substrate-free hGlyT2$^{\Delta185}$ structure (purple). The extracellular gate is open in the outward-open inhibitor-bound hGlyT2$^{\Delta185}$ structures and closed in the inward-open substrate-free hGlyT2$^{\Delta185}$ structure and the inward-occluded glycine-bound hGlyT2$^{\Delta185}$ structure, as indicated by the R216 (TM1b) and D633 (TM10) distances. **g** W194 (TM1) buries into a hydrophobic patch in the outward-open inhibitor-bound GlyT2$^{\Delta185}$ structures where the intracellular pathway is closed. **h** Comparison of substrate-free GlyT2$^{\Delta185}$ structure (purple) with several inward-open SLC6 transporter structures. TM1 is highlighted with the angle between substrate-free GlyT2$^{\Delta185}$ and each SLC6 transporter measured between the Cα of the conserved L211 (GlyT2 numbering) as one fixed point and the first Cα of TM1 of each transporter (S196 in GlyT2) as the moving points. The moving points were selected as follows: E54 in GlyT1 (PDB ID 8WFJ), N105 in GlyT1 (PDB ID 6ZBV), G60 in NET (PDB IDs 8FHG, 8FHH and 8ZOY), K66 in DAT (PDB ID 8Y2C) and T81 in SERT (PDB ID 6DZZ).

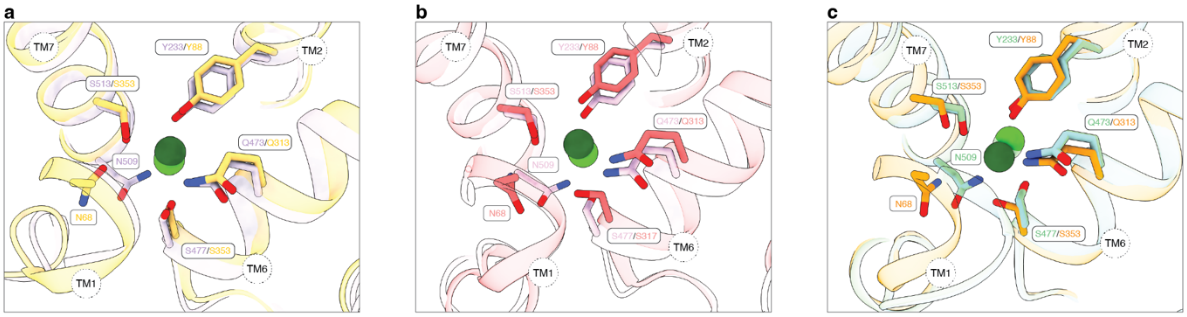

**Supplementary Fig. 14| Chloride coordination in GlyT2 and GlyT1. a** Substrate-free hGlyT2$^{\Delta185}$(purple) and iclepertin-bound GlyT1 (PDB ID 9J8D) (yellow) in the inward-open state, **b** Glycine-bound hGlyT2$^{\Delta185}$ (pink) and glycine bound GlyT1 (PDB ID 8WFI) (red) in an inward-occluded state, and **c** ORG25543-bound hGlyT2$^{\Delta185}$ (green), RPI-GLYT2-82-bound hGlyT2$^{\Delta185}$ (blue) and SSR504734-bound GlyT1 (PDB ID 8WFK) in outward-open conformations are overlayed, showing similar coordination of chloride ion across the three conformations. GlyT1 chloride ions are shown in dark green and GlyT2 chloride ions are shown in lighter green. In **c**, the chloride ion of RPI-GLYT2-82-bound hGlyT2$^{\Delta185}$ is shown in the lightest green. Transmembrane helices are labelled.

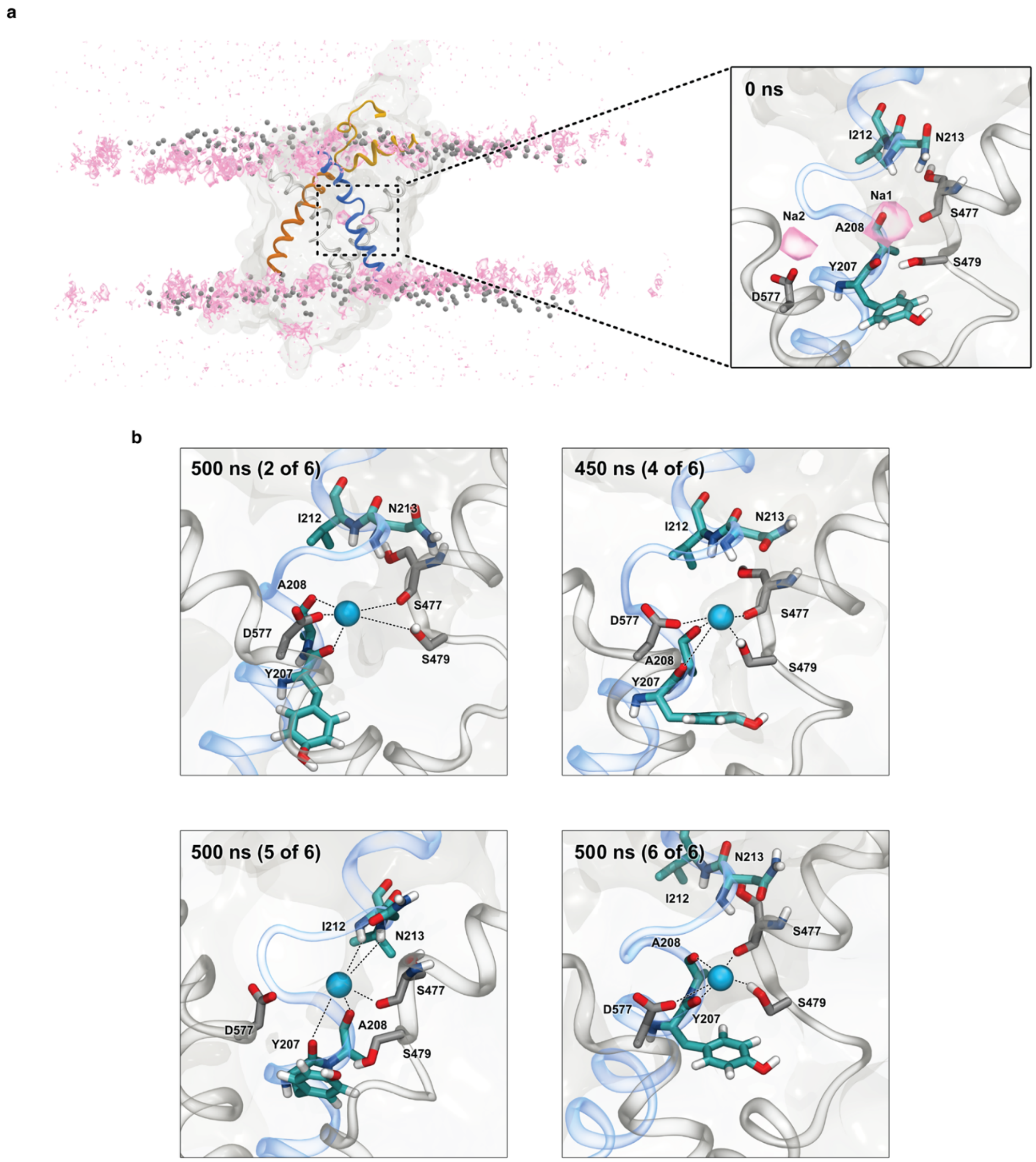

**Supplementary Fig. 15| Ion binding sites in inhibitor-bound GlyT2 determined by MD simulations and cryo-EM. a** $Na^+$ density across the 3000 ns combined replicate simulations, plotted in VMD with a density threshold of 0.05, shows $Na^+$ density is primarily clustered at the membrane/solvent interface in ORG25543-bound GlyT2$^{\Delta185}$. However, there are two hotspots of $Na^+$ density corresponding to the Na1 and Na2 binding sites. **b** When the ORG25543-bound GlyT2$^{\Delta185}$ complex was modelled with water in the sodium binding sites, extracellular $Na^+$ spontaneously bound to the Na1 site in four of the six replicates and remained loosely bound for 50% of the combined replicate MD simulations. While sodium was mobile within the site, it was primarily coordinated by the backbone carbonyl groups of Y207, A208 and S477, the sidechain hydroxyl of S479, and the sidechain carboxylate of D577, which rotated away from the vacant Na2 site to contact the sodium in the Na1 site.

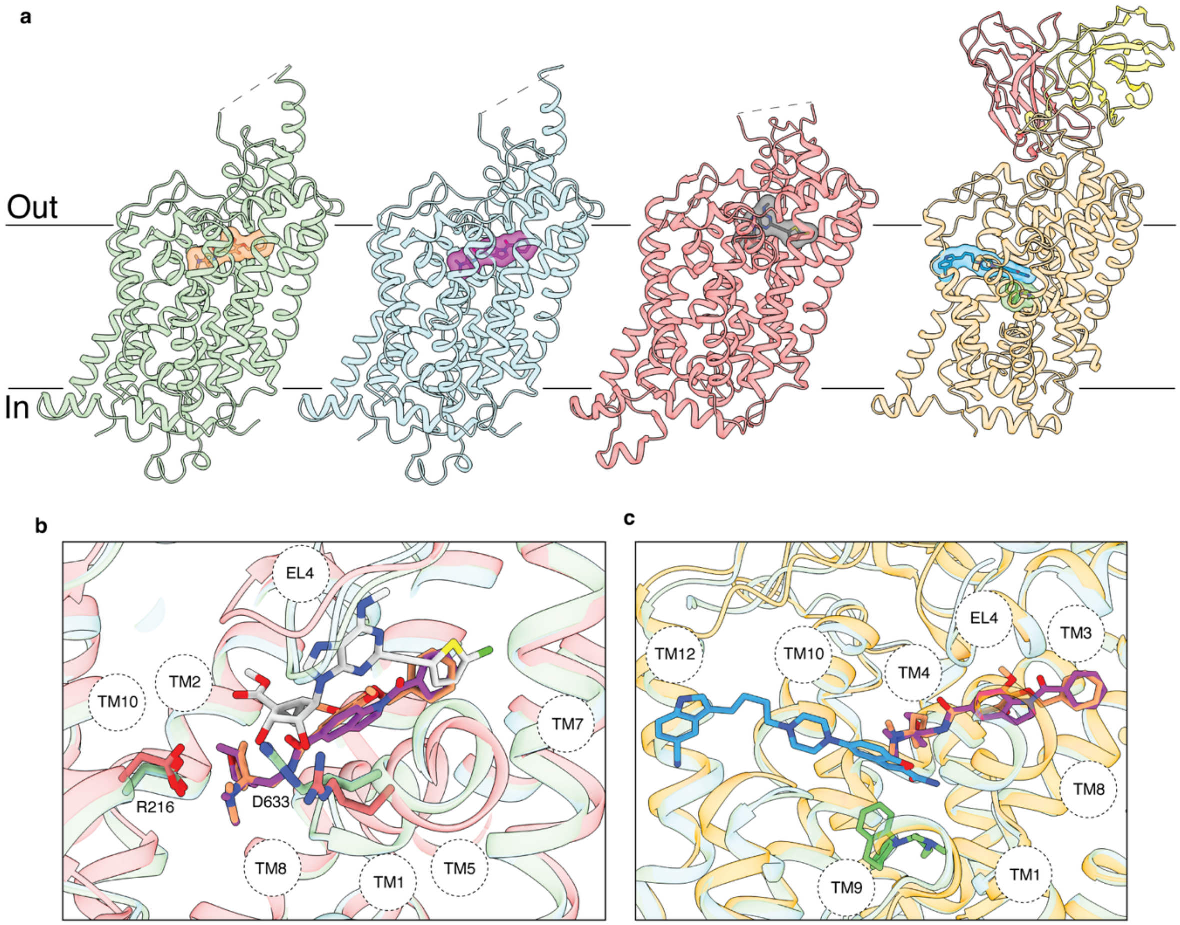

**Supplementary Fig. 16| The comparison of allosteric site in hGlyT2, hDAT and hSERT. a** Comparison of ORG25543-bound hGlyT2$^{\Delta185}$ (green) and RPI-GLYT2-82-bound hGlyT2$^{\Delta185}$ to MRS7292-bound DAT (red, PDB ID 8VBY) and vilazodone and imipramine-bound SERT (yellow, PDB ID 7LWD). **b** Overlay of allosteric binding pocket of ORG25543 (orange), RPI-GLYT2-82 (purple) in GlyT2 and MRS7292 (grey) in DAT. The residues forming the extracellular molecular gate, R216 and D633 (in GlyT2), are shown as sticks. **c** Overlay of allosteric binding pocket of ORG25543 (orange), RPI-GLYT2-82 (purple) in GlyT2 and vilazodone (blue) and imipramine (green) in SERT.

a

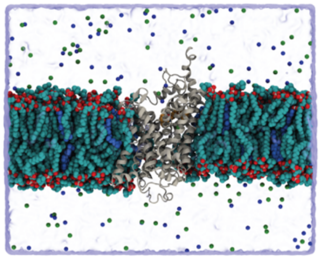

b

| Simulation System | GlyT2 RMSD Backbone Cα | Inhibitor RMSD Heavy Atom |
|---|---|---|
| GlyT2$^{\Delta185}$ – ORG25543 | 2.83 ± 0.44 Å | 1.72 ± 0.45 Å |
| GlyT2$^{\Delta185}$ – RPI-GLYT2-82 | 2.23 ± 0.39 Å | 1.20 ± 0.41 Å |

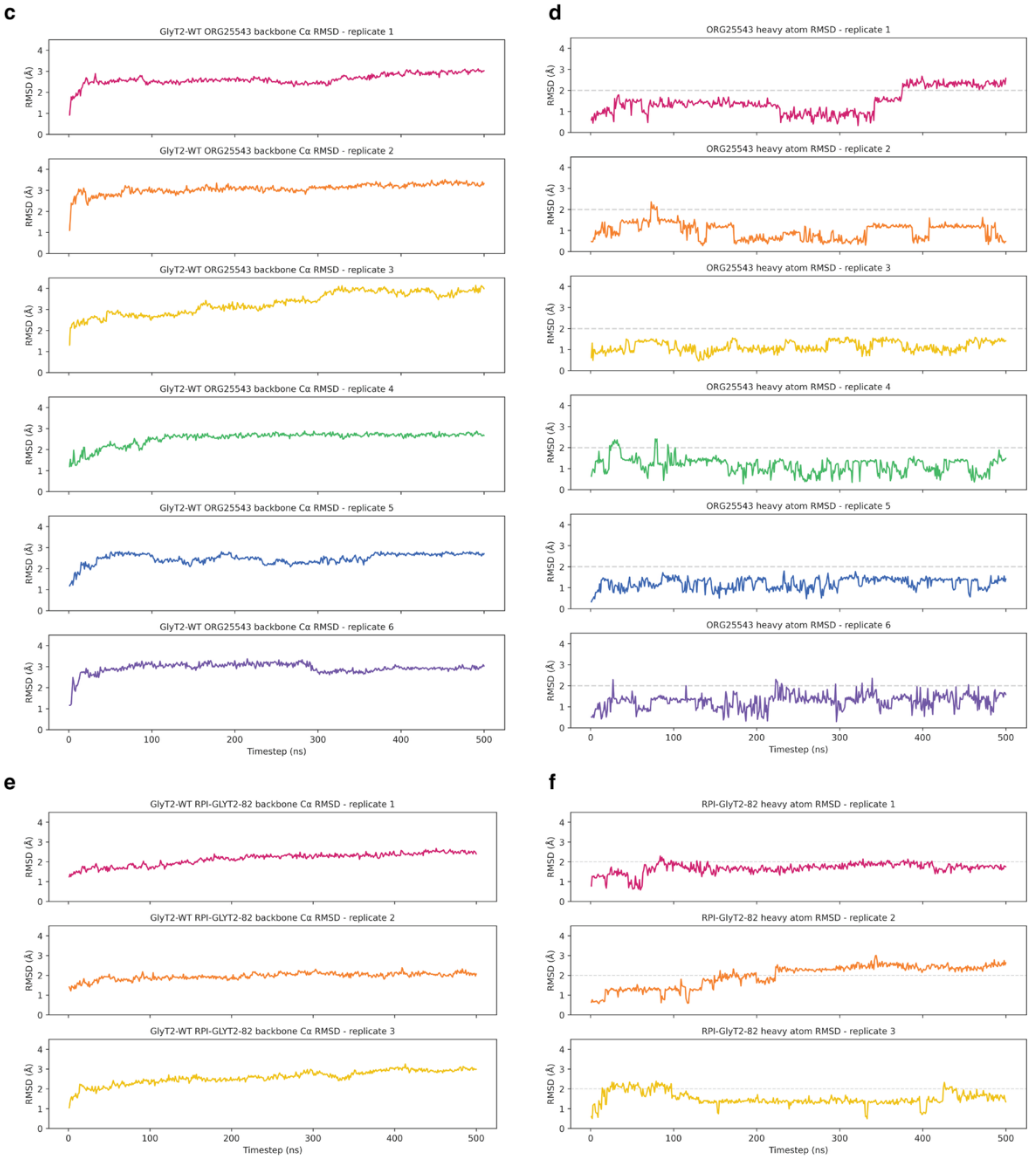

**Supplementary Fig. 17| Root mean squared deviation of the membrane embedded GlyT2 simulations. a** Snapshot of the starting conformation of the solvated, membrane embedded GlyT2 system used to initiate MD simulations. **b** Summary overview of the GlyT2 backbone RMSD, and heavy atom RMSD of ORG25543 and RPI-GLYT2-82 calculated across the combined MD trajectories for each system with respect to the initial GlyT2 conformation. **c-f** Time dependent **c**

GlyT2 backbone RMSD and **d** heavy atom RMSD of ORG25543 for each of the six 500 ns replicate simulations of GlyT2 with ORG25543. Time dependent **e** GlyT2 backbone RMSD and **f** heavy atom RMSD of RPI-GLYT2-82 for each of the three 500 ns replicate simulations of GlyT2 with RPI-GlyT2-82.

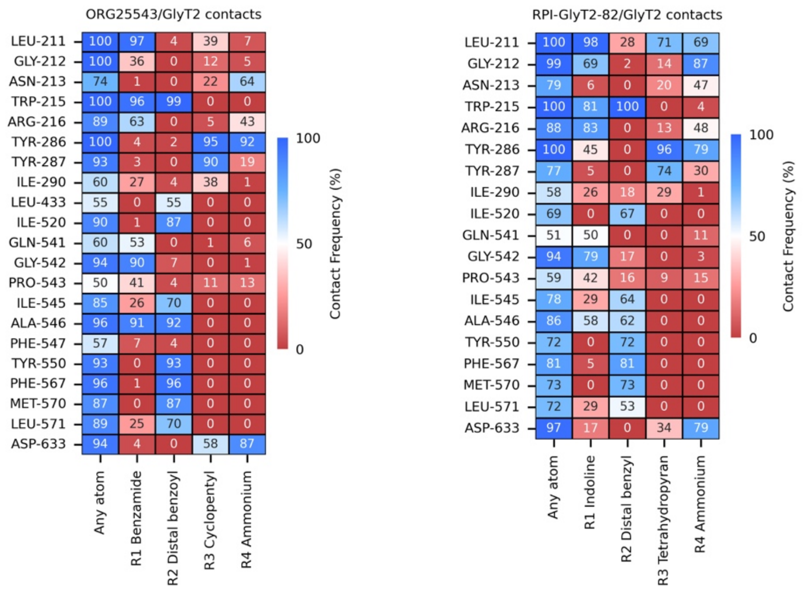

**Supplementary Fig. 18| Inhibitors-GlyT2 contact frequencies.** Heatmaps showing the contact frequencies between ORG25543 functional groups and GlyT2 (left), and between RPI-GLYT2-82 functional groups and GlyT2 (right), in MD simulations of the ORG25543-bound and RPI-GLYT2-82-bound GlyT2 complexes, respectively.

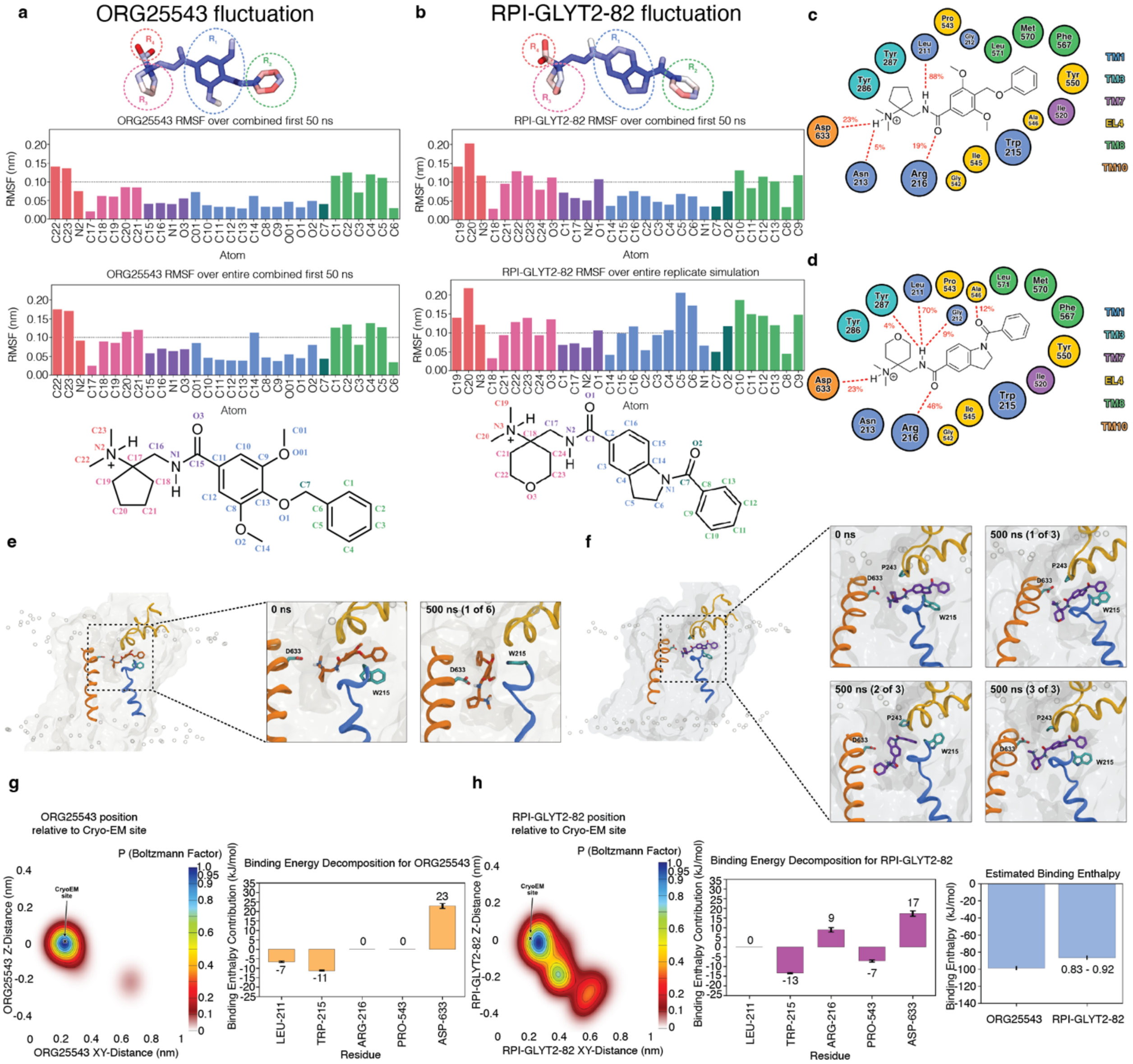

**Supplementary Fig. 19| MD simulations and root mean squared fluctuation (RMSF) of ORG25543 and RPI-GlyT2-82. a** ORG25543 and **b** RPI-GLYT2-82 calculated across the combined replicate simulations for each system. Top panel: licorice structure of ORG25543 coloured by RMSF beta factor (scale: blue = lowest RMSF, red = highest RMSF) and labelled by functional moiety $R_1$, $R_2$, $R_3$ and $R_4$. Middle panels: bar graphs of the per atom RMFS over the first 50 ns of the combined replicate simulation (top) and over the entire trajectory of the combined replicate simulations (bottom). Bottom panel: structural legend of the RMFS plots for **a** ORG25543 and **b** RPI-GlyT2-82 showing the corresponding atom names and colour legend. Pictorial overview of GlyT2 residues contributing to the binding of **c** ORG25543 and **d** RPI-GLYT2-82, coloured by TM helix, and frequency of hydrogen bonding interactions (red lines). **e** Side view of GlyT2 (transparent grey) showing the position of ORG25543 (orange) and its interaction with D633 from TM1 (blue), W215 from TM10 (orange) and EL4 (yellow), showing close-ups at the start of the simulation (0 ns) and after 500 ns of simulation in replica 1 of 6. **f** Side view of GlyT2 (transparent grey) showing the position of RPI-GLYT2-82 (purple) and its interaction with W215 from TM1 (blue), D633 from TM10 (orange) and P243 from EL4 (yellow), showing close-ups at the start of the simulation (0 ns) and after 500 ns of simulation in replica 3 of 3. **g** Heat map of the relative position of ORG25543 across the 1500 ns of unbiased replicate MD simulations, where each point on the plot corresponds to the probability of being in that position relative to the most probable state (computed as a Boltzmann factor) and MM-PBSA per residue binding enthalpy decomposition of key residues from the ORG25543 binding site. The enthalpic contribution of the interaction between D633 and ORG25543 was less favourable than that calculated for the D633/PBSA water interaction in simulations at 310 K. **h** Heat map of the relative position of RPI-GLYT2-82 across the 1500 ns of unbiased replicate MD simulations, where each point on the plot corresponds to the probability of being in that position relative to the most probable state (computed as a Boltzmann factor). MM-PBSA per residue binding enthalpy decomposition of key residues in the RPI-GLYT2-82 binding site. The enthalpic contribution of the interaction between D633 and RPI-GLYT2-82 was less favourable than that calculated for the D633/PBSA water interaction in simulations at 310 K. MM-PBSA estimated relative binding enthalpy of ORG25543 and RPI-GLYT2-82 over the first 50 ns of each replicate simulation, with estimated fold difference in binding enthalpy labelled (95% CI).

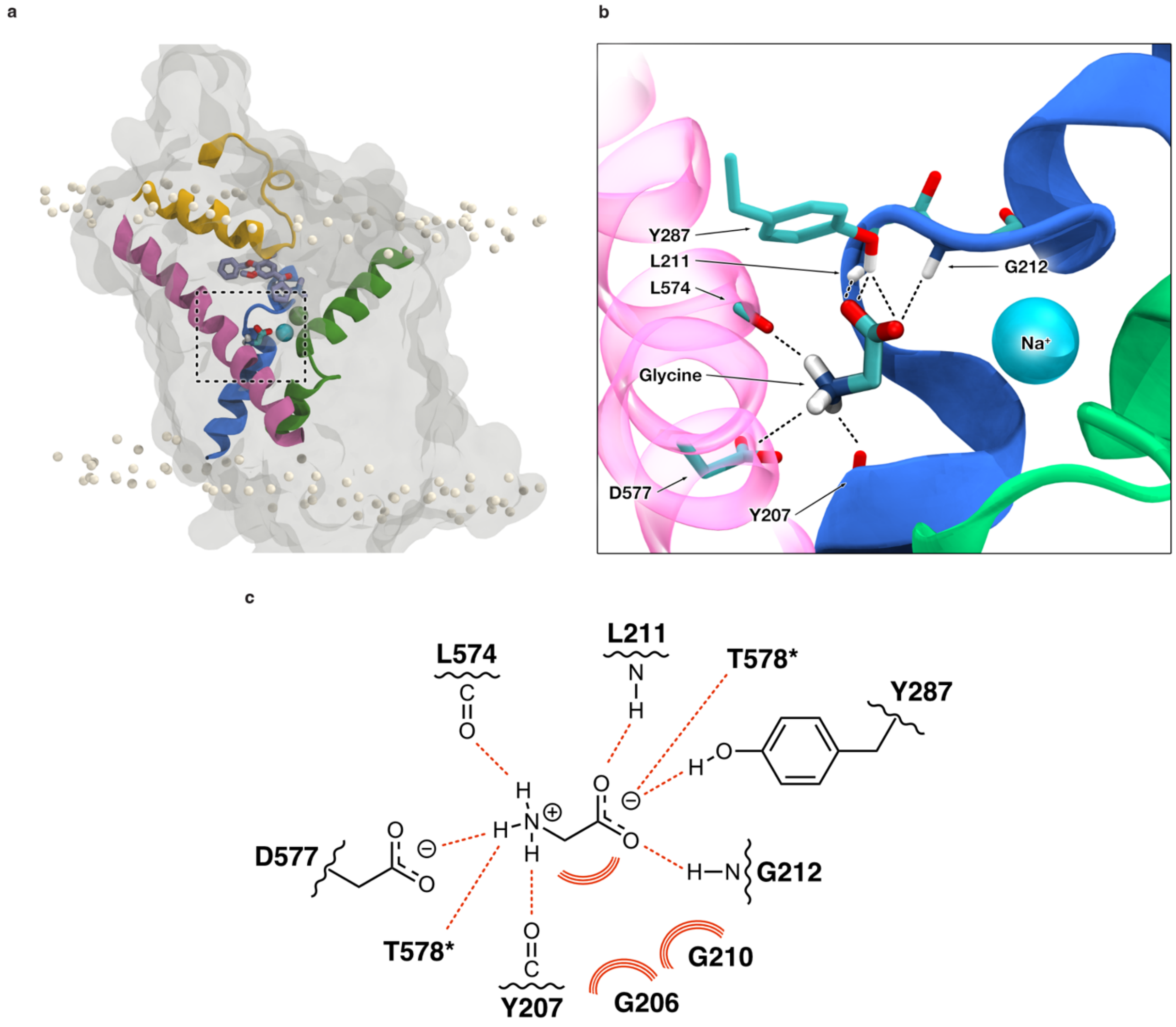

**Supplementary Fig. 20| Glycine docked into the substrate binding cavity of the ORG25543-GlyT2 complex. a** Glycine (cyan sticks) binds to the vacant Na2 binding site, located between TM1 (blue), TM6 (green), and TM8 (pink). ORG25543 (purple sticks) remains in the cryo-EM binding site, below EL4 (yellow). Sodium (cyan) and chloride (green) ions remain in their respective binding sites. **b** Bound glycine is coordinated by the backbone amine groups of L211 and G212, the backbone carbonyl groups of Y207 and L574, and the sidechains of Y287, D577, and T578 (not shown). **c** A schematic showing coordination of glycine by GlyT2 residues, highlighting direct interactions and frequent contacts. Coordination by T578 was frequent, with glycine/T578 contact present in 98% of frames, but occurred through a variety of modes, with T578 coordinating either the ammonium or carboxylate groups of glycine. Both possible coordination modes are shown, marked with an asterisk.

**Supplementary Table 1| RPI-GLYT2-82 displays non-competitive behavior.**

| [RPI-GLYT2-82] (nM) | Glycine $EC_{50}$ ± SEM (μM) | $I/I_{300}$ ± SEM |
|---|---|---|
| 0 | 18.0 ± 1.2 | 1.07 ± 0.01 |
| 200 | 27.9 ± 3.3$^{ns}$ | 0.98 ± 0.026$^{ns}$ |
| 600 | 26.7 ± 6.1$^{ns}$ | 0.76 ± 0.039$^{****}$ |
| 1800 | 24.3 ± 6.3$^{ns}$ | 0.39 ± 0.022$^{****}$ |

$EC_{50}$ Statistics: Brown-Forsythe ANOVA with Dunnett's T3 multiple comparisons (ns $p > 0.05$). $I/I_{300}$ Statistics: Ordinary one-way ANOVA with Dunnet's multiple comparisons (ns $p > 0.05$, **** $p < 0.0001$). Significance indicated for comparisons to control (0 nM RPI-GLYT2-82).

**Supplementary Table 2| Cryo-EM data collection, refinement and validation statistics.**

| | ORG25543-bound GlyT2$^{\Delta185}$ (EMDB-52409) (PDB 9HUE) | RPI-GLYT2-82-bound GlyT2$^{\Delta185}$ (EMDB-52410) (PDB 9HUF) | Glycine-bound GlyT2$^{\Delta185}$ (EMDB-53509) (PDB 9R1H) | Substrate-free GlyT2$^{\Delta185}$ (EMDB-52411) (PDB 9HUG) |
|---|---|---|---|---|
| **Data collection and processing** | | | | |
| Magnification | 165,000 × | 165,000 × | 165,000 × | 165,000 × |
| Voltage (kV) | 300 | 300 | 300 | 300 |
| Electron exposure (e$^-$/Å$^2$) | 60 | 60 | 60 | 60 |
| Defocus range (μm) | –0.6 to –1.8 | –0.6 to –1.8 | –0.6 to –1.8 | –0.6 to –1.8 |
| Pixel size (Å) | 0.729 | 0.728 | 0.728 | 0.728 |
| Symmetry imposed | C1 | C1 | C1 | C1 |
| Initial particle images (no.) | 2,807,000 | 6,700,966 | 3,363,288 | 6,458,661 |
| Final particle images (no.) | 283,838 | 201,519 | 162,255 | 180,698 |
| Map resolution (Å) | 2.49 | 2.79 | 3.02 | 2.97 |
| FSC threshold | 0.143 | 0.143 | 0.143 | 0.143 |
| Map resolution range (Å) | 1.75 – 2.75 | 2.00 – 3.00 | 2.50 – 3.00 | 2.00 – 3.50 |
| **Refinement** | | | | |
| Initial model used (PDB code) | AF-Q9Y345-F1 | 9HUE | AF-Q9Y345-F1 | AF-Q9Y345-F1 |
| Model resolution (Å) | 2.4 | 2.7 | 3.0 | 2.9 |
| FSC threshold | 0.143 | 0.143 | 0.143 | 0.143 |
| Map sharpening *B* factor (Å$^2$) | | | | |
| Model composition | | | | |
| Non-hydrogen atoms | 4703 | 4664 | 4361 | 4491 |
| Protein residues | 571 | 571 | 548 | 558 |
| Ligands | 7 | 6 | 3 | 3 |
| Waters | 3 | 3 | 2 | 0 |
| *B* factors (Å$^2$) | | | | |
| Protein | 56.26 | 63.96 | 69.39 | 65.08 |
| Ligand | 75.19 | 76.04 | 65.16 | 91.03 |
| R.m.s. deviations | | | | |
| Bond lengths (Å) | 0.004 (0) | 0.004 (0) | 0.003 (0) | 0.003 (0) |
| Bond angles (°) | 0.781 (2) | 0.720 (2) | 0.730 (0) | 0.662 (1) |
| Validation | | | | |
| MolProbity score | 1.16 | 1.14 | 1.43 | 1.16 |
| Clashscore | 3.71 | 3.46 | 6.22 | 3.67 |
| Poor rotamers (%) | 0.61 | 0.62 | 1.29 | 0.42 |
| CaBLAM outliers (%) | 0.89 | 0.71 | 0.93 | 0.55 |
| Ramachandran plot | | | | |
| Favored (%) | 98.59 | 98.41 | 98.15 | 98.74 |
| Allowed (%) | 1.41 | 1.59 | 1.85 | 1.26 |
| Disallowed (%) | 0.00 | 0.00 | 0.00 | 0.00 |

**Supplementary Table 3| Per residue contact frequency between $Na^+$ ions and GlyT2 residues in MD simulations of the ORG25543-bound GlyT2 complex.**

| $Na^+$ / GlyT2 contact frequency* | | | |
|---|---|---|---|
| **Residue** | **Binding site** | **Backbone $Na^+$ contact** | **Sidechain $Na^+$ contact** |
| G206 | Na2 | 14.46 % | *n/a* |
| Y207 | Na1 | 27.09 % | 5.63 % |
| A208 | Na1 | 32.19 % | 0.03 % |
| V209 | Na2 | 32.06 % | 0.07 % |
| G212 | Na1 | 15.26 % | *n/a* |
| N213 | Na1 | 18.59 % | 0.07 % |
| S477 | Na1 | 28.82 % | 0.53 % |
| S479 | Na1 | 9.30 % | 14.53% |
| L574 | Na2 | 13.93 % | 1.13 % |
| D577 | Na2 | 0.27% | 18.36% |

* A contact is defined as any atom belonging to the residue backbone or sidechain being within 4 Å of any $Na^+$ ion. Contacts are shown for any residue in contact with $Na^+$ for at least 10 % of the 3000 ns combined replicate simulation.

**Supplementary Table 4| Per residue contact frequency between ORG25543 and RPI-GLYT2-82 functional groups and GlyT2 in MD simulations of the ORG25543-bound and RPI-GLYT2-82-bound GlyT2 complexes.**

| ORG25543 / GlyT2 contact frequency[a] | | | | | |
|---|---|---|---|---|---|
| | **Any atom** | **$R_1$ Benzamide** | **$R_2$ Distal benzoyl** | **$R_3$ Cyclopentyl** | **$R_4$ Ammonium** |
| LEU 211 | **100 %** | **96.6 %** | 3.8 % | 39 % | 7 % |
| GLY 212 | **99.5 %** | 36.3 % | *n.c.* | 12 % | 5.2 % |
| ASN 213 | 74.1 % | 0.8 % | *n.c.* | 22.5 % | 63.9 % |
| TRP 215 | **100 %** | **95.8 %** | **99.2 %** | *n.c.* | *n.c.* |
| ARG 216 | **89.4 %** | 63.1 % | *n.c.* | 5.2 % | 42.6 % |
| TYR 286 | **99.8 %** | 4.2 % | 2.3 % | **95.3 %** | **91.6 %** |
| TYR 287 | **92.7 %** | 3.3 % | *n.c.* | **89.8 %** | 18.6 % |
| ILE 290 | 60.5 % | 26.7 % | 3.8 % | 37.7 % | 1.4 % |
| LEU 433 | 55.4 % | *n.c.* | 55.4 % | *n.c.* | *n.c.* |
| ILE 520 | **89.7 %** | 0.8 % | **87.2 %** | *n.c.* | *n.c.* |
| GLN 541 | 60 % | 53.4 % | *n.c.* | 0.7 % | 6.2 % |
| GLY 542 | **94.4 %** | **90.3 %** | 6.9 % | 0.2 % | 0.8 % |
| PRO 543 | 50.5 % | 40.6 % | 3.7 % | 11 % | 13 % |
| ILE 545 | **85.1 %** | 26.5 % | 69.9 % | *n.c.* | *n.c.* |
| ALA 546 | **95.7 %** | **90.7 %** | **91.7 %** | *n.c.* | *n.c.* |
| PHE 547 | 57.2 % | 6.8 % | 3.9 % | 0.1 % | *n.c.* |
| TYR 550 | **92.9 %** | 0 % | **92.9 %** | *n.c.* | *n.c.* |
| PHE 567 | **95.7 %** | 0.7 % | **95.7 %** | *n.c.* | *n.c.* |
| MET 570 | **86.9 %** | *n.c.* | **86.9 %** | *n.c.* | *n.c.* |
| LEU 571 | **89.3 %** | 25.4 % | 69.5 % | *n.c.* | *n.c.* |
| ASP 633 | **93.6 %** | 4.2 % | *n.c.* | 58.3 % | **86.9 %** |
| **RPI-GLYT2-82 / GlyT2 contact frequency[b]** | | | | | |
| | **Any atom** | **$R_1$ Indoline** | **$R_2$ Distal benzyl** | **$R_3$ Tetrahydropyran** | **$R_4$ Ammonium** |
| LEU 211 | **99.9 %** | **98.5 %** | 28.5 % | 71.4 % | 69.4 % |
| GLY 212 | **98.6 %** | 69.4 % | 1.8 % | 13.9 % | **87.3 %** |
| ASN 213 | 79.2 % | 5.8 % | *n.c.* | 20.3 % | 46.8 % |
| TRP 215 | **99.5 %** | **81.2 %** | **99.5 %** | *n.c.* | 4 % |
| ARG 216 | **88.3 %** | **82.7 %** | 0.1 % | 12.7 % | 47.8 % |
| TYR 286 | **100 %** | 44.6 % | *n.c.* | **95.5 %** | 79.4 % |
| TYR 287 | 76.8 % | 5.4 % | 0.1 % | 73.6 % | 29.5 % |
| ILE 290 | 58.3 % | 25.9 % | 18.3 % | 29.2 % | 1.1 % |
| ILE 520 | 68.8 % | 0.5 % | 67.4 % | *n.c.* | *n.c.* |
| GLN 541 | 51.1 % | 50.4 % | 0.3 % | 0.1 % | 11.1 % |
| GLY 542 | **93.5 %** | 79 % | 16.7 % | 0.5 % | 2.8 % |
| PRO 543 | 59 % | 42.3 % | 15.5 % | 8.8 % | 14.7 % |
| ILE 545 | 77.7 % | 29 % | 63.6 % | *n.c.* | *n.c.* |

| | | | | | |
|---|---|---|---|---|---|
| ALA 546 | 86.3 % | 57.6 % | 62 % | *n.c.* | *n.c.* |
| TYR 550 | 71.8 % | 0.1 % | 71.8 % | *n.c.* | *n.c.* |
| PHE 567 | **81.2 %** | 5.2 % | **81.2 %** | *n.c.* | *n.c.* |
| MET 570 | 73 % | 0.3 % | 73 % | *n.c.* | *n.c.* |
| LEU 571 | 72.2 % | 28.7 % | 52.6 % | *n.c.* | *n.c.* |
| ASP 633 | **97.1 %** | 16.9 % | *n.c.* | 34.5 % | 79 % |

[a]A contact is defined as any atom of the ORG25543 functional group being within 4 Å of any atom of a GlyT2 residue. *n.c.* indicates contact within 4 Å was not observed. Contacts are shown for any residue in contact with ORG25543 for at least 50 % of the 3000 ns combined replicate simulations.

[b]A contact is defined as any atom of the RPI-GLYT2-82 functional group being within 4 Å of any atom of a GlyT2 residue. *n.c.* indicates contact within 4 Å was not observed. Contacts are shown for any residue in contact with RPI-GLYT2-82 for at least 50 % of the 1500 ns combined replicate simulations.

**Supplementary Table 5| Frequency of contact between GlyT2 residues and zwitterionic glycine during glycine docking simulations.**

| | | GlyT2 + ORG25543 + glycine | GlyT2 + RPI-GLYT2-82 + glycine |
|---|---|---|---|
| | | Glycine contact frequency (%) | |
| 206 | GLY | 98 | 14 |
| 207 | TYR | 94 | 67 |
| 209 | VAL | 55 | 15 |
| 210 | GLY | 99 | 49 |
| 211 | LEU | 98 | 47 |
| 212 | GLY | 89 | 61 |
| 287 | TYR | 92 | 76 |
| 476 | PHE | 0 | 72 |
| 479 | SER | 10 | 68 |
| 482 | TRP | 4 | 57 |
| 574 | LEU | 93 | 3 |
| 577 | ASP | 96 | 27 |
| 578 | THR | 98 | 48 |
| 789 | NA | 45 | 48 |

Values report the percentage of simulation frames in which any atom of the GlyT2 residue was within 4 Å of any atom of the docked glycine molecule. Values are reported only for those residues in contact with glycine for more than 40% of the combined replicate simulations for at least one simulation system.

**Supplementary Table 6| Root mean squared deviation of GlyT2, substrates and inhibitors during glycine docking simulations.**

| | **GlyT2 + ORG25543 + glycine** | **GlyT2 + RPI-GLYT2-82 + glycine** |
|---|---|---|
| | **Heavy atom RMSD** | |
| GlyT2 backbone | 1.4 + 0.2 | 1.4 + 0.2 |
| Glycine | 1.8 + 0.7 | 3.4 + 1.6 |
| Inhibitor | 1.6 + 0.7 | 1.8 + 0.6 |
| Chloride | 0.9 + 0.4 | 0.8 + 0.4 |
| Sodium | 1.9 + 1.8 | 4.0 + 2.1 |

All RMSD calculations performed for heavy atoms (excluding protons), relative to initial coordinates fit to the GlyT2 backbone.

**Supplementary Table 7| *In vivo* pharmacokinetic data for RPI-GLYT2-82 following i.p. administration in mice.**

| RPI-GLYT2-82 IP PK Study and Bioanalysis in Mouse (PK Parameters)<br>Study 1[a] | | | | | | | | |
|---|---|---|---|---|---|---|---|---|
| **Route[b]** | **Dose[c] (mg/kg)** | **$t_{1/2}$ (h)[d]** | **$C_{max}$ (ng/mL)[e]** | **$T_{max}$ (h)[f]** | **$AUC_{last}$ (h*ng/mL)[g]** | **$AUC_{inf}$ (h*ng/mL)[h]** | **$V_d$ (L/kg)[i]** | **CL (mL/min/kg)[j]** |
| i.p. | 30 | 1.08 ± 0.05 | 152 ± 31 | 1.0 ± 0 | 241 ± 40 | 246 ± 41 | 192.93 ± 23.32 | 2063.72 ± 313.68 |

| RPI-GLYT2-82 IP In-life Plasma/CNS PK<br>Study 2[k] | | | | | | | |
|---|---|---|---|---|---|---|---|
| **Route[l]** | **Dose[m] (mg/kg)** | **Brain (ng/g)[n]** | **Plasma (ng/mL)[o]** | **Mouse %Brain Tissue Homogenate Binding** | **Mouse %Plasma Protein Binding** | **$C_{u,b}$[q]** | **$C_{u,p}$[r]** |
| i.p. | 50 | 550 ± 171 (30 min) | 2296.3 ± 177.2 (30 min) | 55.7% bound | 53% bound | 597 nM (30 min) | 2648 nM (30 min) |
| | | 39.0 ± 14.8 (120 min) | 132.7 ± 31.1 (120 min) | | | 42.3 nM (120 min) | 153 nM (120 min) |

Data are represented as the mean ± SD. [a]Fundamental PK parameters of RPI-GLYT2-82 after 30 mg/kg, i.p. administration. Dosing groups consisted of three drug naïve adult male ICR mice. [b]Intraperitoneal (i.p.) administration. [c] Test article was administered at the 30 mg/kg dose; test article vehicle = 5% Solutol® HS-15/95% PBS. [d]Apparent half-life of the terminal phase of elimination of compound from blood. [e]Maximum observed concentration of compound in blood. [f]Time of maximum observed concentration of compound in blood. [g]Area under the blood concentration versus time curve from 0 to the last time point that compound was quantifiable in blood. [h]Area under the blood concentration versus time curve from 0 to infinity. [i]Volume of distribution. [j]Total body clearance. [k]In-life plasma and CNS PK study for RPI-GLYT2-82 after 50 mg/kg, i.p. administration. Dosing groups consisted of six drug naïve adult male C57BL/6 mice. [l]Intraperitoneal (i.p.) administration. [m]Test article was administered at the 50 mg/kg dose; test article vehicle = 5% Solutol® HS-15/95% PBS). [n] Total concentration of compound in brain at 30 min and 120 min. [o] Total concentration of drug in compound at 30 min and 120 min. [q]Calculated unbound concentration of compound in brain at 30 min and 120 min. [r] Calculated unbound concentration of compound in plasma at 30 min and 120 min.

**Supplementary Table 8| Molecular dynamics system compositions.**

| | Simulation system | System composition by molecule type | | | | | | | | |
|---|---|---|---|---|---|---|---|---|---|---|
| | | **GlyT2** | **POPC** | **CHOL** | **CHS** | **Inhibitor** | **Glycine** | **Water** | **$Na^+$** | **$Cl^-$** |
| **1** | GlyT2/ORG25543 | 1 | 766 | 200 | 2 | 1 | 0 | 74086 | 335 | 341 |
| **2** | GlyT2/RPI-GLYT2-82 | 1 | 775 | 199 | 1 | 1 | 0 | 73855 | 335 | 341 |
| **3** | GlyT2/ORG25543/Gly | 1 | 766 | 200 | 2 | 1 | 1 | 74086 | 335 | 341 |
| **4** | GlyT2/ RPI-GLYT2-82/Gly | 1 | 775 | 199 | 1 | 1 | 1 | 73855 | 335 | 341 |

Simulation systems 1 & 2 were simulated for multiple replicates of 500 ns MD (system 1 n = 6, system 2 n = 3).
Simulation systems 3 & 4 were simulated for five replicates of 50 ns MD.

#### Supplementary data 1. *In Vivo* Mouse PK and *In Vitro* ADME Study Information and Data

##### Study 1 – RPI-GLYT2-82 IP PK Study and Bioanalysis in Mouse (PK Parameters)

A plasma pharmacokinetics (PK) study was performed in male ICR mice following intraperitoneal (i.p.) administration of RPI-GLYT2-82 at 30 mg/kg. The serial plasma samples were collected at 1, 2, 4, 6 and 24 hours(h) after i.p. administration from three animals at each time point.

**Testing Facility and Test Site**: Eurofins Discovery Partner, 25, Wugong 6th Road, Wugu District, New Taipei City, 24891, Taiwan

**Equipment:** Agilent Poroshell 120 EC-C18 column (2.7 µm, 3.0 x 50 mm; Agilent Technologies, Inc., USA), Animal cage (Allentown, USA), BD® lithium heparin tube (BD®, USA), Centrifuge 5810R (Eppendorf, Germany), Centrifuge tube (50 mL; Labcon, USA), Disposal syringe (1 mL, Terumo Corporation, Japan), Electronic scale (0-1000 g; Tanita Corporation, Japan), Gilson pipettes (# P200 Neo-P10N Micro Pipette; Gilson, France), Goldenrod animal lancet (4 mm, Goldenrod Corporation, USA), LC-MS/MS Triple Quad™ 5500+ (SCIEX, USA), Microcentrifuge tubes 1.5 mL click-cap (Treff AG, Switzerland), Pipette (# P200 Gilson, France), Pipette tips (Costar, USA), Polypropylene 96-well round U-bottom deep well plates and silicone microplate lids (StorPlate-96 U and StorMat-96; PerkinElmer Inc., USA), RAININ pipettes (E4 Multi E12-50XLS+, E12-300XLS, Refurbished Rainin E4™ XLS™ electronic 12 channel pipette, 30-300 µL, E4 Pipette Multi E12-1200XLS+, and EA6-300XLS; RAININ, USA), and Stop watch (Casio, China).

**TEST ARTICLE AND VEHICLE INFORMATION:**

**Dosing vehicle:** 5% Solutol® HS-15/ 95% PBS

**Dosing Solution Analysis:**
The dosing solutions were analyzed by LC-MS/MS. The measured dosing solution concentrations are shown in Table 1. The dosing solutions were diluted into mouse blood and analyzed in triplicate. All concentrations are expressed as mg/mL. The nominal dosing level was used in all calculations.

**Species and strain:** mouse; male ICR

**Number:** 3 animals

Male ICR mice weighing 30 ± 5 g were provided by BioLasco Taiwan Co., Ltd. (under Charles River Laboratories Licensee). Animals were acclimated for 3 days prior to use and were confirmed to be in good health. The animals were housed in animal cages with a space allocation of 30 x 19 x 13 cm. All animals were maintained in a controlled temperature (20 - 24° C) and humidity (30% - 70%) environment with 12 h light/dark cycles. Free access to standard lab diet [MFG (Oriental Yeast Co., Ltd., Japan)] and autoclaved tap water were granted. All aspects of this work including housing, experimentation, and animal disposal were performed in general accordance with the "Guide for the Care and Use of Laboratory Animals: Eighth Edition" (National Academies Press, Washington, D.C., 2011) in the AAALAC-accredited laboratory animal facility. In addition, the animal care and use protocol was reviewed and approved by the IACUC at Pharmacology Discovery Services (PDS) Taiwan, Ltd.

**Animal Dosing and Sample Collection Design:**

| Animal Use | |
|---|---|
| Animal Species/Strain | Mouse / Male ICR |
| Animal Weight Range (g) | 30 ± 5 g |
| Fasting Regimen | None |
| Pre-dose Observation | Body weight (g) |
| **Test Article Dosing and PK Sample Collection** | |
| Test Article ID | RPI-GLYT2-82 |
| Target for Bioanalysis | RPI-GLYT2-82 |
| Route of Administration | IP |
| Dose Levels (mg/kg) | 30 |
| Dose Volume (mL/kg) | 5 |
| Proposed Formulation/Vehicle | 5% DMSO/ 5% Solutol® HS15/ 90% of PBS |
| Number of Animals per Dosing Group | 3 |
| Total Number of Animals | 3 |
| Total Number of Plasma Samples | 15 |
| PK Sample (Plasma) Collection Time | 1, 2, 4, 6 and 24 h |
| Target Blood Sample Volume (mL) | About 0.05 mL via facial vein (1st to 4th time points) or 0.3 mL cardiac puncture (terminal time point) sampling |
| Preferred Anticoagulation | Lithium heparin |
| Preferred Sample Storage | ≤-70° C |

**Plasma Sample Collection from Mice (Serial Sampling)**

Blood aliquots were collected via facial vein (~0.05 mL) for the first four time points or cardiac puncture (~0.3 mL) for the last time point from mice in tubes coated with lithium heparin, mixed gently, and centrifuged at 2,500 ×g for 15 minutes at 4°C, within 1 h of collection. The plasma samples were then harvested and kept frozen at ≤-70°C until further processing. The plasma samples were processed using acetonitrile (ACN) precipitation and analyzed by LC-MS/MS.

**Plasma Sample Analysis**

The exposure levels (ng/mL) of RPI-GLYT2-82 in plasma samples were then determined by LC-MS/MS. Plots of plasma concentrations (mean ± SD) vs. time for RPI-GLYT2-82 were constructed. The fundamental PK parameters after IP administration were obtained from the NCA of the plasma data using WinNonlin (best-fit mode).

**Chromatographic Conditions**

| | | |
|---|---|---|
| Instrument Control No. | : | LCM-MS-002 |
| LC-MS/MS | : | Triple Quad™ 5500+ |
| Data Processor | : | Analyst 1.7 |
| Ionization Mode | : | Electrospray, Positive ions |
| Scan Mode | : | Multiple reaction monitoring (MRM) |
| Instrument Response | : | Peak area ratio |
| Regression Model | : | Weighted linear regression by $1/X^2$ |
| MRM of Analyte | : | 408.3/363.4 (RPI-GLYT2-82) |
| MRM of Internal Standard (IS) | : | 358.3/142.3 (Oxybutynin) |
| Column | : | Agilent Poroshell 120 EC-C18 column 2.7 µm (3.0 x 50 mm) |
| Column Temperature (°C) | : | 40 |
| Run Time (min) | : | 3.5 |
| Injection Volume (µL) | : | 2.0 |
| Mobile Phase | : | Mobile Phase A: ACN/FA= 100/0.2 (v/v) |
| | | Mobile Phase B: Water ($H_2O$)/FA= 100/0.2 (v/v) |
| HPLC Condition | : | |

| Time (min) | Flow Rate (µL/min) | A (%) | B (%) |
|---|---|---|---|
| 0.00 | 500 | 10.0 | 90.0 |
| 0.50 | 500 | 10.0 | 90.0 |

**Bioanalysis for Study 1: PK parameters**

**Exhibit 1.** The fundamental PK parameters ($t_{1/2}$, $T_{max}$, $C_{max}$, $AUC_{last}$, $AUC_{inf}$, AUC/D, AUC Extr, MRT, Vz_F and CL_F) of RPI-GLYT2-82 after RPI-GLYT2-82 (30 mg/kg, i.p.) administration were obtained from the NCA of the plasma data using WinNonlin (best-fit mode). The values are mean ± SD are reported.

| 30 mg/kg, IP | $t_{1/2}$ | $T_{max}$ | $C_{max}$ | $AUC_{last}$ | $AUC_{Inf}$ | AUC/D | AUC Extr | MRT | Vz_F | CL_F |
|---|---|---|---|---|---|---|---|---|---|---|
| | (h) | (h) | (ng/mL) | (h*ng/mL) | (h*ng/mL) | (h*kg*ng/mL/mg) | (%) | (h) | (L/kg) | (mL/min/kg) |
| Mouse#1 | 1.02 | 1.00 | 123 | 214 | 218 | 7.26 | 2.03 | 1.79 | 203.33 | 2294.30 |
| Mouse#2 | 1.13 | 1.00 | 184 | 287 | 293 | 9.77 | 2.22 | 1.74 | 166.23 | 1706.52 |
| Mouse#3 | 1.10 | 1.00 | 149 | 224 | 228 | 7.61 | 2.09 | 1.66 | 209.24 | 2190.33 |
| Mean | 1.08 | 1.00 | 152 | 241 | 246 | 8.21 | 2.11 | 1.73 | 192.93 | 2063.72 |
| SD | 0.05 | 0.00 | 31 | 40 | 41 | 1.36 | 0.09 | 0.07 | 23.32 | 313.68 |

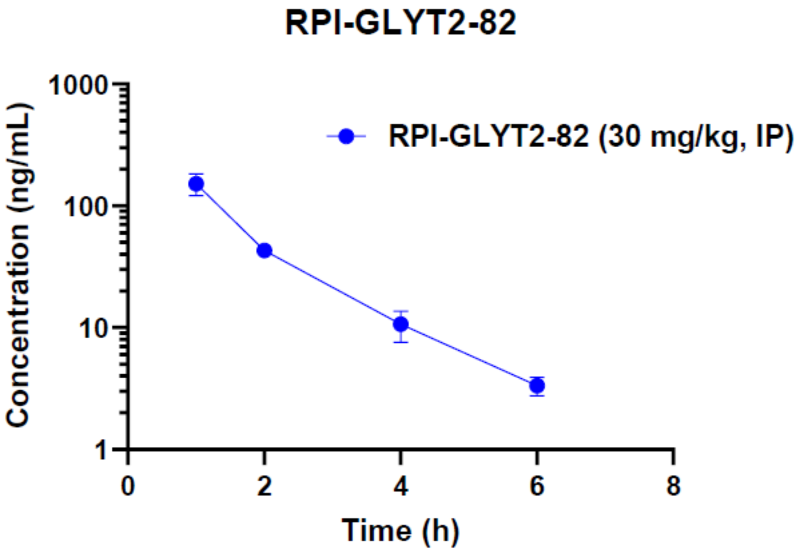

**Exhibit 2**. The exposure levels (ng/mL) of RPI-GLYT2-82 in mouse plasma samples after RPI-GLYT2- 82 (30 mg/kg, IP) administration were analyzed through the LC-MS/MS. The mean ± SD values vs. time were plotted in line chart (semi-log).

| | Sample name | Nominal conc. (ng/mL) | Area ratio | Calculated Conc. (ng/mL) | Accuracy (%) |
|---|---|---|---|---|---|
| Calibration curve | LLOQ | 1 | 0.00395 | 0.957 | 95.69 |
| | STD1 | 5 | 0.01788 | 5.981 | 119.62 |
| | STD2 | 10 | 0.02982 | 10.286 | 102.86 |
| | STD3 | 50 | 0.17639 | 63.145 | 126.29* |
| | STD4 | 100 | 0.30302 | 108.811 | 108.81 |
| | STD5 | 500 | 1.47072 | 529.930 | 105.99 |
| | STD6 | 1000 | 2.90598 | 1047.538 | 104.75 |
| | STD7 | 3000 | 7.13108 | 2571.268 | 85.71 |
| | ULOQ | 5000 | 10.61637 | 3828.194 | 76.56 |
| QCs | QL1 | 20 | 0.06085 | 21.476 | 107.38 |
| | QM1 | 2500 | 6.14130 | 2214.314 | 88.57 |
| | QH1 | 4000 | 8.71220 | 3141.477 | 78.54 |
| | QL2 | 20 | 0.05966 | 21.049 | 105.24 |
| | QM2 | 2500 | 6.04434 | 2179.346 | 87.17 |
| | QH2 | 4000 | 8.60351 | 3102.281 | 77.56 |

*: Out of concentration criteria (Accuracy <75.0% or >125.0%)

**Exhibit 3**. Analytical raw data. The exposure levels of RPI-GLYT2-82 in mouse plasma samples – RPI-GLYT2-82 (30 mg/kg, i.p.) group.

**Study 2 – RPI-GLYT2-82 In-life Plasma and CNS Drug Exposure Study in Mouse**
A plasma and brain drug exposure study was performed in male C57BL/6 mice following i.p. administration of RPI-GLYT2-82-HCl at 50 mg/kg. The brain and plasma samples were collected at 0.5 and 2 h after i.p. administration from three alternative animals at each time point.

**Testing Facility and Test Site**: Eurofins Discovery Partner, 25, Wugong 6th Road, Wugu District, New Taipei City, 24891, Taiwan

**Equipment:** 0-1000 g Electronic scale (Tanita Corporation, Japan), Animal cage (Allentown, USA), Agilent Poroshell 120 EC-C18 column (2.7 µm, 3.0 x 50 mm; Agilent Technologies, Inc., USA), BD® lithium heparin tube (BD®, USA), Centrifuge 5810R (Eppendorf, Germany), Disposal syringe (1 mL, Terumo Corporation, Japan), Gilson pipettes (# P200 Neo-P10N Micro Pipette; Gilson, France), Goldenrod Animal Lancet (4 mm, Goldenrod Corporation, USA), LC-MS/MS Triple Quad™ 5500+ (SCIEX, USA), Microcentrifuge tubes 1.5 mL click-cap (Treff AG, Switzerland), NEPHELOstar® microplate reader (BMG LabTech, Germany), Pipetman (# P200 Gilson, France), Pipette tips (Costar, USA), Polypropylene 96-well round U-bottom deep well plates and silicone microplate lids (StorPlate-96 U and StorMat-96; PerkinElmer Inc., USA), RAININ pipettes (E4 Multi E12-50XLS+, E12-300XLS, Refurbished Rainin E4™ XLS™ electronic 12 channel pipette, 30-300 µL, E4 Pipette Multi E12-1200XLS+, and EA6-300XLS; RAININ, USA), and Stop watch (Casio, China).

**TEST ARTICLE AND VEHICLE INFORMATION:**

**Dosing vehicle:** 5% Solutol® HS-15/ 95% PBS

**Dosing Solution Analysis:**
The dosing solutions were analyzed by LC-MS/MS. The measured dosing solution concentrations are shown in Table 1. The dosing solutions were diluted into mouse blood and analyzed in triplicate. All concentrations are expressed as mg/mL. The nominal dosing level was used in all calculations.

**Species and strain:** mouse; male C57BL/6

**Number:** 6 animals (three animals per timepoint group)

Male C57BL/6 mice weighing 22 ± 2 g were provided by BioLasco Taiwan Co., Ltd. (under Charles River Laboratories Licensee). Animals were acclimated for 3 days prior to use and were confirmed to be in good health. The animals were housed in animal cages with a space allocation of 30 x 19 x 13 cm. All animals were maintained in a controlled temperature (20 - 24° C) and humidity (30% - 70%) environment with 12-h light/dark cycles. Free access to standard lab diet [MFG (Oriental Yeast Co., Ltd., Japan)] and autoclaved tap water were granted. All aspects of this work including housing, experimentation, and animal disposal were performed in general accordance with the "Guide for the Care and Use of Laboratory Animals: Eighth Edition" (National Academies Press, Washington, D.C., 2011) in the AAALAC-accredited laboratory animal facility. In addition, the animal care and use protocol was reviewed and approved by the IACUC at Pharmacology Discovery Services (PDS) Taiwan, Ltd.

**Plasma Sample Collection from Mice (Parallel)**

Animals were terminated under inhalant euthanasia with $CO_2$ for blood collection by cardiac puncture. Blood aliquots (about 0.3 mL) were collected in tubes coated with lithium heparin, mixed gently, then kept on ice and centrifuged at 2,500×g for 15 min at 4°C, within 1 h of collection. The plasma was then harvested and kept frozen at ≤-70°C until further processing.

**Brain Sample Collection from Mice**

Immediately after the blood sampling, the whole brain was quickly removed, rinsed with cold saline (0.9 % NaCl), blotted with dry gauze, weighed, and stored at ≤-70°C until further processing within 1 h of collection. Each brain sample was homogenized in 0.2 mL of cold PBS, pH 7.4 on ice. The brain homogenate from each brain was then stored at ≤-70°C until further processing.

**Chromatographic Conditions**

| | | |
|---|---|---|
| Instrument Control No. | : | LCM-MS-002 |
| LC-MS/MS | : | Triple Quad™ 5500+ |
| Data Processor | : | Analyst 1.7 |
| Ionization Mode | : | Electrospray, Positive ions |
| Scan Mode | : | Multiple reaction monitoring (MRM) |
| Instrument Response | : | Peak area ratio |
| Regression Model | : | Weighted linear regression by $1/X^2$ |
| MRM of Analyte | : | 408.3/363.4 (RPI-GLYT2-82) |
| MRM of Internal Standard (IS) | : | 358.3/142.3 (Oxybutynin) |
| Column | : | Agilent Poroshell 120 EC-C18 column 2.7 µm (3.0 x 50 mm) |
| Column Temperature (°C) | : | 40 |
| Run Time (min) | : | 3.5 |
| Injection Volume (µL) | : | 2.0 |
| Mobile Phase | : | Mobile Phase A: ACN/FA= 100/0.2 (v/v)<br>Mobile Phase B: Water ($H_2O$)/FA= 100/0.2 (v/v) |
| HPLC Condition | : | |

| Time (min) | Flow Rate (µL/min) | A (%) | B (%) |
|---|---|---|---|
| 0.00 | 500 | 10.0 | 90.0 |
| 0.50 | 500 | 10.0 | 90.0 |

#### Bioanalytical Methods

| | | |
|---|---|---|
| Sample Matrix: | : | Mouse plasma and brain samples; brain samples were diluted and homogenized with 0.2 mL PBS. |
| Standard Range | : | 2 - 5000 ng/mL |
| Sample Preparation | : | Protein precipitation by ACN/FA=95/5 (v/v)<br>1) Transfer 20 µL sample in a well of 96-wells plate,<br>2) Add 300 µL of 0.01 ng/µL IS in ACN/FA=95/5 to each well, except 00 and carryover,<br>3) Add 300 µL of ACN/FA=95/5 to 00 and carryover,<br>4) Vortex 1 min and centrifuge 4000 rpm for 5 min,<br>5) Take supernatant 10 µL and add 500 µL $H_2O$/FA= 100/0.2 (v/v) for LC-MS/MS analysis. |

#### Acceptance Criteria

| Sample type | Criteria for Sample Analysis |
|---|---|
| Calibration Standards (STDs) | The calculated concentrations of the calibration STDs, including the lower limit of quantification (LLOQ) and upper limit of quantification (ULOQ), should not deviate more than 25% from the nominal value (75.0% < Accuracy < 125.0%). At least 75% of the non-zero calibration standards (e.g. 6 in 8 calibration standards) should meet the above criteria. |
| 00 (Double blank) and 0 (Blank) | 1. Analyte peak area (00 or 0) ≤ Analyte peak area (LLOQ in calibration curve)<br>2. IS peak area (00) ≤IS peak area (LLOQ in calibration curve) |
| Quality Control (QC) | The calculated of the QC samples should be within 25% of the nominal values (75.0% < Accuracy < 125.0%). At least 2/3 of the QC samples should be within the above limits. |
| Unknown Sample | 1. The analytical concentrations in the unknown samples were below the 75% LLOQ, they were as BLOQ.<br>2. The analytical concentrations in the unknown samples were above the 150% ULOQ, they were coded "AU" (above the curve limit). The original samples were then diluted with the appropriate matrix and analyzed again in a separate run. |

**Bioanalysis for Study 2**

**Exhibit 4.** Body and brain weight for each animal was recorded before dosing and at terminal. The values of mean ± SEM are reported.

| Group | Treatment | Route | Dose | Collection Time (h) | No. | Body Weight (g) | Brain Weight (g) |
|---|---|---|---|---|---|---|---|
| 1 | RPI-GLYT2-82-HCl (PT# 1280359) (RPIN-5) | IP | 5 mL/kg, 50 mg/kg, QD x 1 | 0.5 | 1 | 23 | 0.450 |
| | | | | | 2 | 24 | 0.460 |
| | | | | | 3 | 23 | 0.453 |
| | | | | | Mean | 23.3 | 0.454 |
| | | | | | SEM | 0.3 | 0.003 |
| | | | | 2 | 4 | 24 | 0.427 |
| | | | | | 5 | 24 | 0.469 |
| | | | | | 6 | 23 | 0.462 |
| | | | | | Mean | 23.7 | 0.453 |
| | | | | | SEM | 0.3 | 0.013 |

Body and brain weight for each animal was recorded before dosing and at terminal. The values of mean ± SEM are reported.

**In-life Plasma/ BBB PK Study and BioA in Mouse (Phase II- mouse PK)**

**Exhibit 5.** The exposure levels (in ng/g or ng/mL) of RPI-GLYT2-82-HCl in mouse brain and plasma samples after RPI-GLYT2-82-HCl (50 mg/kg, IP) administration were analyzed through the LC-MS/MS; the brain/plasma ratios were also calculated. The values mean ± SD are reported.

| Group | Treatment | Route | Dose | Collection Time (h) | No. | Brain concentration (ng/g) | Plasma concentration (ng/mL) | Brain/Plasma Ratio |
|---|---|---|---|---|---|---|---|---|
| 1 | RPI-GLYT2-82-HCl (PT# 1280359) (RPIN-5) | IP | 50 mg/kg, QD x 1 | 0.5 | 1 | 740 | 2478 | 0.30 |
| | | | | | 2 | 502 | 2287 | 0.22 |
| | | | | | 3 | 408 | 2124 | 0.19 |
| | | | | | Mean | 550.0 | 2296.3 | 0.24 |
| | | | | | SD | 171.1 | 177.2 | 0.06 |
| | | | | 2 | 4 | 29 | 154 | 0.19 |
| | | | | | 5 | 56 | 147 | 0.38 |
| | | | | | 6 | 32 | 97 | 0.33 |
| | | | | | Mean | 39.0 | 132.7 | 0.30 |
| | | | | | SD | 14.8 | 31.1 | 0.10 |

#### Definitions

| Item | Definitions |
|---|---|
| 00 (Double blank) | Matrix sample that no analyte and IS are added. |
| 0 (Blank) | Matrix sample that no analyte but IS is added. |
| Calibration standards (STDs) | Calibrators, or calibration standards, refer to a biological matrix to which a known amount of analyte has been added. Calibration standards are used to construct calibration curves from which the concentrations of analytes in QC samples and in-study samples are determined. |
| LLOQ | The LLOQ is the lowest amount of an analyte that can be quantitatively determined with acceptable precision and accuracy. |
| ULOQ | The ULOQ is the highest amount of an analyte in a sample that can be quantitatively determined with precision and accuracy. |
| QC | A QC is a biological matrix with a known quantity of analyte that is used to monitor the performance of a bioanalytical method and to assess the integrity and validity of the results of study samples analyzed in an individual run, including QH (high concentration quality control sample), QM (Middle/Medium concentration quality control sample), and QL (Low concentration quality control sample). |

##### Study 3 – *In Vitro* ADME Assay Information: % Plasma Protein Binding

Plasma protein binding (PPB) for RPI-GLYT2-82 in PBS (pH 7.4) was conducted by Eurofins using equilibrium dialysis of plasma with HPLC-UV/Vis detection.

**Exhibit 6.** RPI-GLYT2-82 Plasma Protein Binding Results

| Compound I.D. | Client Compound I.D. | Test Concentration | % Protein Bound | | | % Recovery | | |
|---|---|---|---|---|---|---|---|---|
| | | | 1st | 2nd | Mean | 1st | 2nd | Mean |
| **Protein binding (plasma, human)** | | | | | | | | |
| 100069868-1 | RPI-GLYT2-82 | 1.0E-05 M | 47.2 | 45.1 | 46 | 110 | 111 | 111 |
| **Protein binding (plasma, rat, Sprague-Dawley)** | | | | | | | | |
| 100069868-1 | RPI-GLYT2-82 | 1.0E-05 M | 61.1 | 54.5 | 58 | 102 | 106 | 104 |
| **Protein binding (plasma, mouse, CD-1)** | | | | | | | | |
| 100069868-1 | RPI-GLYT2-82 | 1.0E-05 M | 48.4 | 56.8 | 53 | 105 | 101 | 103 |

**Exhibit 7.** Reference Compounds protein binding results

| Compound I.D. | Test Concentration | % Protein Bound | | | % Recovery | | |
|---|---|---|---|---|---|---|---|
| | | 1st | 2nd | Mean | 1st | 2nd | Mean |
| **Protein binding (plasma, human)** | | | | | | | |
| Acebutolol | 1.0E-05 M | 26.0 | 18.4 | 22 | 105 | 111 | 108 |
| Quinidine | 1.0E-05 M | 56.1 | 57.9 | 57 | 112 | 128 | 120 |
| Sertraline | 1.0E-05 M | 98.4 | 98.2 | 98 | 102 | 104 | 103 |
| Warfarin | 1.0E-05 M | 98.2 | 97.4 | 98 | 109 | 110 | 110 |
| **Protein binding (plasma, rat, Sprague-Dawley)** | | | | | | | |
| Acebutolol | 1.0E-05 M | 19.8 | 12.8 | 16 | 92 | 100 | 96 |
| Quinidine | 1.0E-05 M | 61.7 | 57.5 | 60 | 124 | 119 | 121 |
| Sertraline | 1.0E-05 M | 97.2 | 97.5 | 97 | 99 | 94 | 97 |
| Warfarin | 1.0E-05 M | 99.2 | 99.3 | 99 | 107 | 115 | 111 |
| **Protein binding (plasma, mouse, CD-1)** | | | | | | | |
| Acebutolol | 1.0E-05 M | 14.1 | 17.0 | 16 | 68 | 74 | 71 |
| Quinidine | 1.0E-05 M | 53.0 | 63.4 | 58 | 104 | 109 | 106 |
| Sertraline | 1.0E-05 M | 94.5 | 94.4 | 94 | 109 | 108 | 108 |
| Warfarin | 1.0E-05 M | 90.4 | 88.7 | 90 | 104 | 103 | 103 |

##### Mean Plasma Protein Binding of Control Propranolol in Mouse (CD-1) Plasma

The peak areas of the test compound in the buffer and test samples were used to calculate percent binding and recovery according to the following formulas:

$$\text{Protein binding}(\%) = \frac{Area_p - Area_b}{Area_p} * 100$$

$$\text{Recovery}(\%) = \frac{Area_p + Area_b}{Area_c} * 100$$

Where:
$Area_p$ = Peak area of analyte in protein matrix
$Area_b$ = Peak area of analyte in buffer
$Area_c$ = Peak area of analyte in control sample

#### Study 4 – *In Vitro* ADME Assay Information: Mouse %Brain Tissue Homogenate Binding

Mouse %brain tissue homogenate binding for RPI-GLYT2-82 in PBS (pH 7.4) was conducted by Eurofins using equilibrium dialysis with HPLC-MS-MS detection.

**Exhibit 8.** RPI-GLYT2-82 mouse brain protein binding.

| Compound I.D. | Client Compound I.D. | Test Concentration | % Protein Bound 1st | % Protein Bound 2nd | % Protein Bound Mean | % Recovery 1st | % Recovery 2nd | % Recovery Mean |
|---|---|---|---|---|---|---|---|---|
| **Tissue binding (brain, mouse)** | | | | | | | | |
| 100069867-1 | RPI-GLYT2-82 | 1.0E-05 M | 56.2 | 55.3 | 56 | 119 | 105 | 112 |

| Compound I.D. | Test Concentration | % Protein Bound 1st | % Protein Bound 2nd | % Protein Bound Mean | % Recovery 1st | % Recovery 2nd | % Recovery Mean |
|---|---|---|---|---|---|---|---|
| **Tissue binding (brain, mouse)** | | | | | | | |
| Acebutolol | 1.0E-05 M | 15.1 | 15.2 | 15 | 110 | 124 | 117 |
| Quinidine | 1.0E-05 M | 80.3 | 79.5 | 80 | 112 | 113 | 113 |
| Sertraline | 1.0E-05 M | 99.7 | 99.6 | 100 | 114 | 112 | 113 |
| Warfarin | 1.0E-05 M | 45.2 | 39.8 | 43 | 111 | 115 | 113 |

**Study 5 – Off-Target Selectivity Screening of RPI-GLYT2-82**

A standard selectivity screening assay was performed with RPI-GLYT2-82 to determine off-target binding.

**Testing Facility and Test Site**: Eurofins Cerep, 2, rue du Professeur GARGOUÏL, B.P. 30001, 86 600 Celle l'Evescault, France

**Exhibit 9**. *In Vitro* binding assays and testing conditions for the RPI-GLYT2-82 off-target selectivity screen.

| Assay | Source | Ligand | Conc. | Kd | Non Specific | Incubation | Detection Method | Bibl. |
|---|---|---|---|---|---|---|---|---|
| **Receptors** | | | | | | | | |
| **$A_{2A}$ *(h)* (agonist radioligand)** | human recombinant (HEK-293 cells) | [$^3$H]CGS 21680 | 6 nM | 27 nM | NECA (10 µM) | 120 min RT | Scintillation counting | 141 |
| **alpha$_{1A}$ *(h)* (antagonist radioligand)** | human recombinant (CHO cells) | [$^3$H]prazosin | 0.1 nM | 0.1 nM | epinephrine (0.1 mM) | 60 min RT | Scintillation counting | 897 |
| **alpha$_{2A}$ *(h)* (antagonist radioligand)** | human recombinant (CHO cells) | [$^3$H]RX 821002 | 1 nM | 0.8 nM | (-)epinephrine (100 µM) | 60 min RT | Scintillation counting | 542 |
| **beta$_1$ *(h)* (agonist radioligand)** | human recombinant (HEK-293 cells) | [$^3$H](-)CGP 12177 | 0.3 nM | 0.39 nM | alprenolol (50 µM) | 60 min RT | Scintillation counting | 548 |
| **beta$_2$ *(h)* (antagonist radioligand)** | human recombinant (CHO cells) | [$^3$H](-)CGP 12177 | 0.3 nM | 0.15 nM | alprenolol (50 µM) | 120 min RT | Scintillation counting | 794 |
| **$CB_2$ *(h)* (agonist radioligand)** | human recombinant (CHO cells) | [$^3$H]WIN 55212-2 | 0.8 nM | 1.5 nM | WIN 55212-2 (5 µM) | 120 min 37°C | Scintillation counting | 165 |
| **$CB_1$ *(h)* (agonist radioligand)** | human recombinant (Chem-RBL cells) | [$^3$H]CP 55940 | 2 nM | 0.9 nM | AM281 (10 µM) | 30 min 22°C | Scintillation counting | 657 |
| **$CCK_1$ ($CCK_A$) *(h)* (agonist radioligand)** | human recombinant (CHO cells) | [$^{125}$I]CCK-8s | 0.08 nM | 0.24 nM | CCK-8s (1 µM) | 60 min RT | Scintillation counting | 562 |
| **$D_1$ *(h)* (antagonist radioligand)** | human recombinant (CHO cells) | [$^3$H]SCH 23390 | 0.3 nM | 0.2 nM | SCH 23390 (1 µM) | 60 min RT | Scintillation counting | 281 |
| **$D_{2S}$ *(h)* (agonist radioligand)** | human recombinant (HEK-293 cells) | [$^3$H]7-OH-DPAT | 1 nM | 0.68 nM | butaclamol (10 µM) | 60 min RT | Scintillation counting | 87 |
| **$ET_A$ *(h)* (agonist radioligand)** | human recombinant (CHO cells) | [$^{125}$I]endothelin-1 | 0.03 nM | 0.03 nM | endothelin-1 (100 nM) | 120 min 37°C | Scintillation counting | 30 |
| **$H_1$ *(h)* (antagonist radioligand)** | human recombinant (HEK-293 cells) | [$^3$H]pyrilamine | 1 nM | 1.7 nM | pyrilamine (1 µM) | 60 min RT | Scintillation counting | 492 |
| **$H_2$ *(h)* (antagonist radioligand)** | human recombinant (CHO cells) | [$^{125}$I]APT | 0.075 nM | 2.9 nM | tiotidine (100 µM) | 120 min RT | Scintillation counting | 540 |
| **$M_1$ *(h)* (antagonist radioligand)** | human recombinant (CHO cells) | [$^3$H]pirenzepine | 2 nM | 13 nM | atropine (1 µM) | 60 min RT | Scintillation counting | 59 |
| **$M_2$ *(h)* (antagonist radioligand)** | human recombinant (CHO cells) | [$^3$H]AF-DX 384 | 2 nM | 4.6 nM | atropine (1 µM) | 60 min RT | Scintillation counting | 59 |

| Assay | Source | Ligand | Conc. | Kd | Non Specific | Incubation | Detection Method | Bibl. |
|---|---|---|---|---|---|---|---|---|
| **$M_3$ *(h)* (antagonist radioligand)** | human recombinant (CHO cells) | [$^3$H]4-DAMP | 0.2 nM | 0.5 nM | atropine (1 µM) | 60 min RT | Scintillation counting | 546 |
| **N neuronal alpha4beta2 *(h)* (agonist radioligand)** | human recombinant (SH-SY5Y cells) | [$^3$H]cytisine | 0.6 nM | 0.3 nM | nicotine (10 µM) | 120 min 4°C | Scintillation counting | 1084 |
| **delta (DOP) *(h)* (agonist radioligand)** | human recombinant (Chem-1 (RBL) cells) | [$^3$H]DADLE | 0.5 nM | 0.6 nM | naltrexone (10 µM) | 60 min RT | Scintillation counting | 501 |
| **kappa *(h)* (KOP) (agonist radioligand)** | human recombinant (RBL cells) | [3H]U69593 | 0.5 nM | 0.6 nM | naloxone (10µM) | 60 min RT | Scintillation counting | 222 |
| **µ (MOP) *(h)* (agonist radioligand)** | human recombinant (HEK-293 cells) | [$^3$H]DAMGO | 0.5 nM | 0.35 nM | naloxone (10 µM) | 120 min RT | Scintillation counting | 260 |
| **5-$HT_{1A}$ *(h)* (agonist radioligand)** | human recombinant (HEK-293 cells) | [$^3$H]8-OH-DPAT | 0.5 nM | 0.5 nM | 8-OH-DPAT (10 µM) | 60 min RT | Scintillation counting | 164 |
| **5-$HT_{1B}$ *(h)* (antagonist radioligand)** | human recombinant (Chem-1 (RBL) cells) | [3H]GR125743 | 1 nM | 0.8 nM | Serotonine (30 µM) | 60 min 37°C | Scintillation counting | 1451 |
| **5-$HT_{2A}$ *(h)* (agonist radioligand)** | human recombinant (HEK-293 cells) | [$^{125}$I](±)DOI | 0.1 nM | 0.3 nM | (±)DOI (1 µM) | 60 min RT | Scintillation counting | 288 |
| **5-$HT_{2B}$ *(h)* (agonist radioligand)** | human recombinant (CHO cells) | [$^{125}$I](±)DOI | 0.2 nM | 0.2 nM | (±)DOI (1 µM) | 60 min RT | Scintillation counting | 571 |
| **GR *(h)* (agonist radioligand)** | human endogenous (IM-9 cells) | [$^3$H]dexamethasone | 1.5 nM | 1.5 nM | triamcinolone (10 µM) | 24 hr 4°C | Scintillation counting | 283 |
| **AR(h) (agonist radioligand)** | human endogenous (LNCaP cells) | [$^3$H]methyltrienolone | 1 nM | 0.8 nM | testostérone (1 µM) | 4 hr 22°C | Scintillation counting | 498 |
| **$V_{1a}$ *(h)* (agonist radioligand)** | human recombinant (CHO cells) | [$^3$H]AVP | 0.3 nM | 0.5 nM | AVP (1 µM) | 60 min RT | Scintillation counting | 343 |
| **Ion channels** | | | | | | | | |
| **BZD (central) (agonist radioligand)** | rat cerebral cortex | [$^3$H]flunitrazepam | 0.4 nM | 2.1 nM | diazepam (3 µM) | 60 min 4°C | Scintillation counting | 227 |
| **NMDA (antagonist radioligand)** | rat cerebral cortex | [$^3$H]CGP 39653 | 5 nM | 23 nM | L-glutamate (100 µM) | 60 min 4°C | Scintillation counting | 221 |
| **5-$HT_3$ *(h)* (antagonist radioligand)** | human recombinant (CHO cells) | [$^3$H]BRL 43694 | 0.5 nM | 1.15 nM | MDL 72222 (10 µM) | 120 min RT | Scintillation counting | 109 |
| **$Ca^{2+}$ channel (L, dihydropyridine site) (antagonist radioligand)** | rat cerebral cortex | [$^3$H]nitrendipine | 0.25 nM | 0.27 nM | nitrendipine (1 µM) | 90 min RT | Scintillation counting | 996 |

| Assay | Source | Ligand | Conc. | Kd | Non Specific | Incubation | Detection Method | Bibl. |
|---|---|---|---|---|---|---|---|---|
| **Potassium Channel hERG (human)- [3H] Dofetilide** | human recombinant (HEK-293 cells) | [3H]Dofetilide | 3 nM | 6.6 nM | Terfenadine (25 µM) | 60 min RT | Scintillation counting | 1398 |
| **$K_V$ channel (antagonist radioligand)** | rat cerebral cortex | [$^{125}$I]α-dendrotoxin | 0.01 nM | 0.04 nM | α-dendrotoxin (50 nM) | 60 min RT | Scintillation counting | 225 |
| **$Na^+$ channel (site 2) (antagonist radioligand)** | rat cerebral cortex | [$^3$H]batrachotoxin | 10 nM | 31 nM | veratridine (300 µM) | 60 min 37°C | Scintillation counting | 28 |
| **Transporters** | | | | | | | | |
| **norepinephrine transporter *(h)* (antagonist radioligand)** | human recombinant (CHO cells) | [$^3$H]nisoxetine | 1 nM | 2.9 nM | desipramine (1 µM) | 120 min 4°C | Scintillation counting | 180 |
| **dopamine transporter *(h)* (antagonist radioligand)** | human recombinant (CHO cells) | [$^3$H]BTCP | 4 nM | 4.5 nM | BTCP (10 µM) | 120 min 4°C | Scintillation counting | 190 |
| **5-HT transporter *(h)* (antagonist radioligand)** | human recombinant (CHO cells) | [$^3$H]imipramine | 2 nM | 1.7 nM | imipramine (10 µM) | 60 min RT | Scintillation counting | 566 |
| **Other enzymes** | | | | | | | | |
| **MAO-A (antagonist radioligand)** | rat cerebral cortex | [$^3$H]Ro 41-1049 | 10 nM | 14 nM | clorgyline (1 µM) | 60 min 37°C | Scintillation counting | 36 |

| Assay | Source | Substrate/ Stimulus/Tracer | Incubation | Measured Component | Detection Method | Bibl. |
|---|---|---|---|---|---|---|
| **Kinases** | | | | | | |
| **Lck kinase *(h)*** | human recombinant (insect cells) | ATP + Ulight-Poly GAT[EAY(1:1:1)]n (25 nM) | 10 min RT | phospho-Ulight-Poly GAT[EAY(1:1:1)]n | LANCE | 556 |
| **Other enzymes** | | | | | | |
| **COX1(h)** | human recombinant | Arachidonic acid (3µM) + ADHP ( 25 µM) | 3 min RT | Resorufin (oxydized ADHP) | Fluorimetry | 1480 |
| **COX2(h)** | human recombinant (Sf9 cells) | arachidonic acid (1.2 µM)+ ADHP (25 µM) | 5 min RT | Resorufin (oxydized ADHP) | Fluorimetry | 1480 |
| **PDE3A *(h)*** | human recombinant (Sf21 cells) | [3H]cAMP + cAMP (0.5µM) | 15 min RT | [3H]5'AMP | Scintillation counting | 1399 |
| **PDE4D2 *(h)*** | human recombinant (Sf9 cells) | [3H]cAMP + cAMP (0.5µM) | 20 min RT | [3H]5'AMP | Scintillation counting | 1399 |
| **acetylcholinesterase *(h)*** | human recombinant (HEK-293 cells) | Acetylthiocholine (400 µM) | 30 min RT | 5 thio 2 nitrobenzoic acid | Photometry | 63 |

**Analysis and expression of results – *In Vitro* binding assays**

The results are expressed as a percent of control specific binding

(Measured specific binding/control specific binding) × 100

and as a percent inhibition of control specific binding

100 – [(Measured specific binding/control specific binding) × 100]

obtained in the presence of RPI-GLYT2-82.

The $IC_{50}$ values (concentration causing a half-maximal inhibition of control specific binding) and Hill coefficients (nH) were determined by non-linear regression analysis of the competition curves generated with mean replicate values using Hill equation curve fitting:

$$Y = D + \left[\frac{A-D}{1+\left(\frac{C}{C_{50}}\right)^{nH}}\right]$$

where Y = specific binding, A = left asymptote of the curve, D = right asymptote of the curve, C = compound concentration, $C_{50} = IC_{50}$, and nH = slope factor. This analysis was performed using software developed at Cerep (Hill software) and validated by comparison with data generated by the commercial software SigmaPlot® 4.0 for Windows® (© 1997 by SPSS Inc.). The inhibition constants (Ki) were calculated using the Cheng Prusoff equation:

$$K_i = \frac{IC_{50}}{\left(1 + \frac{L}{K_D}\right)}$$

where L = concentration of ligand in the assay, and $K_D$ = affinity of the ligand for the receptor.

**Analysis and expression of results – *In Vitro* enzyme and uptake assays**

The results are expressed as a percent of control specific activity

(Measured specific activity/control specific activity) × 100

and as a percent inhibition of control specific activity

100 – [(Measured specific activity/control specific activity) × 100]

obtained in the presence of RPI-GLYT2-82.

The $IC_{50}$ values (concentration causing a half-maximal inhibition of control specific activity) and Hill coefficients (nH) were determined by non-linear regression analysis of the competition curves generated with mean replicate values using Hill equation curve fitting:

$$Y = D + \left[\frac{A-D}{1+\left(\frac{C}{C_{50}}\right)^{nH}}\right]$$

where Y = specific activity, A = left asymptote of the curve, D = right asymptote of the curve, C = compound concentration, $C_{50} = IC_{50}$, and nH = slope factor. This analysis was performed using software developed at Cerep (Hill software) and validated by comparison with data generated by the commercial software SigmaPlot® 4.0 for Windows® (© 1997 by SPSS Inc.).

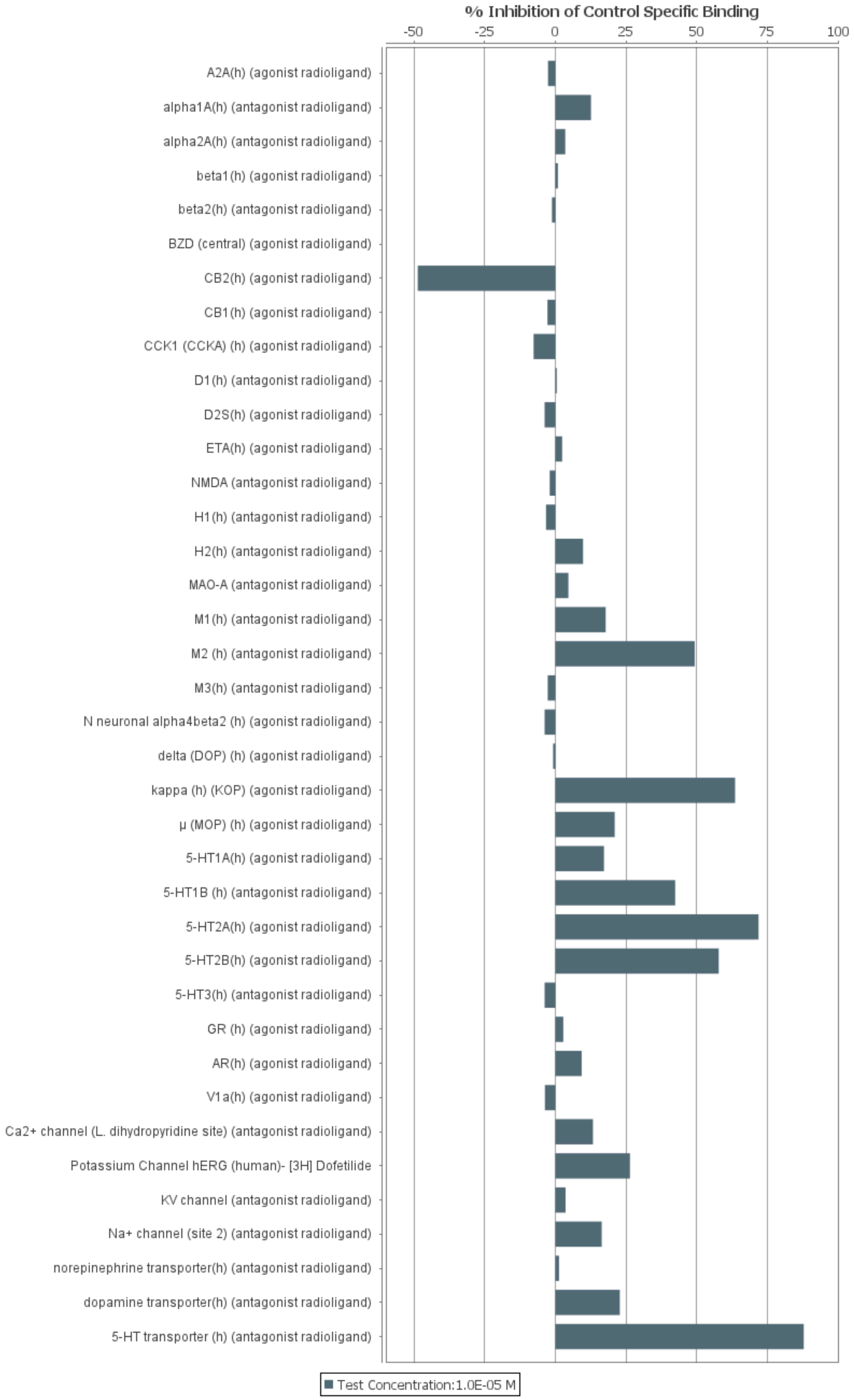

% Inhibition of Control Specific Binding
-50
-25
0
25
50
75
100
A2A(h) (agonist radioligand)
alpha1A(h) (antagonist radioligand)
alpha2A(h) (antagonist radioligand)
beta1(h) (agonist radioligand)
beta2(h) (antagonist radioligand)
BZD (central) (agonist radioligand)
CB2(h) (agonist radioligand)
CB1(h) (agonist radioligand)
CCK1 (CCKA) (h) (agonist radioligand)
D1(h) (antagonist radioligand)
D2S(h) (agonist radioligand)
ETA(h) (agonist radioligand)
NMDA (antagonist radioligand)
H1(h) (antagonist radioligand)
H2(h) (antagonist radioligand)
MAO-A (antagonist radioligand)
M1(h) (antagonist radioligand)
M2 (h) (antagonist radioligand)
M3(h) (antagonist radioligand)
N neuronal alpha4beta2 (h) (agonist radioligand)
delta (DOP) (h) (agonist radioligand)
kappa (h) (KOP) (agonist radioligand)
μ (MOP) (h) (agonist radioligand)
5-HT1A(h) (agonist radioligand)
5-HT1B (h) (antagonist radioligand)
5-HT2A(h) (agonist radioligand)
5-HT2B(h) (agonist radioligand)
5-HT3(h) (antagonist radioligand)
GR (h) (agonist radioligand)
AR(h) (agonist radioligand)
V1a(h) (agonist radioligand)
Ca2+ channel (L. dihydropyridine site) (antagonist radioligand)
Potassium Channel hERG (human)- [3H] Dofetilide
KV channel (antagonist radioligand)
Na+ channel (site 2) (antagonist radioligand)
norepinephrine transporter(h) (antagonist radioligand)
dopamine transporter(h) (antagonist radioligand)
5-HT transporter (h) (antagonist radioligand)
Test Concentration:1.0E-05 M

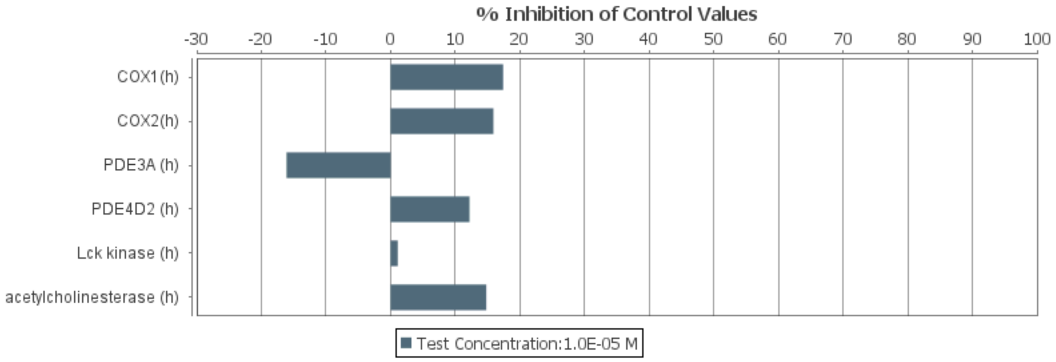

**Exhibit 10.** RPI-GLYT2-82 selectivity screening. RPI-GLYT2-82 was tested at $1 \times 10^{-5}$ M. Compound binding was calculated as a % inhibition of the binding of a ligand specific for each target (upper graph). Compound enzyme inhibition effect was calculated as a % inhibition of control enzyme activity (bottom graph).

**Exhibit 11.** Test compound (RPI-GLYT2-82) results.

| Compound I.D. | Client Compound I.D. | Test Concentration | % Inhibition of Control Specific Binding | | |
|---|---|---|---|---|---|
| | | | 1st | 2nd | Mean |
| **$A_{2A}$*(h)* (agonist radioligand)** | | | | | |
| 100069967-1 | RPI-GLYT2-82 | 1.0E-05 M | -2.5 | -2.7 | -2.6 |
| **alpha$_{1A}$*(h)* (antagonist radioligand)** | | | | | |
| 100069967-1 | RPI-GLYT2-82 | 1.0E-05 M | 11.2 | 14.0 | 12.6 |
| **alpha$_{2A}$*(h)* (antagonist radioligand)** | | | | | |
| 100069967-1 | RPI-GLYT2-82 | 1.0E-05 M | 4.7 | 2.4 | 3.5 |
| **beta$_1$*(h)* (agonist radioligand)** | | | | | |
| 100069967-1 | RPI-GLYT2-82 | 1.0E-05 M | 2.1 | -0.2 | 0.9 |
| **beta$_2$*(h)* (antagonist radioligand)** | | | | | |
| 100069967-1 | RPI-GLYT2-82 | 1.0E-05 M | 0.1 | -2.6 | -1.2 |
| **BZD (central) (agonist radioligand)** | | | | | |
| 100069967-1 | RPI-GLYT2-82 | 1.0E-05 M | 4.0 | -4.0 | 0.0 |
| **$CB_2$*(h)* (agonist radioligand)** | | | | | |
| 100069967-1 | RPI-GLYT2-82 | 1.0E-05 M | -40.2 | -57.2 | -48.7 |
| **$CB_1$*(h)* (agonist radioligand)** | | | | | |
| 100069967-1 | RPI-GLYT2-82 | 1.0E-05 M | -6.4 | 0.8 | -2.8 |
| **$CCK_1$ ($CCK_A$) *(h)* (agonist radioligand)** | | | | | |
| 100069967-1 | RPI-GLYT2-82 | 1.0E-05 M | -14.6 | -0.8 | -7.7 |
| **$D_1$*(h)* (antagonist radioligand)** | | | | | |
| 100069967-1 | RPI-GLYT2-82 | 1.0E-05 M | -5.7 | 6.7 | 0.5 |
| **$D_{2S}$*(h)* (agonist radioligand)** | | | | | |
| 100069967-1 | RPI-GLYT2-82 | 1.0E-05 M | -6.9 | -0.7 | -3.8 |
| **$ET_A$*(h)* (agonist radioligand)** | | | | | |
| 100069967-1 | RPI-GLYT2-82 | 1.0E-05 M | 5.8 | -0.9 | 2.4 |
| **NMDA (antagonist radioligand)** | | | | | |
| 100069967-1 | RPI-GLYT2-82 | 1.0E-05 M | -3.1 | -0.9 | -2.0 |
| **$H_1$*(h)* (antagonist radioligand)** | | | | | |
| 100069967-1 | RPI-GLYT2-82 | 1.0E-05 M | -2.1 | -4.6 | -3.3 |
| **$H_2$*(h)* (antagonist radioligand)** | | | | | |
| 100069967-1 | RPI-GLYT2-82 | 1.0E-05 M | 8.9 | 10.6 | 9.8 |
| **MAO-A (antagonist radioligand)** | | | | | |
| 100069967-1 | RPI-GLYT2-82 | 1.0E-05 M | 10.8 | -1.7 | 4.6 |
| **$M_1$*(h)* (antagonist radioligand)** | | | | | |
| 100069967-1 | RPI-GLYT2-82 | 1.0E-05 M | 17.4 | 18.3 | 17.8 |
| **$M_2$ *(h)* (antagonist radioligand)** | | | | | |
| 100069967-1 | RPI-GLYT2-82 | 1.0E-05 M | 49.0 | 49.6 | 49.3 |
| **$M_3$*(h)* (antagonist radioligand)** | | | | | |
| 100069967-1 | RPI-GLYT2-82 | 1.0E-05 M | -8.8 | 3.3 | -2.7 |
| **N neuronal alpha4beta2 *(h)* (agonist radioligand)** | | | | | |
| 100069967-1 | RPI-GLYT2-82 | 1.0E-05 M | -1.7 | -5.8 | -3.8 |
| **delta (DOP) *(h)* (agonist radioligand)** | | | | | |
| 100069967-1 | RPI-GLYT2-82 | 1.0E-05 M | 0.2 | -1.8 | -0.8 |
| **kappa *(h)* (KOP) (agonist radioligand)** | | | | | |
| 100069967-1 | RPI-GLYT2-82 | 1.0E-05 M | 63.0 | 64.2 | **63.6** |
| **μ (MOP) *(h)* (agonist radioligand)** | | | | | |
| 100069967-1 | RPI-GLYT2-82 | 1.0E-05 M | 16.4 | 25.5 | 21.0 |
| **5-$HT_{1A}$*(h)* (agonist radioligand)** | | | | | |
| 100069967-1 | RPI-GLYT2-82 | 1.0E-05 M | 20.2 | 14.3 | 17.2 |
| **5-$HT_{1B}$ *(h)* (antagonist radioligand)** | | | | | |
| 100069967-1 | RPI-GLYT2-82 | 1.0E-05 M | 39.4 | 45.3 | 42.4 |
| **5-$HT_{2A}$*(h)* (agonist radioligand)** | | | | | |
| 100069967-1 | RPI-GLYT2-82 | 1.0E-05 M | 73.7 | 70.1 | **71.9** |

| Compound I.D. | Client Compound I.D. | Test Concentration | % Inhibition of Control Specific Binding | | |
|---|---|---|---|---|---|
| | | | 1st | 2nd | Mean |
| **$5\text{-}HT_{2B}$*(h)* (agonist radioligand)** | | | | | |
| 100069967-1 | RPI-GLYT2-82 | 1.0E-05 M | 57.9 | 57.7 | **57.8** |
| **$5\text{-}HT_3$*(h)* (antagonist radioligand)** | | | | | |
| 100069967-1 | RPI-GLYT2-82 | 1.0E-05 M | -2.3 | -5.3 | -3.8 |
| **GR *(h)* (agonist radioligand)** | | | | | |
| 100069967-1 | RPI-GLYT2-82 | 1.0E-05 M | 0.6 | 5.0 | 2.8 |
| **AR(h) (agonist radioligand)** | | | | | |
| 100069967-1 | RPI-GLYT2-82 | 1.0E-05 M | 5.6 | 12.9 | 9.3 |
| **$V_{1a}$*(h)* (agonist radioligand)** | | | | | |
| 100069967-1 | RPI-GLYT2-82 | 1.0E-05 M | -2.6 | -4.8 | -3.7 |
| **$Ca^{2+}$ channel (L, dihydropyridine site) (antagonist radioligand)** | | | | | |
| 100069967-1 | RPI-GLYT2-82 | 1.0E-05 M | 20.2 | 6.4 | 13.3 |
| **Potassium Channel hERG (human)- [3H] Dofetilide** | | | | | |
| 100069967-1 | RPI-GLYT2-82 | 1.0E-05 M | 26.3 | 26.5 | 26.4 |
| **$K_V$ channel (antagonist radioligand)** | | | | | |
| 100069967-1 | RPI-GLYT2-82 | 1.0E-05 M | 3.1 | 4.1 | 3.6 |
| **$Na^+$ channel (site 2) (antagonist radioligand)** | | | | | |
| 100069967-1 | RPI-GLYT2-82 | 1.0E-05 M | 20.0 | 12.8 | 16.4 |
| **norepinephrine transporter*(h)* (antagonist radioligand)** | | | | | |
| 100069967-1 | RPI-GLYT2-82 | 1.0E-05 M | 10.3 | -7.8 | 1.3 |
| **dopamine transporter*(h)* (antagonist radioligand)** | | | | | |
| 100069967-1 | RPI-GLYT2-82 | 1.0E-05 M | 19.7 | 25.9 | 22.8 |
| **5-HT transporter *(h)* (antagonist radioligand)** | | | | | |
| 100069967-1 | RPI-GLYT2-82 | 1.0E-05 M | 88.6 | 87.3 | **87.9** |

| Compound I.D. | Client Compound I.D. | Test Concentration | % Inhibition of Control Values | | |
|---|---|---|---|---|---|
| | | | 1st | 2nd | Mean |
| **COX1(h)** | | | | | |
| 100069967-1 | RPI-GLYT2-82 | 1.0E-05 M | 11.7 | 23.2 | 17.4 |
| **COX2(h)** | | | | | |
| 100069967-1 | RPI-GLYT2-82 | 1.0E-05 M | 18.7 | 13.2 | 15.9 |
| **PDE3A *(h)*** | | | | | |
| 100069967-1 | RPI-GLYT2-82 | 1.0E-05 M | -17.6 | -14.7 | -16.1 |
| **PDE4D2 *(h)*** | | | | | |
| 100069967-1 | RPI-GLYT2-82 | 1.0E-05 M | 12.3 | 12.1 | 12.2 |
| **Lck kinase *(h)*** | | | | | |
| 100069967-1 | RPI-GLYT2-82 | 1.0E-05 M | 5.4 | -3.1 | 1.1 |
| **acetylcholinesterase *(h)*** | | | | | |
| 100069967-1 | RPI-GLYT2-82 | 1.0E-05 M | 15.3 | 14.4 | 14.8 |

**Exhibit 12.** Reference compound results.

| Compound I.D. | $IC_{50}$(M) | $K_i$(M) | nH |
|---|---|---|---|
| **$A_{2A}$*(h)* (agonist radioligand)** | | | |
| NECA | 6.3E-08 M | 5.2E-08 M | 0.7 |
| **$alpha_{1A}$*(h)* (antagonist radioligand)** | | | |
| WB 4101 | 3.8E-10 M | 1.9E-10 M | 0.9 |
| **$alpha_{2A}$*(h)* (antagonist radioligand)** | | | |
| yohimbine | 7.1E-09 M | 3.1E-09 M | 1.0 |
| **$beta_1$*(h)* (agonist radioligand)** | | | |
| atenolol | 2.3E-07 M | 1.3E-07 M | 1.0 |
| **$beta_2$*(h)* (antagonist radioligand)** | | | |
| ICI 118551 | 1.6E-09 M | 5.2E-10 M | 0.9 |
| **BZD (central) (agonist radioligand)** | | | |
| diazepam | 9.8E-09 M | 8.3E-09 M | 0.8 |
| **$CB_2$*(h)* (agonist radioligand)** | | | |
| WIN 55212-2 | 1.0E-09 M | 6.8E-10 M | 1.6 |
| **$CB_1$*(h)* (agonist radioligand)** | | | |
| CP 55940 | 2.7E-09 M | 8.3E-10 M | 1.4 |
| **$CCK_1$ ($CCK_A$) *(h)* (agonist radioligand)** | | | |
| CCK-8s | 7.9E-11 M | 5.9E-11 M | 1.0 |
| **$D_1$*(h)* (antagonist radioligand)** | | | |
| SCH 23390 | 4.0E-10 M | 1.6E-10 M | 1.3 |
| **$D_{2S}$*(h)* (agonist radioligand)** | | | |
| 7-OH-DPAT | 2.9E-09 M | 1.2E-09 M | >3 |
| **$ET_A$*(h)* (agonist radioligand)** | | | |
| endothelin-1 | 4.1E-11 M | 2.0E-11 M | 1.1 |
| **NMDA (antagonist radioligand)** | | | |
| CGS 19755 | 3.2E-07 M | 2.6E-07 M | 0.9 |
| **$H_1$*(h)* (antagonist radioligand)** | | | |
| pyrilamine | 2.5E-09 M | 1.5E-09 M | 1.1 |
| **$H_2$*(h)* (antagonist radioligand)** | | | |
| cimetidine | 5.9E-07 M | 5.8E-07 M | 1.4 |
| **MAO-A (antagonist radioligand)** | | | |
| clorgyline | 2.1E-09 M | 1.2E-09 M | 1.9 |
| **$M_1$*(h)* (antagonist radioligand)** | | | |
| pirenzepine | 4.0E-08 M | 3.4E-08 M | >3 |
| **$M_2$ *(h)* (antagonist radioligand)** | | | |
| methoctramine | 6.1E-08 M | 4.3E-08 M | 0.8 |
| **$M_3$*(h)* (antagonist radioligand)** | | | |
| 4-DAMP | 2.0E-09 M | 1.4E-09 M | 1.7 |
| **N neuronal alpha4beta2 *(h)* (agonist radioligand)** | | | |
| nicotine | 6.8E-09 M | 2.3E-09 M | 0.8 |
| **delta (DOP) *(h)* (agonist radioligand)** | | | |
| DPDPE | 2.6E-09 M | 1.4E-09 M | 1.1 |
| **kappa *(h)* (KOP) (agonist radioligand)** | | | |
| U50488 | 7.3E-10 M | 4.0E-10 M | 1.3 |
| **μ (MOP) *(h)* (agonist radioligand)** | | | |
| DAMGO | 9.4E-10 M | 3.9E-10 M | 0.8 |
| **5-$HT_{1A}$*(h)* (agonist radioligand)** | | | |
| 8-OH-DPAT | 6.0E-10 M | 3.0E-10 M | 0.8 |
| **5-$HT_{1B}$ *(h)* (antagonist radioligand)** | | | |
| Serotonine | 3.5E-07 M | 1.5E-07 M | 1.2 |

| Compound I.D. | $IC_{50}$(M) | $K_i$(M) | nH |
|---|---|---|---|
| **$5\text{-}HT_{2A}$*(h)* (agonist radioligand)** | | | |
| (±)DOI | 2.2E-10 M | 1.6E-10 M | 0.8 |
| **$5\text{-}HT_{2B}$*(h)* (agonist radioligand)** | | | |
| (±)DOI | 2.6E-09 M | 1.3E-09 M | 0.8 |
| **$5\text{-}HT_3$*(h)* (antagonist radioligand)** | | | |
| MDL 72222 | 8.0E-09 M | 5.6E-09 M | 1.0 |
| **GR *(h)* (agonist radioligand)** | | | |
| dexamethasone | 1.6E-09 M | 7.9E-10 M | 1.0 |
| **AR(h) (agonist radioligand)** | | | |
| testosterone | 6.8E-09 M | 3.0E-09 M | 1.3 |
| **$V_{1a}$*(h)* (agonist radioligand)** | | | |
| $[d(CH_2)_5{}^1,Tyr(Me)_2]$-AVP | 8.2E-10 M | 5.1E-10 M | 1.1 |
| **$Ca^{2+}$ channel (L, dihydropyridine site) (antagonist radioligand)** | | | |
| nitrendipine | 9.6E-10 M | 5.0E-10 M | 0.8 |
| **Potassium Channel hERG (human)- [3H] Dofetilide** | | | |
| Terfenadine | 6.6E-08 M | 4.5E-08 M | 1.3 |
| **$K_V$ channel (antagonist radioligand)** | | | |
| α-dendrotoxin | 8.3E-11 M | 6.6E-11 M | 0.7 |
| **$Na^+$ channel (site 2) (antagonist radioligand)** | | | |
| veratridine | 1.2E-05 M | 9.1E-06 M | 0.7 |
| **norepinephrine transporter*(h)* (antagonist radioligand)** | | | |
| protriptyline | 5.0E-09 M | 3.7E-09 M | 1.3 |
| **dopamine transporter*(h)* (antagonist radioligand)** | | | |
| BTCP | 1.1E-08 M | 5.7E-09 M | 1.2 |
| **5-HT transporter *(h)* (antagonist radioligand)** | | | |
| imipramine | 2.5E-09 M | 1.2E-09 M | 1.1 |

| Compound I.D. | $IC_{50}$(M) | nH |
|---|---|---|
| **COX1(h)** | | |
| Diclofenac | 2.7E-08 M | 1.8 |
| **COX2(h)** | | |
| NS398 | 5.1E-07 M | 2.1 |
| **PDE3A *(h)*** | | |
| milrinone | 2.7E-07 M | 0.8 |
| **PDE4D2 *(h)*** | | |
| Ro 20-1724 | 1.7E-07 M | 0.6 |
| **Lck kinase *(h)*** | | |
| staurosporine | 3.3E-08 M | 2.0 |
| **acetylcholinesterase *(h)*** | | |
| galanthamine | 5.5E-07 M | 0.9 |

#### Further analysis– *In Vitro* cellular receptor functional, enzyme and uptake assays

To further characterise non-specific activity of RPI-GLYT2-82, results showing an inhibition or stimulation higher than 50% were considered to represent significant effects of the test compounds. This threshold was reached with the κ-opioid receptor (KOP), 5-HT2A receptor and the serotonin transporter and were investigated further.

**Exhibit 13.** $EC_{50}$ determination: Test compound (RPI-GLYT2-82) results

| Compound I.D. | Client Compound I.D. | $EC_{50}$ | Test Concentration | % of Control Agonist Response 1st | 2nd | Mean |
|---|---|---|---|---|---|---|
| **κ (KOP) (h) (agonist effect)** | | | | | | |
| 100070115-1 | RPI-GLYT2-82 | N.C. | 3.0E-08 M | 1.0 | -1.4 | -0.2 |
| | | | 1.0E-07 M | -1.3 | -1.4 | -1.3 |
| | | | 3.0E-07 M | 1.4 | 0.7 | 1.0 |
| | | | 1.0E-06 M | -0.6 | 0.4 | -0.1 |
| | | | 2.0E-06 M | -1.1 | 0.8 | -0.2 |
| | | | 5.0E-06 M | 2.6 | -0.1 | 1.3 |
| | | | 1.0E-05 M | -1.2 | -0.6 | -0.9 |
| | | | 3.0E-05 M | -0.4 | -0.9 | -0.7 |

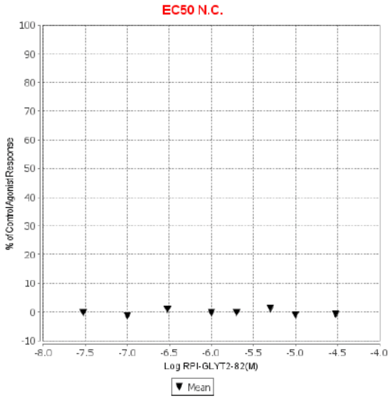

**Figure 1. RPI-GLYT2-82 on κ (KOP) (h) (agonist effect)**

*N.C.*: $EC_{50}$ value not calculable. Concentration-response curve shows less than 25% effect at the highest validated testing concentration.

Reference agonist result

| Compound I.D. | $EC_{50}$(M) | nH |
|---|---|---|
| **κ (KOP) (h) (agonist effect)** | | |
| U50,488 | 6.2E-08 M | n/a |

**Exhibit 14.** $IC_{50}$ determination: Test compound (RPI-GLYT2-82) results

| Compound I.D. | Client Compound I.D. | $IC_{50}$(M) | $K_B$(M) | nH | Test Concentration | % Inhibition of Control Agonist Response 1st | 2nd | Mean |
|---|---|---|---|---|---|---|---|---|
| **κ (KOP) (h) (antagonist effect)** | | | | | | | | |
| 100070115-1 | RPI-GLYT2-82 | N.C. | n/a | n/a | 3.0E-08 M | -2.8 | 7.2 | 2.2 |
| | | | | | 1.0E-07 M | 7.9 | 12.1 | 10.0 |
| | | | | | 3.0E-07 M | -1.9 | -6.7 | -4.3 |
| | | | | | 1.0E-06 M | -5.9 | 0.8 | -2.5 |
| | | | | | 2.0E-06 M | 0.8 | -14.6 | -6.9 |
| | | | | | 5.0E-06 M | -13.2 | -8.6 | -10.9 |
| | | | | | 1.0E-05 M | -10.0 | 6.8 | -1.6 |
| | | | | | 3.0E-05 M | 0.1 | -5.7 | -2.8 |

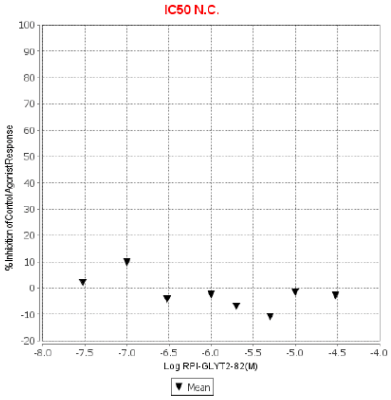

**Figure 3. RPI-GLYT2-82 on κ (KOP) (h) (antagonist effect)**

| | | | | | | | | |
|---|---|---|---|---|---|---|---|---|
| **5-HT$_{2A}$*(h)* (antagonist effect)** | | | | | | | | |
| 100070115-1 | RPI-GLYT2-82 | 1.9E-06 M | 3.8E-07 M | n/a | 3.0E-09 M | -12.0 | 15.0 | 1.5 |
| | | | | | 3.0E-08 M | -2.1 | -9.8 | -6.0 |
| | | | | | 1.0E-07 M | -7.0 | -12.0 | -9.5 |
| | | | | | 3.0E-07 M | -12.9 | 0.7 | -6.1 |
| | | | | | 1.0E-06 M | 36.8 | 21.1 | 29.0 |
| | | | | | 3.0E-06 M | 64.7 | 58.2 | 61.4 |
| | | | | | 1.0E-05 M | 86.0 | 87.0 | 86.5 |
| | | | | | 1.0E-04 M | 101.6 | 101.2 | 101.4 |

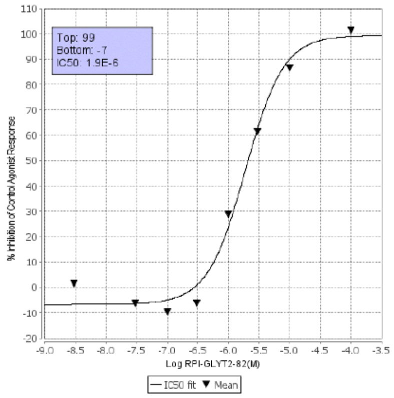

**Figure 4. RPI-GLYT2-82 on 5-HT$_{2A}$*(h)* (antagonist effect)**

*N.C.*: IC$_{50}$ value not calculable. Concentration-response curve shows less than 25% effect at the highest validated testing concentration.

| | | | | | | | | | |
|---|---|---|---|---|---|---|---|---|---|
| **Serotonin transporter uptake (h)** | | | | | | | | | |
| 100070115-1 | RPI-GLYT2-82 | 4.7E-06 M | n/a | 1.0E-09 M | -31.2 | -32.9 | -32.1 | {}, OUTLIER | {}, OUTLIER |
| | | | | 1.0E-08 M | -9.3 | -14.3 | -11.8 | | |
| | | | | 3.0E-08 M | -8.3 | -2.6 | -5.5 | | |
| | | | | 1.0E-07 M | 16.1 | 12.0 | 14.0 | | |
| | | | | 3.0E-07 M | 8.2 | 5.5 | 6.8 | | |
| | | | | 1.0E-06 M | 26.5 | 21.7 | 24.1 | | |
| | | | | 3.0E-06 M | 27.9 | 34.7 | 31.3 | | |
| | | | | 3.0E-05 M | 79.2 | 82.4 | 80.8 | | |

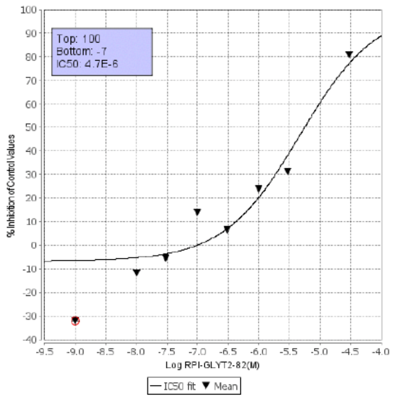

**Figure 5. RPI-GLYT2-82 on Serotonin transporter uptake (h)**

*{}*: That replicate was excluded from the calculation
*OUTLIER*: Data is outside of the expected range of values for the data set and was excluded from calculation.

### Reference compounds results

| Compound I.D. | IC$_{50}$(M) | K$_B$(M) | nH |
|---|---|---|---|
| **κ (KOP) (h) (antagonist effect)** | | | |
| ML190 | 4.6E-09 M | 8.4E-10 M | n/a |
| **5-HT$_{2A}$*(h)* (antagonist effect)** | | | |
| ketanserin | 3.9E-09 M | 7.7E-10 M | n/a |

| | | |
|---|---|---|
| **Serotonin transporter uptake (h)** | | |
| imipramine | 2.7E-08 M | n/a |

#### Supplementary methods – Synthesis of RPI-GLYT2-82

All reactions were performed under a dry atmosphere of nitrogen unless otherwise specified. Indicated reaction temperatures refer to the reaction bath, while room temperature (rt) is noted as 25 °C. Commercial grade reagents and anhydrous solvents were used as received from vendors and no attempts were made to purify or dry these components further. Removal of solvents under reduced pressure was accomplished with a Buchi rotary evaporator at approximately 28 mm Hg pressure using a Teflon-linked KNF vacuum pump. The measurement of pH for neutralizations or acidifications was measured with Hydrion pH paper (MicroEssential Lab). Thin layer chromatography (TLC) was performed using 1” × 3” AnalTech No. 02521 silica gel plates with fluorescent indicator. Visualization of TLC plates was made by observation with either short wave UV light (254 nm lamp), 10% phosphomolybdic acid in ethanol, or in iodine vapours. Preparative thin layer chromatography was performed using Analtech, 20 × 20 cm, 1000-micron preparative TLC plates. Flash column chromatography was carried out using a Biotage Selekt Unit and a Biotage® Selekt System with Biotage Sfär silica gel columns. Proton NMR spectra were obtained on a 600 MHz Bruker AV III nuclear magnetic resonance spectrometer. Chemical shifts (δ) are reported in parts per million (ppm) and coupling constant (J) values are given in Hz, with the following spectral pattern designations: s, singlet; d, doublet; t, triplet, q, quartet; quint, quintet; m, multiplet; dd, doublet of doublets; dt, doublet of triplets; dq; doublet of quartets; br, broad signal. Tetramethylsilane was used as an internal reference. Peak listing, multiplicity designations, and coupling constant calculations were conducted using Mnova v.14 software (Mestrelab Research). Carbon NMR spectra were obtained on a 600 MHz Bruker AV III nuclear magnetic resonance spectrometer and tetramethylsilane was used as an internal reference. Any melting points provided are uncorrected and were obtained using a Stanford Research Systems OptiMelt melting point apparatus (MPA100) with an automated melting point system. Mass spectroscopic (MS) analyses were performed using ESI ionization on a Shimadzu LC-MS 2020 single quadrupole mass spectrometer with a binary solvent system A and B using a gradient elusion [A, H2O with 0.01% trifluoroacetic acid; B, CH3CN with 0.01% trifluoroacetic acid] and flow rate = 0.5 mL/min). A Shimadzu Nexcol C18 5µM, 50 × 3.0 mm was used. High pressure liquid chromatography (HPLC) purity analysis was performed using a Shimadzu LC-2050C with a binary solvent system A and B using a gradient elusion [A, H2O with 0.01% trifluoroacetic acid; B, CH3CN with 0.01% trifluoroacetic acid] and flow rate = 0.5 mL/min, with UV detection at 254 nm (system equipped with a photodiode array (PDA) detector). An X Bridge C18 5µm 4.6 × 150 mm was used. High resolution mass spectrometry (HRMS) analysis was performed using a Thermo Scientific Orbitrap Eclipse Tribrid mass spectrometer. Structural confirmation of RPI-GLYT2-82 was determined via $^{1}$H NMR, $^{13}$C NMR, and HRMS and the purity was assessed to be ≥95% via $^{1}$H NMR, $^{13}$C NMR, and HPLC.

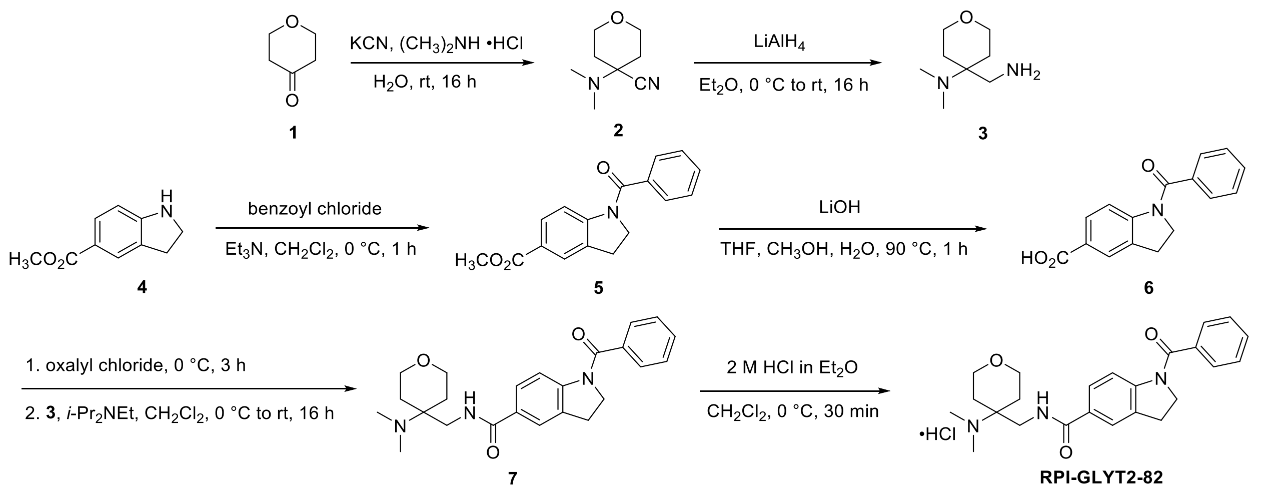

**Exhibit 1. Synthesis overview of compound RPI-GLYT2-82**

*1-Benzoyl-N-((4-(dimethylamino)tetrahydro-2H-pyran-4-yl)methyl)indoline-5-carboxamide Hydrochloride (**RPI-GLYT2-82**)*. Step A: To a solution of tetrahydro-4H-pyran-4-one (**1**, 10.0 g, 100 mmol) in $H_2O$ (500 mL) was added *N,N*-dimethylamine hydrochloride (8.15 g, 100 mmol) followed by KCN (6.50 g, 100 mmol). The reaction mixture was stirred at rt for 16 h with completion of the reaction determined by TLC. The reaction mixture was extracted with $Et_2O$ (3 × 500 mL), and the combined organic layers were washed sequentially with $H_2O$ (2 × 500 mL) and saturated brine solution (200 mL). The organic phase was dried over anhydrous $Na_2SO_4$, filtered, and the solvent was removed under reduced pressure to afford 4-(dimethylamino)tetrahydro-2H-pyran-4-carbonitrile (**2**) as a colorless oil (10.0 g, 65% yield). Amino nitrile **2** was used in the next step without further purification. $^1$H NMR (600 MHz, $CD_3OD$) δ 3.9-4.0 (m, 2 H), 3.58-3.62 (m, 2 H), 2.35 (s, 6 H), 2.11-2.15 (m, 2 H), 1.68–1.73 (m, 2 H).

Step B: To a 0 °C cooled solution of 4-(dimethylamino)tetrahydro-2H-pyran-4-carbonitrile (**2**, 5.0 g, 32.46 mmol) in anhydrous $Et_2O$ (250 mL) stirring under an $N_2$ atmosphere was slowly added $LiAlH_4$ (2.5 g, 64.93 mmol). The reaction mixture continued to stir for 16 h while gradually warming to rt. After confirming complete consumption of the starting material by TLC, the reaction mixture was diluted with additional $Et_2O$ (250 mL) and cooled to 0 °C. Dropwise addition of $H_2O$ (2.5 mL) was conducted slowly so as to maintain the temperature at 0 °C, followed by sequential dropwise addition of 15% aqueous NaOH solution (2.5 mL) and additional $H_2O$ (7.5 mL). The reaction mixture was allowed to warm to rt over 1 h followed by addition of $MgSO_4$ (10 g). The resulting mixture was allowed to stir at rt for an additional 30 min. The resulting precipitate was filtered, and the filtrate was concentrated under reduced pressure to yield 4-(aminomethyl)-N,N-dimethyltetrahydro-2H-pyran-4-amine (**3**) as a colorless oil (4.0 g, 80% yield). Amine **3** was used as is in the next step without further purification. $^1$H NMR (600 MHz, $CD_3OD$) δ 3.82-3.85 (m, 2 H), 3.55-3.59 (m, 2 H), 2.86 (s, 2 H), 2.32 (s, 6 H), 1.78-1.83 (m, 2 H), 1.53–1.56 (m, 2 H); $^{13}$C NMR (600 MHz, DMSO-$d_6$) δ 63.4, 55.8, 42.5, 37.7, 30.1; ESI MS (*m/z*): 159 $[M + H]^+$.

Step C: To a 0 °C cooled solution of methyl indoline-5-carboxylate (**4**, 2.00 g, 11.2 mmol) and $Et_3N$ (5.10 mL, 36.9 mmol) in $CH_2Cl_2$ (50 mL) was added benzoyl chloride (1.43 mL, 12.3 mmol) dropwise. The reaction mixture stirred for 16 h under an atmosphere of $N_2$ while gradually warming to rt. Completion of the reaction was confirmed by TLC. The mixture was diluted with

$CH_2Cl_2$ (200 mL) and washed sequentially with saturated aqueous $NH_4Cl$ solution (50 mL), saturated aqueous $NaHCO_3$ solution (50 mL), and saturated brine solution (50 mL). The organic layer was dried over $Na_2SO_4$, filtered, and the solvent was removed under reduced pressure. The resulting crude product was purified by flash column chromatography (10–30% EtOAc in hexanes) to afford methyl 1-benzoylindoline-5-carboxylate (**5**) as a white powder (3.10 g, 98% yield). $^1$H NMR (600 MHz, $CD_3OD$) δ 7.9 (bs, 1 H), 7.85 (bs, 1 H), 7.59-761 (m, 2 H), 7.56-7.57 (m, 2 H), 7.52-7.54 (m, 2 H), 4.14 (t, *J* = 8.2 Hz, 2H), 3.89 (s, 3H), 3.19 (t, *J* = 8.2 Hz, 2H); ESI MS (*m/z*): 282 $[M + H]^+$.

Step D: To a solution of methyl 1-benzoylindoline-5-carboxylate (**5**, 3.10 g, 11.03 mmol) in a 3:2:1 mixture of THF, $CH_3OH$, and $H_2O$ (50 mL) was added $LiOH{\cdot}H_2O$ (0.397 g, 16.5 mmol) and the mixture was heated at 90 °C for 1 h. Completion of the reaction was confirmed by TLC. THF and $CH_3OH$ were evaporated under reduced pressure, and the resulting aqueous mixture was further diluted with $H_2O$ (100 mL). The aqueous mixture was extracted with EtOAc (2 × 100 mL) and then acidified to pH ~2 with 5 N HCl. The resulting precipitate was collected by filtration, washed with $H_2O$, and dried under vacuum to afford 1-benzoylindoline-5-carboxylic acid (**6**) as an off-white solid (2.50 g, 86% yield). $^1$H NMR (600 MHz, DMSO-$d_6$) δ 12.7 (bs, 1 H), 7.94-7.96 (m, 1 H), 7.80-7.82 (m, 2 H), 7.60-7.64 (m, 2 H), 7.49-7.56 (m, 3H), 4.05 (t, *J* = 8.4 Hz, 2H), 3.13 (t, *J* = 8.3 Hz, 2H); $^{13}$C NMR (600 MHz, DMSO-$d_6$) δ 169.0, 167.4, 146.9, 137.0, 133.6, 133.2, 129.7, 129.6, 128.9, 127.3, 126.5, 126.3, 116.2, 51.2, 27.7; ESI MS (*m/z*): 268 $[M + H]^+$.

Step E: To a 0 °C cooled solution of 1-benzoylindoline-5-carboxylic acid (**6**, 1.0 g, 3.74 mmol) in $CH_2Cl_2$ (25 mL) was added oxalyl chloride (0.64 mL, 7.48 mmol) dropwise under an atmosphere of $N_2$. A catalytic amount of DMF (0.5 mL) was added, and the mixture stirred for 3 h while gradually warming to rt. The solvent and excess oxalyl chloride were removed under reduced pressure, yielding the crude acid chloride intermediate as a light yellow solid. Separately, a 0 °C cooled solution of 4-(aminomethyl)-*N*,*N*-dimethyltetrahydro-2H-pyran-4-amine (**3**, 0.88 g, 5.61 mmol) and *i*-$Pr_2NEt$ (3.4 mL, 18.7 mmol) in $CH_2Cl_2$ (15 mL) was prepared. To this solution was slowly added a solution of the crude acid chloride intermediate dissolved in $CH_2Cl_2$ (15 mL) under an atmosphere of $N_2$. The resulting reaction mixture stirred for 16 h while gradually warming to rt. After confirming complete consumption of the starting materials by TLC, the reaction mixture was diluted with $CH_2Cl_2$ (200 mL) and sequentially washed with saturated aqueous $NaHCO_3$ solution (50 mL), saturated aqueous $NH_4Cl$ solution (50 mL), and saturated brine solution (50 mL). The organic layer was dried over anhydrous $Na_2SO_4$, filtered, and the solvent was evaporated under reduced pressure. The crude product was purified by flash column chromatography (5–10% $CH_3OH$ in $CH_2Cl_2$) to yield 1-benzoyl-N-((4-(dimethylamino)tetrahydro-2H-pyran-4-yl)methyl)indoline-5-carboxamide (**7**) as a white solid (1.3 g, 86% yield). $^1$H NMR (600 MHz, $CDCl_3$) δ 7.7 (bs, 1 H), 7.57 (d, *J* = 7.6 Hz, 2H), 7.51-7.53 (m, 2 H), 7.47-7.49 (m, 2 H), 6.86 (bs, 1H), 4.15 (bs, 1H), 3.89 (d, *J* = 11.8 Hz, 2H), 3.68 (d, *J* = 4.6 Hz, 2H), 3.64 (t, *J* = 11.3 Hz, 2H), 3.64 (t, *J* = 11.3 Hz, 2H), 3.18 (t, *J* = 8.34 Hz, 2H), 2.33 (s, 6H), 1.89-1.94(m, 2H), 1.41 (d, *J*= 4.6 Hz, 2H); ESI MS (*m/z*): 408 $[M + H]^+$.

Step F: To a 0 °C cooled solution of 1-benzoyl-N-((4-(dimethylamino)tetrahydro-2H-pyran-4-yl)methyl)indoline-5-carboxamide (**7**, 1.3 g, 3.19 mmol) in $CH_2Cl_2$ (20 mL) was added a 2.0 M solution of HCl in $Et_2O$ (3.2 mL, 6.38 mmol). The reaction mixture was stirred at 0 °C under an atmosphere of $N_2$ for 30 min, and the solvent was evaporated under reduced pressure to yield an off-white solid, which was triturated with hexanes and subsequently lyophilized from $CH_3CN$ and $H_2O$ to yield 1-benzoyl-N-((4-(dimethylamino)tetrahydro-2H-pyran-4-yl)methyl)indoline-5-carboxamide hydrochloride (**RPI-GLYT2-82**) as a white powder (1.3 g, 93% yield). $^1$H NMR

(600 MHz, DMSO-$d_6$) δ 10.2 (bs, 1H), 8.66 (bs, 1H), 7.85 (s, 1H), 7.8 (bs, 1H), 7.6-7.61 (m, 2H), 7.50-7.55 (m, 3H), 4.05 (t, $J$ = 8.2 Hz, 2H), 3.85-3.89 (m, 4H), 3.63-3.67(m, 2H) 3.14 (t, $J$ = 8.3 Hz, 2H), 2.78 (d, $J$ = 8.3 Hz, 6H), 1.91 (m, 4H); $^{13}$C NMR (600 MHz, DMSO-$d_6$) δ 168.9, 167.7, 145.9, 137.1, 133.5, 130.8, 129.3, 129, 127.7, 127.4, 125, 64.6, 63.7, 37.2, 37.9, 36.9, 30.1, 27.9; HRMS (m/z): calc for $C_{24}H_{30}N_3O_3$ $[M+H]^+$: 408.2287 obtained 408.2282; HPLC 99.2% (AUC), $t_R$ = 14.6 min.

**Exhibit 2. $^{1}$H NMR Data for RPI-GLYT2-82**

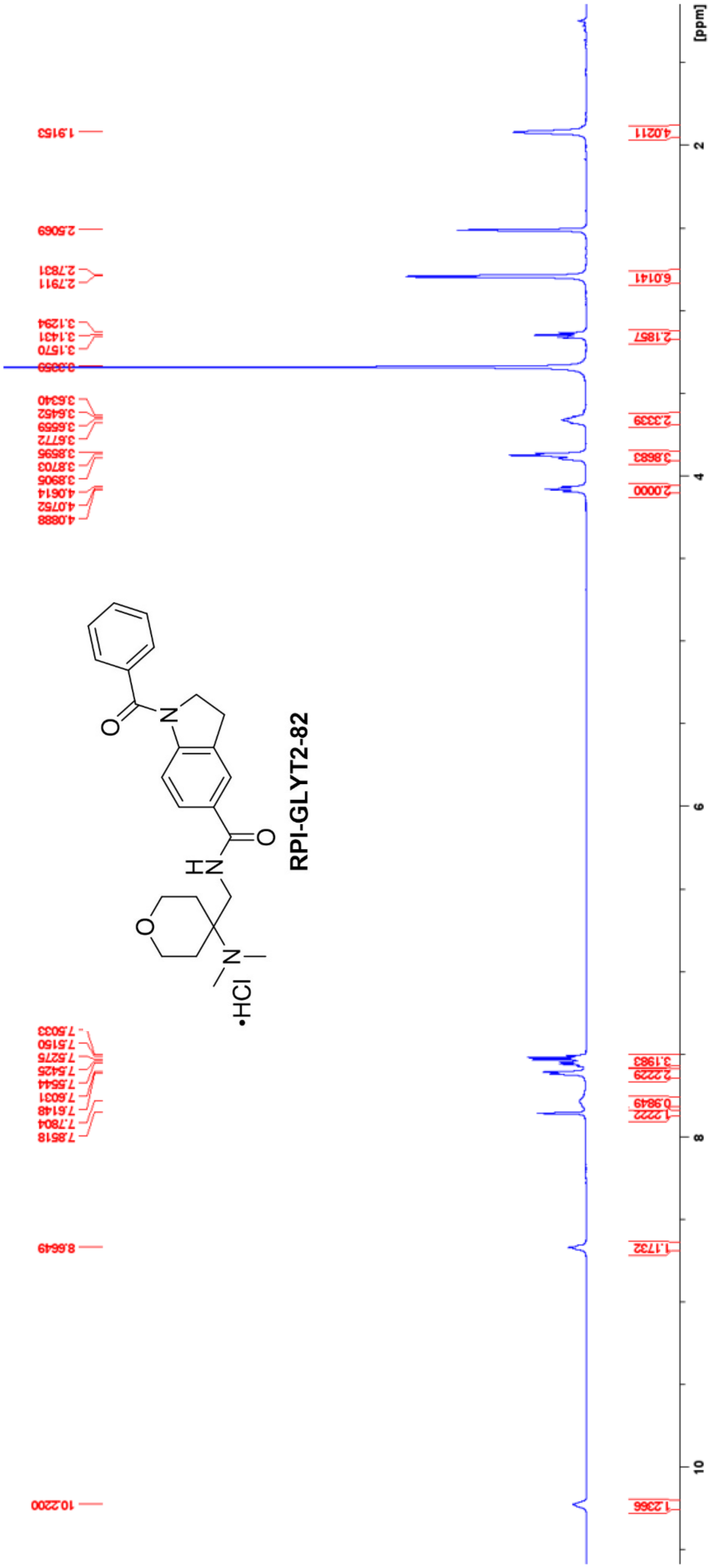

**Exhibit 3. $^{13}C$ NMR Data RPI-GLYT2-82**

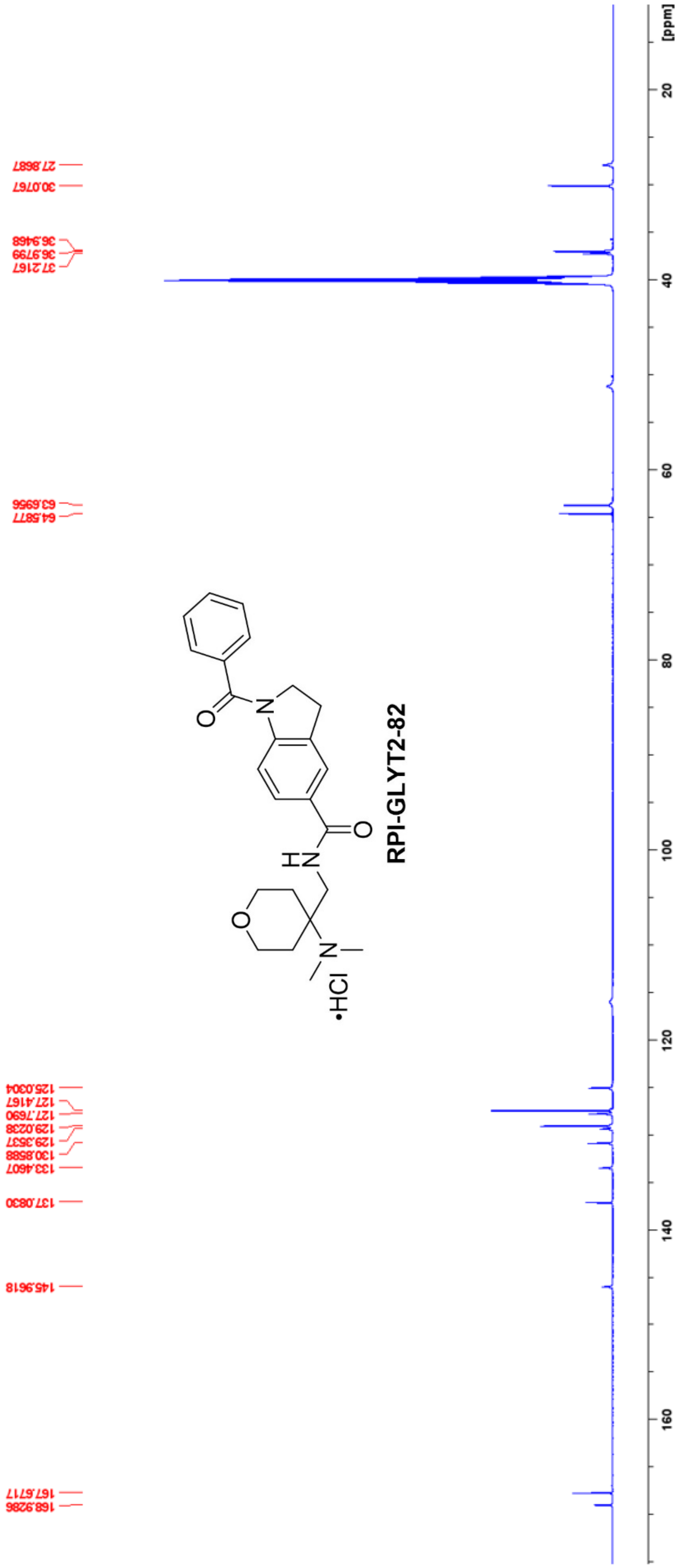

#### Exhibit 4. HPLC for RPI-GLYT2-82

Mobile Phase
Aqueous Reservoir (A): 0.01% TFA in H2O
Organic Reservoir (B): 0.01% TFA in CH3CN
Flow Rate: 0.3 mL/min
Injection Volume: 20 µL
Wavelength: 254 nm

HPLC Gradient

| Time (min) | %A | %B |
|---|---|---|
| 0.01 | 95 | 05 |
| 3.00 | 95 | 05 |
| 13.00 | 05 | 95 |
| 18.00 | 05 | 95 |
| 30.00 | Stop | |

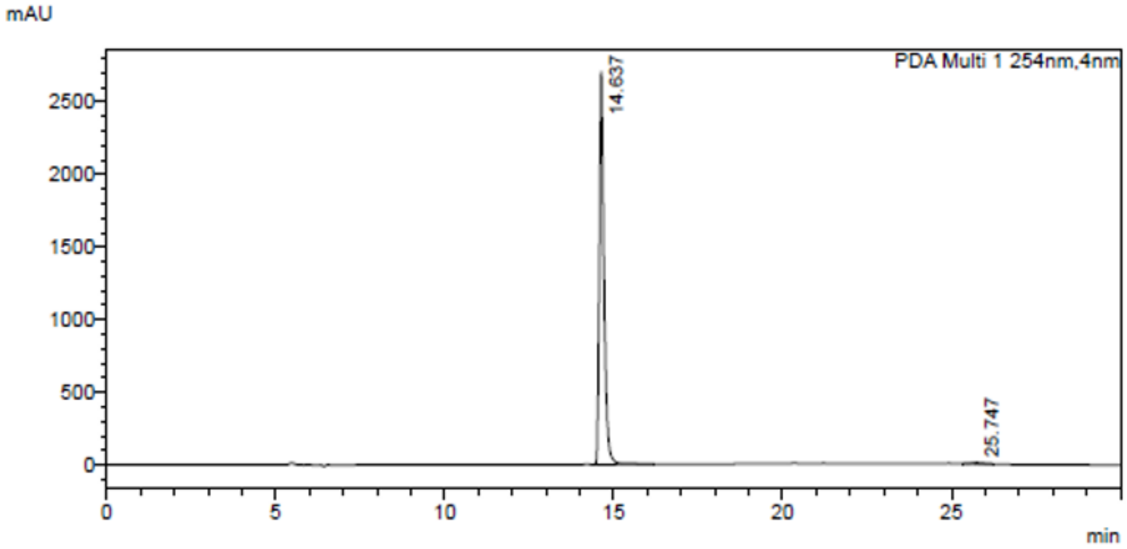

PDA Ch1 254nm

| Peak# | Ret. Time | Height | Area | Area% |
|---|---|---|---|---|
| 1 | 14.637 | 2705233 | 25898085 | 99.257 |
| 2 | 25.747 | 8182 | 193896 | 0.743 |
| Total | | 2713415 | 26091981 | 100.000 |

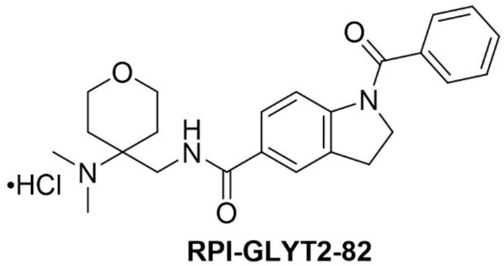

**RPI-GLYT2-82**

### Exhibit 5. HRMS Data for RPI-GLYT2-82

D:\Users\...\20241210Chris\Chris-RPI-N82 12/10/24 13:47:34

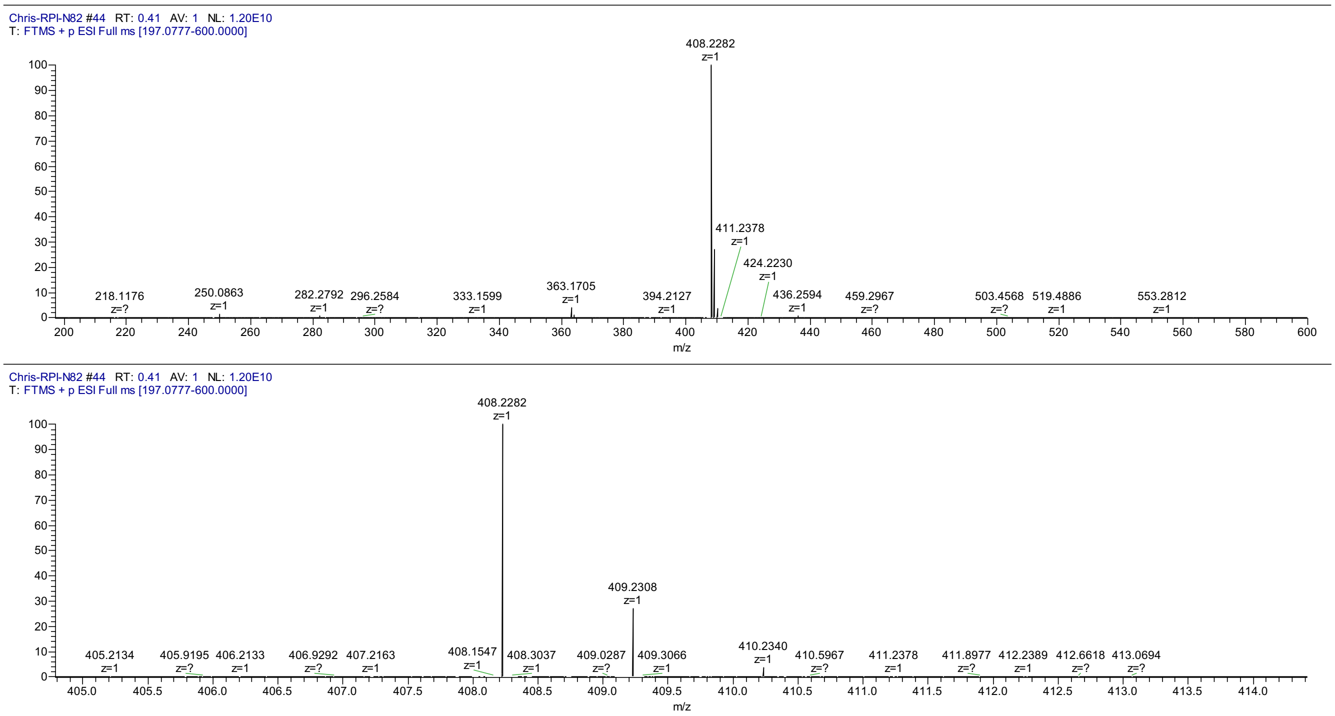

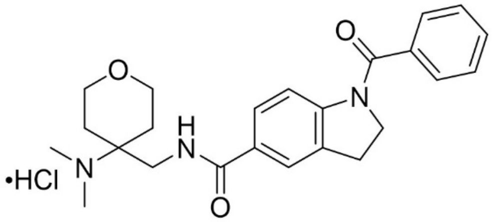

**RPI-GLYT2-82**
